# An agent-based approach for designing effective protection

**DOI:** 10.64898/2026.04.03.716393

**Authors:** Elisabeth Slooten, L. Scott Myers, Jacob Nabe-Nielsen

## Abstract

We developed an agent-based model (ABM) to assess how area-based controls on fishing methods can reduce fishing mortality and population declines. The model incorporates the behavior and distributions of dolphins and fishing vessels, and realistic displacement of fishing effort when protection is extended. Our case study is New Zealand dolphin – Hector’s and Maui dolphins. The model was designed and calibrated using pattern-oriented modeling. Our results show that mortality due to entanglement in fishing gears has been reduced thanks to a gradual increase in dolphin protection. However, current protection is not as effective as previously thought, and scarce populations are negatively affected by Allee effects. Neither national nor international goals for reducing bycatch are met by current dolphin protection. The IUCN has recommended banning gillnet and trawl fisheries in New Zealand waters < 100m deep. For most New Zealand dolphin populations, this would be effective in achieving national and international goals for reducing bycatch. Only two populations would require additional protection. This modelling approach is also suitable for assessing impacts of bycatch and ship strikes for other marine species, making it suitable for informing management decisions in many regions.

## Introduction

Most human impacts on marine mammals, seabirds and other threatened marine species have a strong spatial aspect, and quantifying the population-level consequences of these impacts is important for conservation of the affected species (Wiegand et al. 2002; Halpern et al. 2008, 2019). For example, vessel sounds that induce behavioral changes in nearby harbor porpoises (Frankish et al. 2023) and establishment of offshore wind farms that result in local displacement of birds (Lamb et al. 2024) have larger impacts when they overlap with important foraging areas. Likewise, the severity of direct (bycatch) and indirect (food depletion) impacts of fishing on marine mammals depends on the extent of overlap between fishing and the impacted population (van Beest et al. 2017, Burgess et al. 2018). To study the long-term consequences of such human impacts it is essential to consider the distribution and movements of the affected animals, which determines the proportion of the population that gets impacted, the animals’ ability to recolonize suitable habitats and their ability to find mates. These aspects of population dynamics are rarely considered when managing marine species.

The combined population impacts of different pressures that vary in time and space, such as bycatch and Allee effects, can be studied with agent-based models (ABMs) that incorporate spatial structure and realistic animal movements (e.g. van Beest et al. 2017; Nabe-Nielsen et al. 2018; de Jager et al. 2019; Railsback and Grimm 2019). In ABMs the movements and survival of individual animals can be modelled explicitly in realistic landscapes, allowing population dynamics to emerge from the same mechanisms as they do in nature (Grimm and Railsback 2005; Stillman et al. 2015). By incorporating both dolphins and fishing vessels, our approach allows for an assessment of the efficiency of different conservation options.

Fisheries mortality is the most serious threat to marine mammals and many other endangered marine species (Lewison et al. 2004; Read et al. 2006; Avilla et al. 2018; Brownell et al. 2019; United Nations Office of Legal Affairs 2021; Elliott et al. 2023; Lusseau et al. 2023; Authier 2024). Bycatch in fishing gear has contributed to substantial declines of many marine mammal populations, and in the extinction of at least one species (Wade et al. 2021) and is driving current declines in vaquita (Jaramillo-Legorreta et al. 2019), Atlantic humpback dolphin (Weir et al. 2011) and other endangered cetaceans (Brownell et al. 2019; Ballance et al. 2021). Haider et al. (2017) recommend that management of marine mammal bycatch should also consider Allee effects caused by small population size or density limiting population growth and suggest that sustainable levels of bycatch for populations experiencing an Allee effect should be between half and two-third that of other populations (Haider et al. 2017).

New Zealand dolphin *(Cephalorhynchus hectori)*, an endangered dolphin species, is impacted by fisheries mortality in gillnet and trawl fisheries. Our research provides much needed information on the effectiveness of dolphin protection measures, consisting of protected areas where gillnets, trawling or both are prohibited. We estimate the effectiveness of current protection in reducing bycatch below national and international thresholds, compared to past protection and protection throughout waters < 100 m deep, as recommended by the International Union for the Conservation of Nature (IUCN 2012; Figure 1).

**Figure 1.**
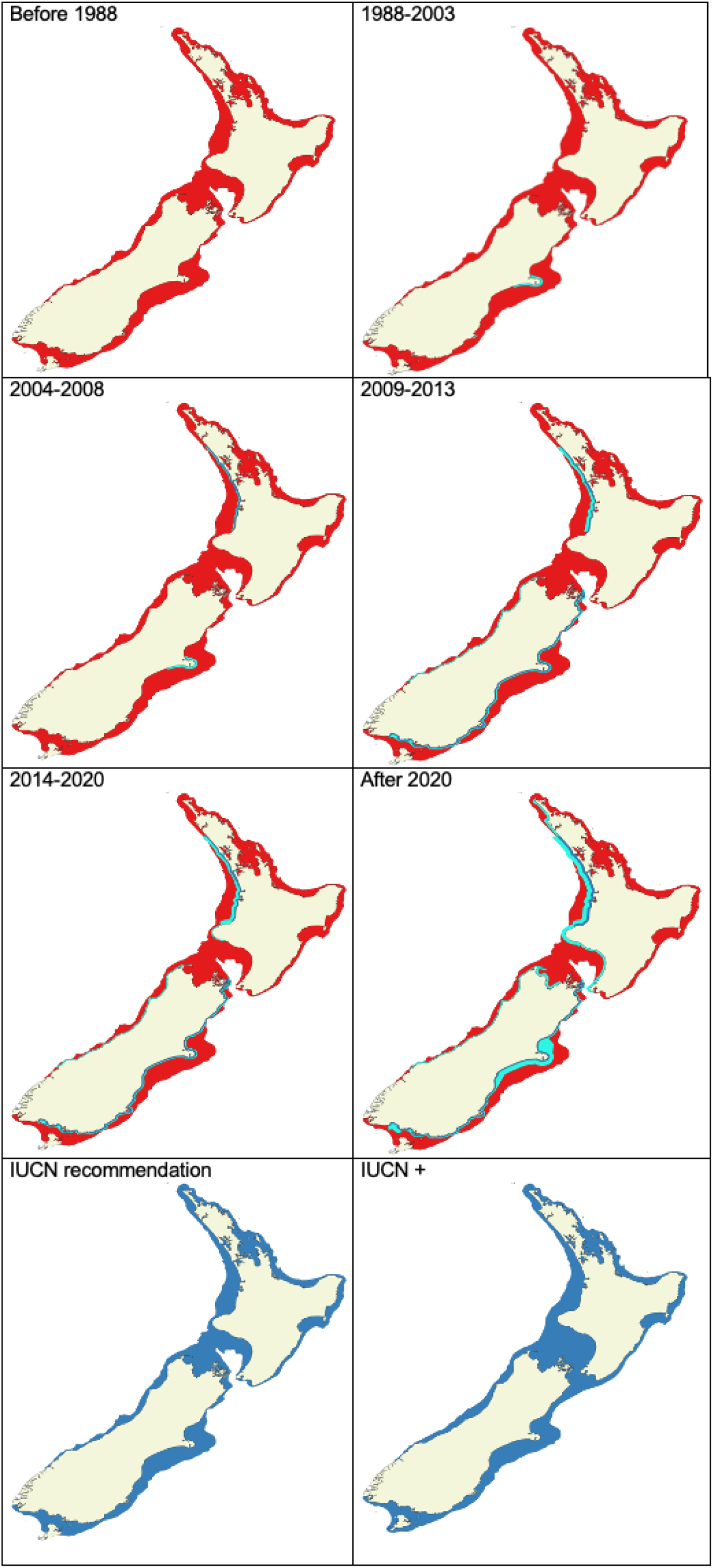
History of dolphin protection, with the distribution of Hector’s and Māui dolphin indicated in red, areas where gillnets are banned in light blue and areas where both gillnets and trawling are banned in dark blue.

Fisheries mortality ; has caused declines to approximately 30% of original population size around the South Island and to below 5% around the North Island (Slooten 2007, 2013, 2020; Slooten and Dawson 2010, 2021; Slooten and Davies 2012; Slooten et al. 2000; Cooke et al. 2019). Survival rates in the best studied population increased from 86% when there was no protection from fisheries bycatch to 91% with partial protection (Gormley et al. 2012; Wickman 2024), reducing population declines and highlighting the need to prevent bycatch in certain areas. The dolphins are caught in two different types of fishing nets: gillnets and trawls. Gillnets are stationary in the water and made of very thin nylon mesh, which can trap dolphins. Trawling vessels drag large, heavy nets either along the seafloor or through mid-water. Dolphins are attracted to trawlers, repeatedly diving down to the net to take fish. Frequently large groups of dolphins follow trawlers (Rayment and Webster 2009). Camera monitoring of trawlers over the last few years indicates that bycatch in trawl nets is more common than previously thought (MPI 2025a,b).

We built an ABM of dolphins, gillnets, and trawlers in the coastal waters around New Zealand to make realistic assessments of how the dolphins are affected by fisheries mortality. We use the model to assess the effectiveness of current protection measures (Figure 1) compared to past protection and protection measures proposed by the International Union for the Conservation of Nature (IUCN 2012). Our model fills a gap in the tools that can be used for assessing whether conservation measures are sufficient to prevent future population decline in different populations. It could be adapted for other species and other human impacts. In this paper we address the following questions: how effective is current dolphin protection in reducing dolphin bycatch, is current bycatch below national and international thresholds, and how many dolphins are caught each year in gillnet and trawl fisheries under various protection levels. We conclude that the level of protection recommended by the IUCN is effective for most populations. However, two populations require a higher level of protection, which we call IUCN+. These are the populations around the North Island and north-east South Island of New Zealand.

## Materials and Methods

Our analysis incorporates data on the spatial and temporal distribution of dolphins, gillnets and trawling vessels, dolphin life history characteristics and movements of dolphins and trawlers. Bycatch rates are calibrated based on data from fisheries observers and camera monitoring. Below we describe these data, and subsequently we describe how they are used for constructing and calibrating the ABM.

### 1. Study species

New Zealand dolphin *(Cephalorhynchus hectori)* is found only in New Zealand waters, with a northern (Māui dolphin) and southern sub-species (Hector’s dolphin). Māui dolphin is Critically Endangered (Constantine 2023). Several populations of Hector’s dolphin are also very small and vulnerable. For example, there are an estimated 41-42 Hector’s dolphins off Otago (Turek et al. 2013; Williams 2022) and in other areas the species appears to have been extirpated altogether (McGrath 2020).

Dolphin distribution and abundance data are available from population surveys (Dawson and Slooten 1988; Dawson et al. 2004; Slooten et al. 2004, 2006; Rayment et al. 2009, 2010; Turek et al. 2013; MacKenzie and Clement 2014, 2016, 2019; Constantine et al. 2021; IWC 2023). In areas with high dolphin densities off the east and west coast of the South Island most sightings are in waters < 100m deep with a small proportion (<5%) of sightings in deeper water. In low density areas, data from small boat surveys and acoustic surveys show the same pattern of dolphin distribution with respect to water depth and distance from shore (e.g. Turek et al. 2013; Harvey 2021; Williams 2022; Bennington 2024).

Estimates of group size, movements, survival and reproductive rates are available from 40 years of monitoring New Zealand dolphins (e.g. Slooten 1991; Slooten and Lad 1991; Gormley 2009; Wickman 2024; Bennington 2025). Females give birth to their first calf at age 7–9 years and have one calf every 2–4 years. The most common group size is 2–10 individuals. New Zealand dolphins are attracted to trawlers (Rayment and Webster 2009), with large numbers of dolphins observed following trawlers (Figure A.1). Dolphins typically move 1–2km per hour, with occasional movements up to 7km per hour (Figure B.24). Dolphin groups occasionally travel in one direction for half an hour or so, but rarely travel in the same direction for longer than that.

Home ranges are on average 50km of coastline, with a maximum distance travelled of 107km (Rayment et al. 2009). Individual dolphins are consistently seen year-round in the same geographical area (Bräger et al. 2002; Rayment et al. 2009; Bräger and Bräger 2018), with <1% of the animals moving among different populations >100km apart per year (Fletcher et al. 2002). When male cetaceans search for sexually active females the fertilization rate of females depends in part on the amount of time needed for a male to travel between groups relative to the length of time each female is fertile and the degree of synchrony among females (e.g. Whitehead 1987, Whitehead and Weilgart 2000). Such mating systems can result in Allee effects, with relatively lower fertilization rates in low-density areas (Slooten 1991).

### 2. Fisheries and bycatch data

Fisheries data were obtained from the Ministry for Primary Industries (MPI 2023, 2024, 2025a,b), Global Fishing Watch (GFW 2025) and other publicly available sources (e.g. Dragonfly 2025), including fishing locations, numbers of fishing vessels per port, number of trawls and length of each trawl. Government observers and camera monitoring programs provided information on the probability of finding an entangled Hector’s or Maui dolphin when hauling a gillnet or trawl net (Baird and Bradford 2000; Starr and Langley 2000; DOC 2025; MPI 2025a,b). The Banks Peninsula area (Figure 2) has relatively high dolphin density as well as observer and camera coverage (MPI 2025a,b) and therefore the most robust data on dolphin bycatch.

**Figure 2.**
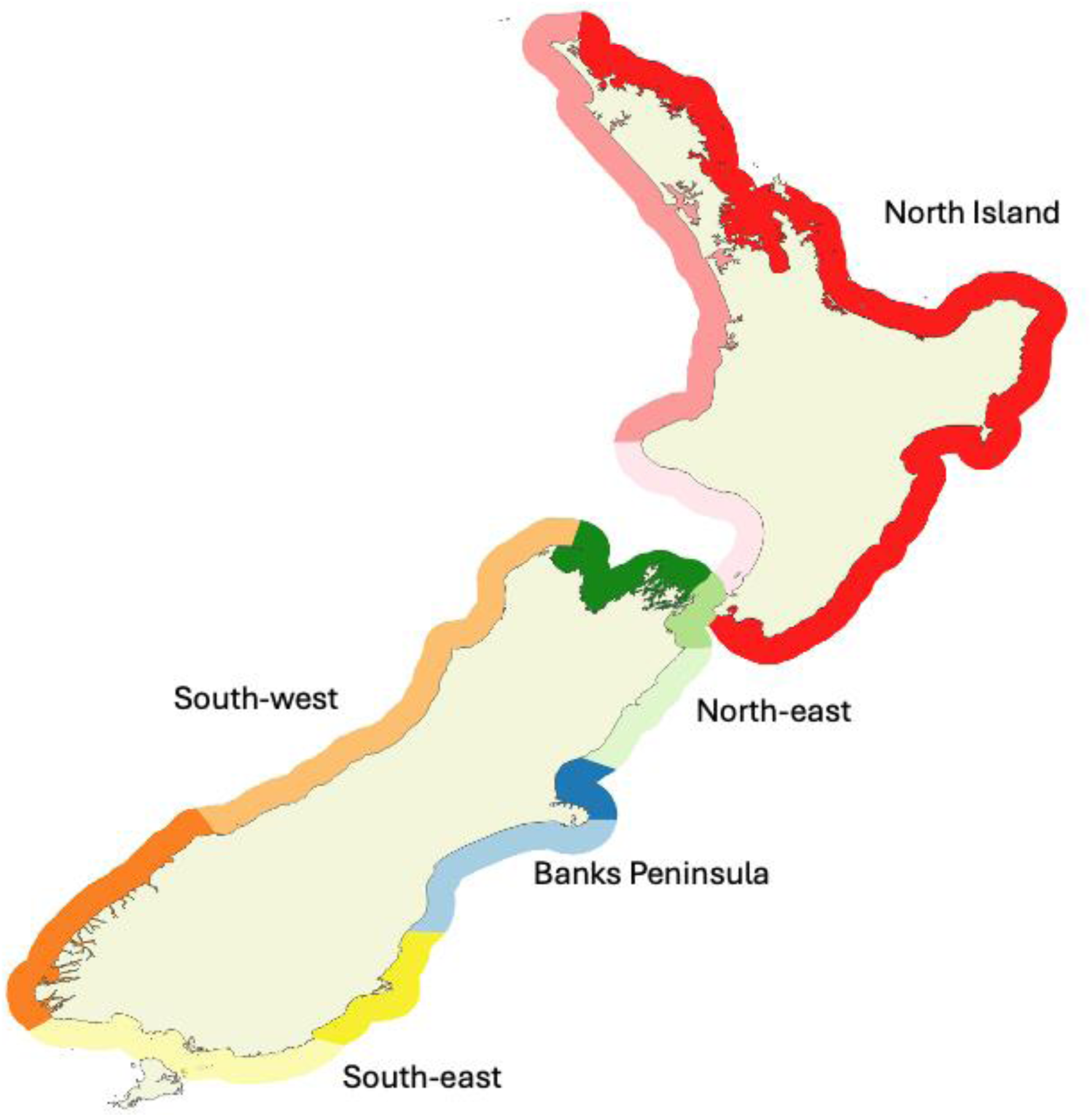
Map of the five modelled management areas with different hues of each colour showing the placement areas where dolphins, gillnets and trawlers are placed at the start of each simulation (e.g. Banks Peninsula north in dark blue and Banks Peninsula south in light blue).

### 3. Model description

We developed an ABM to study how bycatch of New Zealand dolphins is influenced by different levels of protection. The following section describes the model using the ODD (Overview, Design concepts, and Details) protocol (Grimm et al. 2020). The first three sections of the ODD are included below, with a full ODD in Section B of the Supporting Information. The model was developed using NetLogo (Wilensky 1999).

#### 3.1 Purposes and patterns

The purpose of our model is to predict the effects of gillnet and trawler by-catch deaths on the long-term population dynamics of Māui and Hector’s dolphins around New Zealand under various protection schemes and levels of fishing effort, incorporating realistic dolphin and fishing vessel movements and densities, and considering how reduced population densities affect the likelihood of mating.

To consider our model realistic enough for its purpose, we evaluated its success at reproducing the following patterns: 1. dolphin movements in their home range in response to water depth and distance from shore, 2. movement of dolphins attracted to trawlers, 3. dolphin group size variation, 4. trawler movements and distribution of trawl effort, 5. bycatch rates for gillnets and trawlers.

We used the pattern-oriented modelling approach (POM) (Wiegand et al. 2003; Grimm et al. 2005; Kramer-Schadt et al. 2007) to evaluate the model entities’ movements and confirm realistic behaviors during two phases of the modelling process: during development to parameterize the model and during evaluation to test if model outputs are realistic. The resulting overlap in space and time between dolphins, trawlers and gillnets, as well as bycatch rates and mortality patterns match independent estimates of these parameters (see Section 4, below).

#### 3.2 Entities, State Variables, and Scales

The model world is 1024 x 1024 patches, each representing 1.63 x 1.63km. The world thus covers ∼ 2.786 x 10^6^ km^2^. This area includes the entire land mass of the North and South Islands of New Zealand (∼269,000 km^2^) with the remainder representing the surrounding ocean (Figure A.2; NIWA 2016). There is no world-wrapping. One time step in the model represents 1 hour.

The model includes the following entities: dolphins, trawlers, gillnets, ports, waypoints (navigation reference points for trawlers near ports), landscape patches (land or ocean) and modelled management areas based on fisheries statistical areas and dolphin surveys (Figure 2). Dolphins are “super-agents”; each represents a group of individual dolphins of various ages and genders. They are characterized by their sub-species (Hector’s or Māui), group size, age, gender, estrus/pregnancy/post-partum status of females, velocity, traveling direction, home range, and location (Table B.1). They are also characterized by whether they move towards or flock around a trawler, or if they “wander” within their home range when trawlers are not nearby. Trawlers are characterized by their size and home port according to data provided by MPI (2023), by their location, speed, destination, activity (trawling, in port, etc.) and by where they fish, length of trawls, length of fishing trips, and time in port between fishing trips. Gillnets are characterized by location and soak time. Ports are characterized by name and geographic location. Waypoints are characterized by location and associated port. Table B.1 lists all entities and state variables.

#### 3.3 Process Overview and Scheduling

A top-level flowchart showing an overview of the model processes that run every time step (hour) is shown in Figure 3. The number of trawlers and gillnets in the simulation are adjusted on January 1. New protection schemes are introduced on October 1 at the start of the fishing year. At startup, dolphins are placed in a home range and distributed offshore (i.e. alongshore distribution and distance from shore) to match population survey data.

**Figure 3.**
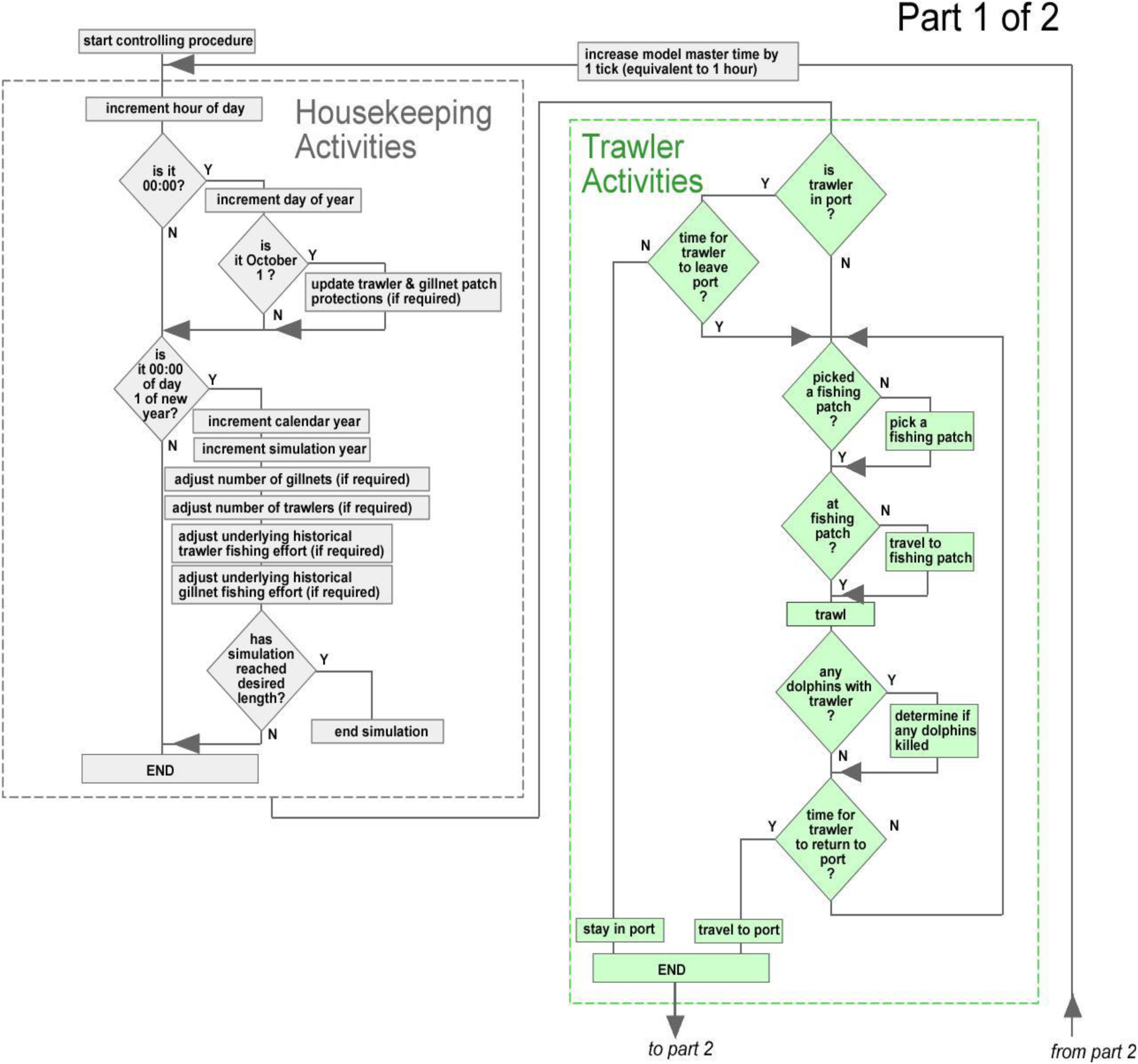
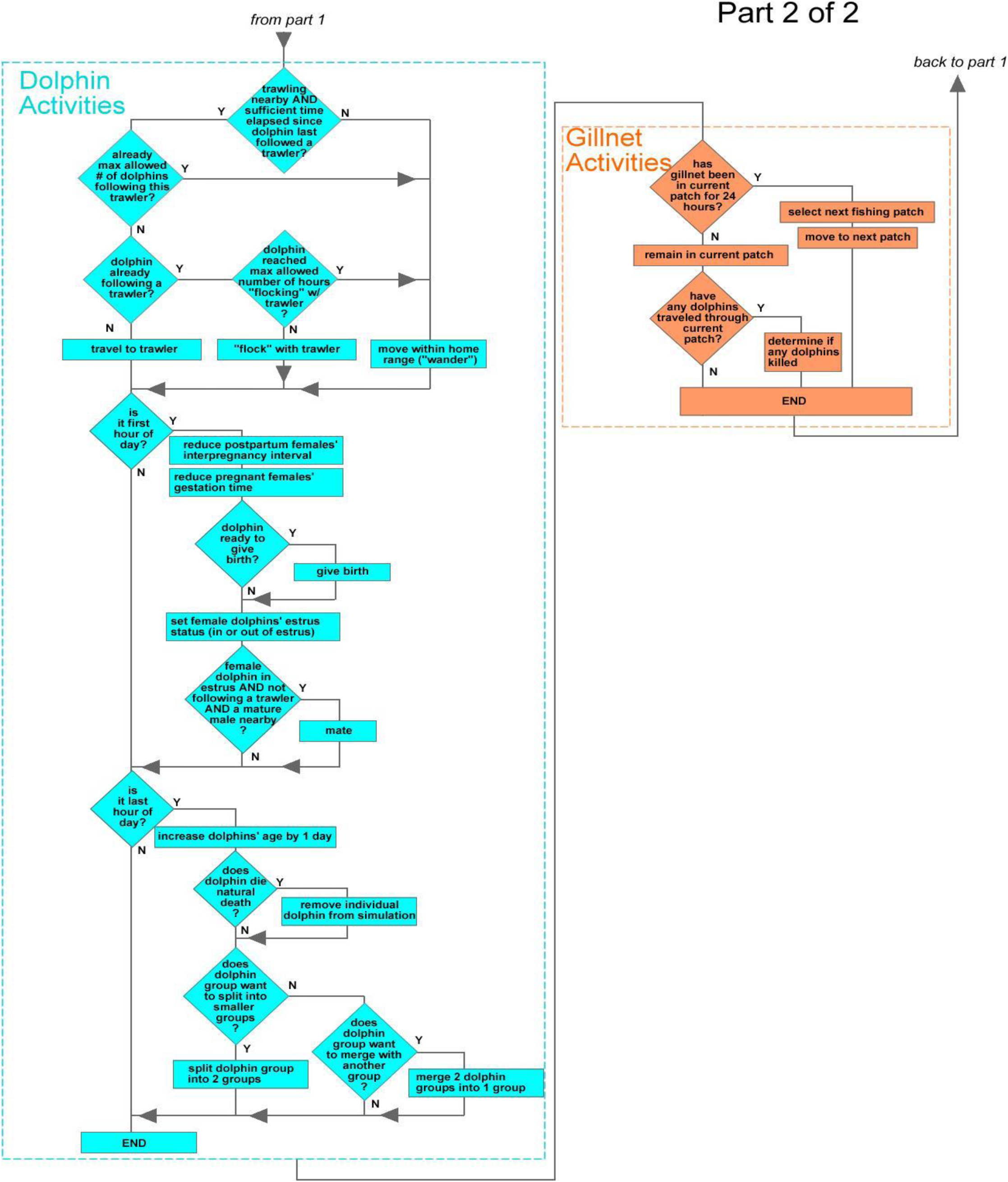
Main Controlling Procedure Flowchart: gray steps (top left) are run by the model as part of its ‘housekeeping’ obligations, green steps (top right) are run by trawlers, blue steps (bottom left) are executed by dolphins and orange steps (bottom right) are performed by gillnets.

Trawlers leave port when scheduled to do so (Figure 3), deploy their nets to fish (which attracts dolphins and can cause bycatch), and return to port at the end of their trip. Because trawlers take multi-hour fishing trips, no trawler performs every procedure in Figure 3 every hour. Each trawler has unique parameters for when it leaves port, trip duration, length of trawls, residence time in port between trips, velocity, etc. They remain in their home port until it is time to leave on a fishing trip. Then they select the duration of the fishing trip, velocity, and destination, with areas that have historically high fishing effort more likely to be selected.

Medium trawlers (16–25m long) take multi-day fishing trips and cannot fish in patches protected from trawling, while small trawlers (< 15m) only take day trips and can fish anywhere. They travel to their selected fishing spot and pick a trawl length (km) and trawling direction which tends to follow the shoreline or a depth contour. Average trawl speed is 2.2 knots for small and 4 knots for medium trawlers. Average travel speed is 8 knots for small and 9 knots for medium trawlers. As they trawl, they attract dolphins within detection range while their nets are in the water. The detection of trawlers by dolphins is determined by trawler noise (150dB for small and 171dB for medium trawlers; Daly and White, 2021) and the attenuation of trawler noise with distance (see Supporting Information section B for more detail). After each completed trawl, the model records any dolphin bycatch and the trawler evaluates whether it continues fishing or returns to port based on remaining trip length and projected travel time. Those that keep fishing can either stay in their current location and start a new trawl or select a different fishing area before they resume fishing. Trawlers ultimately return to port to conclude their trip and remain there until it is time to depart on their next trip (intervals between trips are based on historical data and are longer for medium trawlers). Because there can be multiple trawlers in a simulation, a single hour in the simulation could find some trawlers in port, while others are traveling to a fishing patch, fishing (potentially generating dolphin bycatch) or traveling home.

Dolphins decide on their next activity and velocity based on: whether there is trawling nearby, whether they detect it, whether they are already flocking with the trawler, or if not flocking, whether they choose to travel to it. “Flocking” dolphins travel adjacent (< 10m) to trawlers that are actually fishing, making them susceptible to death from bycatch. If not travelling to or flocking with a trawler, dolphins remain in their home range and “wander”, which is a catch-all term for time spent in non-trawler focused movement and behavior. Home ranges are individual to each dolphin and are assigned at model start-up based on field data for observed Hector’s dolphin home ranges. Dolphins “wander” via a vector-based random walk that keeps dolphins within their home range and within the maximum allowed water depth (Figure B.26). Each time a dolphin selects a new wandering heading it uses its current heading plus a random normal value with mean zero and a Standard Deviation (wiggle). This initial, unbiased heading is then modified based on the depth bias parameter k which controls the steepness of a logistic curve, defining the strength of the dolphins’ response to water depth. With high values of k, dolphins only respond when they are close to the maximum water depth (100m) and then respond strongly by changing their direction away from deep water. At low values of k, dolphins respond more gradually, turning away from deeper water when still in relatively shallow water and showing a relatively weaker response in terms of turning angle.

Dolphins reproduce on a mating/gestation/calving schedule that mimics field data and age-at-maturity constraints. The model includes an Allee effect which allows mature females to become pregnant only if there is a mature male within 5km. This causes the reproductive rate to be lower in when population densities are low. Field data from different New Zealand dolphin populations indicate that the proportion of calves is lower in smaller populations. The 5km range used in our model results in similar differences in the proportion of calves among populations of different sizes (Slooten 1991; Slooten and Lad 1991; Slooten et al. 1993; Turek et al. 2013; Harvey 2021; Constantine et al. 2021; Williams 2022; Bennington 2025). Dolphins adjust group size by merging into larger groups and splitting into smaller groups (“fission-fusion” behavior), ensuring that group sizes are as observed in the field (Figures C.5 and C.6). Their probability of dying from old age increases as animals approach the maximum age of 30 years.

Gillnets stay in the same patch for a 24-hour “soak time” where they might be encountered by dolphins, potentially resulting in bycatch. When soak time is complete, they relocate to another patch to soak for 24 hours.

##### Scheduling

The order of procedures is the same for each time step, however not all procedures run in every step; e.g., dolphin age is incremented only at the end of every day. The order in which specific dolphins, trawlers, and gillnets execute their steps is random for each time step, and their dynamic state variables are updated as soon as a new value is calculated.

### 4. Model calibration

Data on trawler behavior from MPI (2023, 2024) and GFW (2025) included length of fishing trips (in hours), length of each trawl (in km), number of days in port between fishing trips and total number of trawls. The following parameters were iteratively adjusted until the spatial distribution of fishing effort in the model approximated the government data: Maximum travel time between leaving port and the first trawl of the day, travel time between subsequent trawl events, minimum and maximum trawl start depth and the probability of each subsequent trawl being within the same fishing area or switching to a new fishing area (Figure 3). The fit between model fishing effort and government data on fishing effort was determined in R, using the *kde.points* function in the *GISTools* package to make kernel density estimates for simulated and observed positions of trawl starts (Figure C.1). Sums of squared differences (SSD) were used to determine which input parameters provided the best fit between simulated and observed positions of trawl starts. The same process was followed to ensure the spatial distribution of gillnets and dolphins matched data on the distribution of dolphins and fishing effort. For more information, please see Section C of the Supporting Information (Parameter estimation and pattern-oriented modelling).

We calibrated the model to ensure that dolphin movements and spatial distribution matched field data. To ensure that the dolphins responded realistically to water depth, the model was calibrated by generating heatmaps for a wide range of values of depth bias (k) and turning angle (wiggle). Results from the calibration runs were compared with dolphin density estimates from the three survey strata (0-4, 4-12 and 12-20nmi offshore, with the best model fit resulting from k = 15 and wiggle = 20 (Figure C.4). The probability of dolphins moving from one area to another was based on field data directly (Fletcher et al. 2002).

The probability of dolphin attraction to trawlers was calibrated to match field observations of groups of dolphins feeding close to trawl nets (Figure C.7). The probability of a dolphin dying when it is in the same location as a trawler or gillnet was estimated in the Banks Peninsula area (Figure 2). This is the only area with sufficient camera monitoring and observer coverage to yield robust estimates of dolphin deaths in these two fishing gears. We iteratively modified the probability of capture when a dolphin or group of dolphins encountered a gillnet or trawler until the catch rate per trawl or per 10km of gillnet matched the government estimates based on observers and camera monitoring (Figure 4).

**Figure 4.**
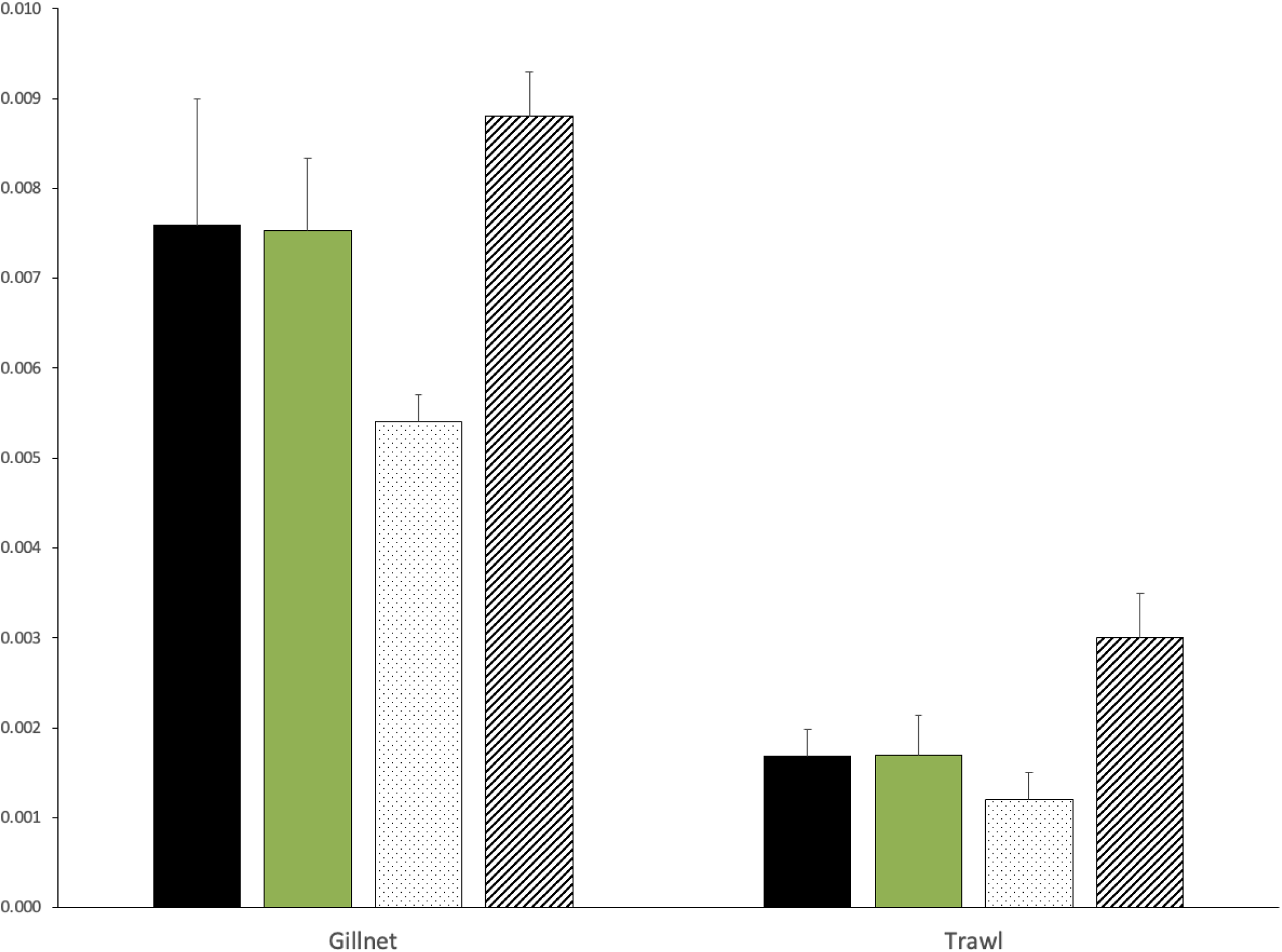
Catch rates (dolphin deaths per trawl and per 10km of gillnet) showing government estimates based on observer and camera monitoring, compared to the best model fit and two poor fits based on the risk of dolphin capture per encounter with a gillnet (0.003, 0.0052, 0.0055 respectively, for poor fit 1 shown in stipple, best fit and poor fit 2 shown in stripes) and per encounter with a trawl net (0.00013, 0.00014, 0.00015 respectively).

### 5. Scenarios

Dolphin protection has improved over time (Figure 1). To investigate the effects of changes in the protection level in 1988, 2003, 2008, 2013 and 2020 (Slooten and Dawson 2021) we updated the protection scheme in each of those years by preventing trawlers and gillnets from entering protected areas. We also explored two potential future protection schemes: the IUCN (2012) recommended using the 100m depth contour as the offshore boundary of protection and the option we have called IUCN+ which adds a small buffer around the 100m depth contour in places where depth contours are relatively steep and there is deep water close to shore.

We ran model scenarios with different population densities to test for the presence of an Allee effect. The model requires a mature male to be nearby before a female will become pregnant. We ran the model for the South-east population with population sizes of 10, 100 and 1000 individuals to estimate birth rates under different population density conditions.

## Results

Our model produced realistic emergent dolphin and trawler distributions after calibration and also reproduced bycatch rates from observer and camera monitoring (Figure C.1), making it suitable for assessing population effects of changes in protection.

The current level of protection (Figure 1) results in a decrease in the annual number of dolphins caught in all areas as compared to numbers caught before 2020 (Figure 5). Past protection was implemented in 2008 for South Island areas and 2013 for the North Island.

**Figure 5.**
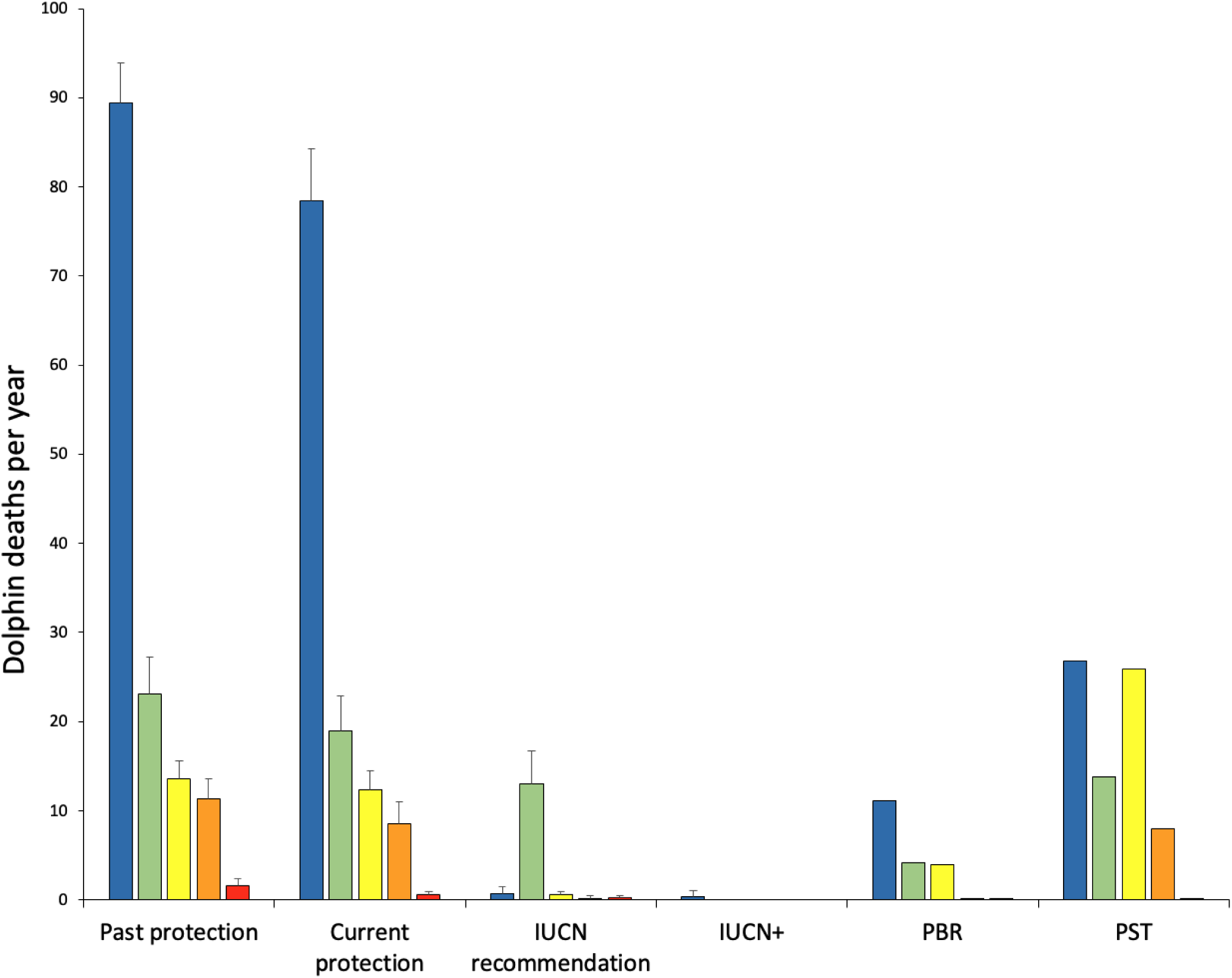
Modelled fisheries mortality for five different areas (area colours from Figure 2, from left to right: Banks Peninsula, North-east, South-east, South-west and North Island) and four different levels of protection (see Figure 1), with error bars showing 95% CI based on 10 model runs. These estimated levels of bycatch are compared with two different thresholds for total human impact: The PBR developed by the US National Fisheries Service and the PST developed by the New Zealand Ministry for Primary Industries. As these are maximum limits, or thresholds, they do not include confidence intervals.

Differences among populations reflect the effectiveness of different protection options in different areas. In 2020, Māui dolphin bycatch dropped from about 1.6 dolphins per year to about 0.6 (one dolphin death in fishing nets every 1.7 years). The protection recommended by the IUCN would further decrease bycatch, with the IUCN+ protection level eliminating bycatch. Likewise, the IUCN+ option dramatically reduced bycatch of Hector’s dolphins off the north-eastern South Island, where bycatch gradually declines when comparing 2008, 2020 and IUCN levels of protection, and the IUCN+ option eliminates bycatch (Figure 5).

The IUCN+ option provides much better protection in Cook Strait and around Kaikoura, where a deep ocean trench extends into inshore waters. In all other South Island areas, the IUCN option is very effective at reducing bycatch. Around south-eastern South Island, current protection has decreased bycatch compared to previous years, the IUCN option comes close to eliminating bycatch, and the IUCN+ option eliminates it. The west coast of the South Island has very little protection (Figure 1), with gillnets banned from the coastline to 2nmi offshore for 4 months of the year (November-February), but no protection from trawling. As expected, there is no appreciable decrease in bycatch between 2008 and 2020 (Figure 5). Implementing the IUCN option would dramatically decrease bycatch and the IUCN+ option would eliminate bycatch in this area.

In the Banks Peninsula area (Figure 2) the results indicate that bycatch was slightly lower in 2020 compared to 2008 (Figure 5). The area where gillnets are banned was extended substantially in 2020, but there has been no improvement in dolphin protection from trawl fisheries. Shallow waters extend well offshore in this area, and Hector’s dolphins range much further offshore than elsewhere around New Zealand, with gillnet and trawl fisheries still occurring in a large portion of Hector’s dolphin habitat. The area south of Banks Peninsula (shown in light blue in Figure 2) has a large Hector’s dolphin population that extends about three times as far offshore as gillnet protection. Both the IUCN and IUCN+ options are highly effective (Figure 5) as the 100m depth contour is well offshore here (Figure A.2). Current bycatch exceeds the New Zealand government threshold for total human impact (PST: Population Sustainability Threshold) for all areas except the west coast of the South Island, and exceeds the US government threshold (PBR: Potential Biological Removal) for all areas (Figure 5).Allee Effect runs for the South-east population show that the birth rate is lower in smaller populations (Figure 6).

**Figure 6.**
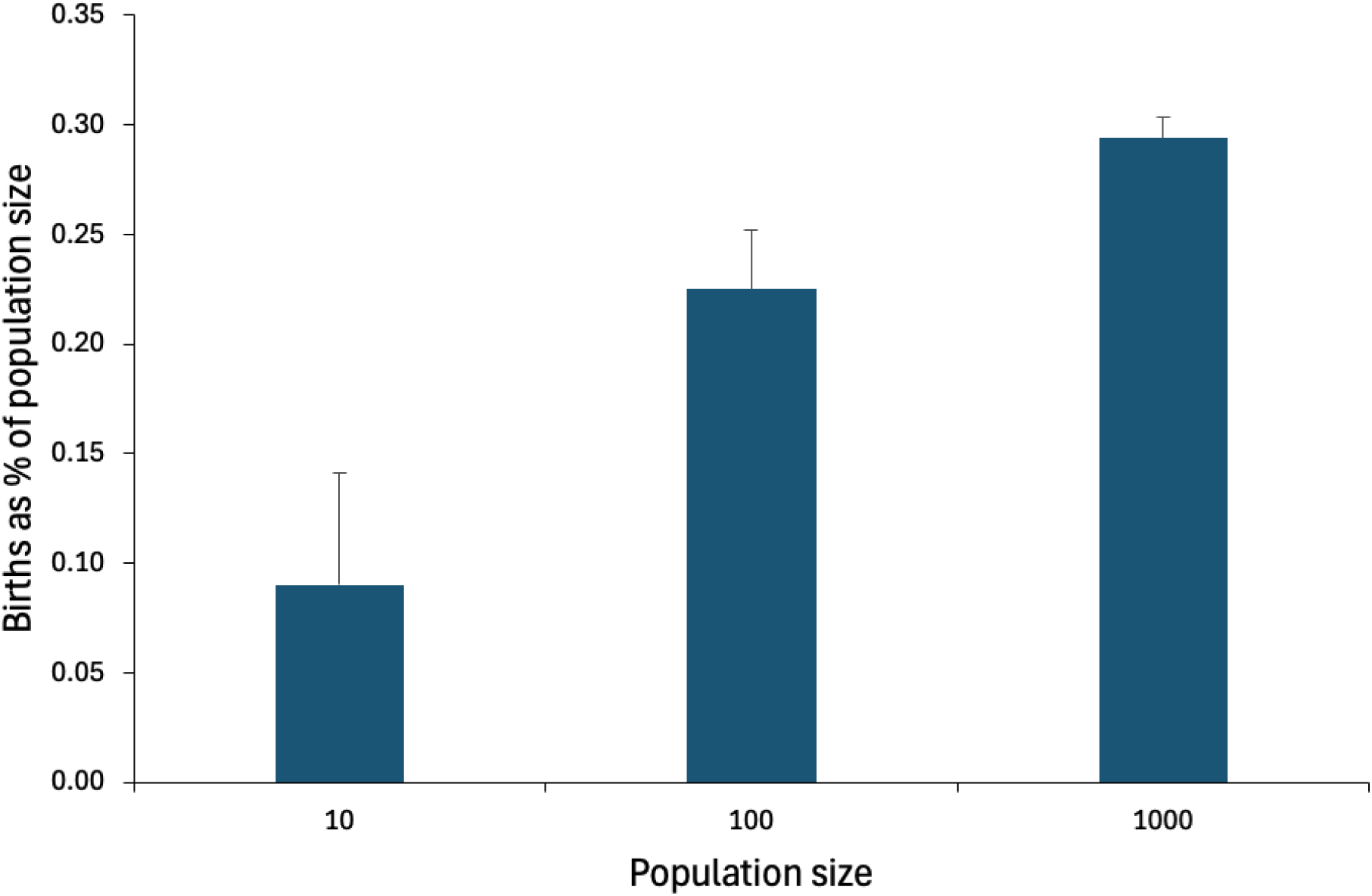
Model runs for the South-east population with different population densities showed lower reproductive rates for smaller populations, due to the Allee effect which requires a mature male to be available nearby for a mature female to become pregnant (n = 10 runs, 95% Confidence Intervals shown).

## Discussion

Annual dolphin bycatch has declined over time, as a result of improved dolphin protection. However, current protection is less effective than previously thought (e.g. Roberts et al. 2019, TMP 2019, MPI 2021). Current bycatch in commercial gillnet and trawl fisheries exceeds national and international limits on total human impact. Additional threats to New Zealand dolphins include recreational and customary fisheries, aquaculture, indirect impacts of fishing and aquaculture on dolphins (e.g. habitat modification, reduced prey availability), pollution, disease and marine mining. Therefore, it is important to reduce commercial fisheries bycatch to levels well below thresholds for total human impact. Previous analyses used outdated catch rates, from before camera monitoring (MPI 2021, 2025a,b), and a habitat model to predict dolphin distribution based on turbidity (Roberts et al. 2019). The habitat model resulted in lower estimated bycatch but a very poor match to the survey data (IWC 2023). Our analysis uses dolphin distribution alongshore and offshore directly from population survey data (e.g. MacKenzie and Clement 2014, 2016, 2019). Survey data show that the overlap between dolphins and fisheries causing bycatch has been reduced, but not by as much as the habitat model (Roberts et al. 2019) suggests.

Current bycatch of the Critically Endangered Māui dolphin sub-species is estimated at one dolphin every 1.7 years. This is more than five times higher than the threshold set by the New Zealand government (TMP 2019) for total human impact and unsustainable given the current Māui population of around 48 individuals. The IWC (2024) has recommended immediately eliminating Māui dolphin bycatch. Protection has gradually increased over time, however estimates of bycatch have also increased as better information becomes available from more extensive observer or camera monitoring. This study shows that increasing the size of the area where dolphins are protected is an effective way to reduce fisheries mortality. The level of protection recommended by the IUCN and IWC would reduce bycatch to levels approaching zero. The IUCN+ level of protection is substantially better than the IUCN recommendation for Māui dolphins off the North Island and the Hector’s dolphin population off north-east South Island. The New Zealand government’s goal is to “reduce the mortality of non-target species from marine fisheries to zero” (MPI 2021 p.12). This can be achieved for Hector’s and Māui dolphins by increasing protection to the IUCN+ level.

Future opportunities for improving the model include adding dolphin foraging, since prey availability is likely to be an important driver of dolphin movements. However, most of the fish and squid consumed by New Zealand dolphins are not target species of fisheries and there has been very little research on these prey species (Miller et al. 2013). In our model dolphin movements are based on their known preferences for shallow waters and their attraction to trawling. Once information on the distribution and movements of dolphin prey becomes available, we plan to add it to the model.

Advantages of ABMs include realistic dolphin movements, with dolphins moving into and out of protected areas rather than the ‘static’ approach used in most models estimating overlap between dolphins and fishing. ABMs are well suited for situations with source-sink dynamics, where areas with high human impact (sink) deplete sub-populations in neighboring protected areas (source) due to dolphin migration from protected to unprotected areas. Realistic Allee effects can be easily included in individual-based models and ABMs, showing how population reductions due to human impacts can reduce the potential for recovery. For example, Kuparinen et al. (2014) showed both reduced rates of recovery and increased uncertainty in recovery time frames for small populations. ABMs have been used to study bycatch of marine mammals, seabirds and turtles (e.g. Nabe-Nielsen et al. 2014; van Beest et al. 2017; Lusseau et al. 2023) and would benefit many other species such as vaquita, humpback dolphins and river dolphins. Our model could be adapted to study road kill (e.g. Kramer-Schadt et al. 2007), polar bears scavenging around human settlements, orca and sperm whales removing fish from long-lines (e.g. Nolan et al. 2006; Amelot et al. 2022). ABMs have also been used to study ship strike of marine mammals (e.g. Baille and Zitterbart 2022), including whale movements and behavior around different types of vessels, avoidance strategies based on vessel proximity, speed and trajectory to better quantify the risk of collision and the effectiveness of mitigation (Dunlop 2024).

ABMs facilitate research on displacement of human impacts outside protected areas, which can result in moving conservation problems from one area to another rather than solving the problem (e.g. Lusseau et al. 2023). Risk models based simply on the statistical overlap of protected species and human impacts fail to include dynamic responses within the system, such as an impact being displaced rather than removed. Our model is also able to assess population impacts of likely future distributions of human impacts, making it ideal for testing different potential conservation options. With the current model we are thus finally able to provide recommendations for protecting the threatened New Zealand dolphin, while taking into account the dynamic nature of fisheries and their impacts on the species. If the IUCN+ level of protection were implemented soon, this would greatly facilitate the recovery of New Zealand dolphins. It would also ensure that New Zealand complies with international standards such as the Import Rule under the US Marine Mammal Protection Act.

## Supporting information

A. Additional Figures.
B. Detailed model description following the ODD protocol (Overview, Design concepts and Details) for describing individual- and agent-based models (Grimm et al. 2020).
C. Model calibrations.

## References

Amelot M, Plard F, Guinet C, Arnould JPY, Gasco N, Tixier P. 2022. Increasing numbers of killer whale individuals use fisheries as feeding opportunities within subantarctic populations. Biology Letters 18: 20210328.

Authier M. 2024. Despite the perception that it is rare, by-catch of marine mammals can lead to population decline. ICES Journal of Marine Science fsae151 10.1093/icesjms/fsae151

Avila IC, Kaschner K, Dormann CF. 2018. Current global risks to marine mammals: taking stock of the threats. Biological Conservation 221: 44–58.

Baille LMR, Zitterbart DP. 2022. Effectiveness of surface-based detection methods for vessel strike mitigation of North Atlantic right whales. Endangered Species Research 49: 10.3354.

Baird S, Bradford E. 2000. Estimation of Hector’s dolphin bycatch from inshore fisheries, 1997/98 fishing year. Report for Department of Conservation, Wellington, New Zealand.

Ballance LT, Gerrodette T, Lennert-Cody C, Pitman RL, Squires D. 2021. A history of the tuna-dolphin problem: Successes, failures, and lessons learned. Frontiers of Marine Science 8: 1700.

Bennington S. 2025. Investigating the recovery potential of Hector’s dolphin *(Cephalorhynchus hectori)*. PhD thesis, University of Otago, New Zealand.

Bräger S, Dawson SM, Slooten E, Smith S, Stone GS, Yoshinaga A. 2002. Site fidelity and along-shore range in Hector’s dolphin, an endangered marine dolphin from New Zealand. Biological Conservation 108: 281–287.

Bräger S, Bräger Z. 2018. Range utilization and movement patterns of coastal Hector’s dolphins *(Cephalorhynchus hectori)*. Aquatic Mammals 44: 633–642.

Brownell RL, Reeves RR, Read AJ, Smith BD, Thomas PO, Ralls K, Amano M, Berggren P, Myo Chit A, Collins T, Currey R, Dolar ML, Genov T, Hobbs RC, Kreb D, Marsh H, Zhigang M, Perrin WF, Phay S, Rojas-Bracho L, Ryan GE, Shelden KEW, Slooten E, Taylor BL, Vidal O, Ding W, Whitty TS, Wang JY. 2019. Bycatch in gillnet fisheries threatens critically endangered small cetaceans and many others. Endangered Species Research 40: 285–296.

Burgess MG, McDermott GR, Owashi B, Peavey Reeves LE, Clavelle T, Ovando D, Wallace BP, Lewison RL, Gaines SD, Costello C. 2018. Protecting marine mammals, turtles, and birds by rebuilding global fisheries. Science 359: 1255–58.

Constantine R. 2023. Cephalorhynchus hectori ssp. maui. The IUCN Red List of Threatened Species 2023: 10.2305/IUCN.UK.2023-1.RLTS.T39427A50380174.en.

Constantine R, Steel D, Carroll E, Hamner RM, Hansen C, Hickman G, Hillock K, Ogle M, Tukua P, Baker CS. 2021. Estimating the abundance and effective population size of Māui dolphins *(Cephalorhynchus hectori maui)* in 2020-2021 using microsatellite genotypes, with retrospective matching to 2001. Report to Department of Conservation, Auckland, New Zealand.

Cooke JG, Constantine R, Hamner RM, Steel D, Baker CS. 2019. Population dynamic modeling of the Māui dolphin based on genotype capture-recapture with projections involving bycatch and disease risk. Report to Ministry for Primary Industries.

Daly E, White M. 2021. Bottom trawling noise: Are fishing vessels polluting to deeper acoustic habitats? Marine Pollution Bulletin 162: 111877.

Dawson SM, Slooten E. 1988. Hector’s Dolphin *Cephalorhynchus hectori:* Distribution and abundance. Reports of the International Whaling Commission Special Issue 9: 315–324.

Dawson S, Slooten E, Dufresne S, Wade P, Clement D. 2004. Small-boat surveys for coastal dolphins: Line-transect surveys for Hector’s dolphins *(Cephalorhynchus hectori)*. Fishery Bulletin 102: 441–451.

de Jager M, Hengeveld GM, Mooij WM, Slooten E. 2019. Modelling the spatial dynamics of Māui dolphins using individual-based models. Ecological Modelling 402: 59–65.

DOC 2025. Department of Conservation database of New Zealand dolphin mortalities: hectors-and-maui-dolphin-incident-database

Dragonfly Consulting 2025. Protected species bycatch in New Zealand fisheries. https://psc.dragonfly.co.nz/2018v1/released/

Dunlop R. 2024. Use of behavioural response method to assess the risk of collision between migrating humpback whales and vessels. Marine Pollution Bulletin 199: 115986

Elliott B, Tarzia M, Read AJ. 2023. Cetacean bycatch management in regional fisheries management organizations: current progress, gaps, and looking ahead. Frontiers of Marine Science 9:1006894.

Fletcher D, Dawson S, Slooten E. 2002. Designing a mark-recapture study to allow for local emigration. Journal of Agricultural, Biological and Environmental Statistics 7(4): 1–8.

Frankish CK, Christ A, de Jong F, Tougaard J, Teilmann J, Dietz R, von Benda-Beckmann AM, Binnerts B, Nabe-Nielsen J. 2023. Effect of ship noise on the behaviour of harbour porpoises *(Phocoena Phocoena)*. Marine Pollution Bulletin 197.

GFW. 2025. Global Fishing Watch. Online database of fishing activity using AIS, satellite imagery and AI: https://globalfishingwatch.org/

Gormley AM. 2009. Population modelling of Hector’s dolphins. PhD thesis, University of Otago, Dunedin, New Zealand.

Gormley AM, Slooten E, Dawson SM, Barker RJ, Rayment W, du Fresne S, Bräger S. 2012. First evidence that marine protected areas can work for marine mammals. Journal of Applied Ecology 49:474–480.

Grimm V, Railsback SF. 2005. Individual-Based Modeling and Ecology. Princeton University Press.

Grimm V, Railsback SF, Vincenot CE, Berger U, Gallagher C, DeAngelis DL, Ayllón, D et al. 2020. The ODD Protocol for Describing Agent-Based and Other Simulation Models: A Second Update to Improve Clarity, Replication, and Structural Realism. Journal of Artificial Societies and Social Simulation 23: 7.

Grimm V, Revilla E, Berger U, Jeltsch F, Mooij WM, Railsback SF, Thulke H-H, Weiner J, Wiegand T, DeAngelis DL. 2005. Pattern-oriented modeling of agent-based complex systems: lessons from ecology. Science 310, 987–991.

Haider HS, Oldfield SC, Tu T, Moreno RK, Diffendorfer JE, Eager EA, Erickson RA. 2017. Incorporating Allee effects into the potential biological removal level. Natural Resource Modeling 2017: e12133

Halpern BS, Walbridge S, Selkoe KA, Kappel CV, Micheli F, D’Agrosa C, Bruno JF, et al. 2008. A global map of human impact on marine ecosystems. Science 319: 948–52.

Halpern BS, Frazier M, Afflerbach J, Lowndes JS, Micheli F, O’Hara C, Scarborough C, Selkoe KA. 2019. Recent pace of change in human impact on the World’s ocean. Scientific Reports 9: 11609.

Harvey M. 2021. Ecology and conservation of Hector’s dolphins in Porpoise Bay, Southland. MSc thesis, University of Otago, New Zealand.

IUCN 2012. Resolution 142. Actions to avert the extinctions of rare dolphins: Māui dolphins, Hector’s dolphins, Vaquita porpoises and South Asian river and freshwater dependent dolphins and porpoises. https://portals.iucn.org/library/sites/library/files/resrecfiles/WCC_2012_REC_142_EN.pdf

IWC 2023. International Whaling Commission, Report of the 2023 Scientific Committee Meeting. 2023 Scientific Committee Report

IWC 2024. International Whaling Commission, Report of the 2024 Scientific Committee Meeting. 2024 Scientific Committee Report

Jaramillo-Legorreta AM, Cardenas-Hinojosa G, Nieto-Garcia E, et al. 2019. Decline towards extinction of Mexico’s Vaquita Porpoise *(Phocoena sinus)*. Royal Society Open Science 6:190598.

Kramer-Schadt S, Revilla E, Wiegand T, Grimm V. 2007. Patterns for parameters in simulation models. Ecological Modelling 204: 553–556.

Kuparinen A, Keith DM, Hutchings JA. 2014. Allee Effect and the uncertainty of population recovery. Conservation Biology 28: 790–798.

Lamb J, Gulka J, Adams E, Cook A, Williams KA. 2024. A synthetic analysis of post-construction displacement and attraction of marine birds at offshore wind energy installations. Environmental Impact Assessment Review 108: 107611.

Lewison RL, Crowder LB, Read AJ, Freeman SA. 2004. Understanding impacts of fisheries bycatch on marine megafauna. Trends in Ecology and Evolution 19: 598–604.

Lusseau D, Kindt-Larsen L, van Beest FM. 2023. Emergent interactions in the management of multiple threats to the conservation of harbour porpoises. Science of the Total Environment 855: 158936.

MacKenzie DL, Clement DM. 2014. Abundance and distribution of South Island east coast Hector’s dolphin. New Zealand Aquatic Environment and Biodiversity Report 123, Report to Ministry for Primary Industries, Wellington, New Zealand.

MacKenzie DL, Clement DM. 2016. Abundance and distribution of South Island west coast Hector’s dolphin. New Zealand Aquatic Environment and Biodiversity Report 168, Report to Ministry for Primary Industries, Wellington, New Zealand.

MacKenzie DL, Clement DM. 2019. Abundance and distribution of Hector’s dolphin on South Coast South Island. New Zealand Aquatic Environment and Biodiversity Report 236, Report to Ministry for Primary Industries, Wellington, New Zealand.

McGrath G. 2020. The history of New Zealand / Aotearoa dolphins *Cephalorhynchus hectori* abundance and distribution. MSc thesis, University of Otago, Dunedin, New Zealand. https://ourarchive.otago.ac.nz/handle/10523/10467

Miller EJ, Lalas C, Dawson S, Ratz H, Slooten E. 2013. Hector’s dolphin diet: The species, sizes and relative importance of prey eaten by *Cephalorhynchus hectori*, investigated using stomach content analysis. Marine Mammal Science 29: 606–628.

MPI 2021. Protecting South Island Hector’s dolphins. Further consultation on fisheries measures. October 2021.

MPI 2023 and 2024. Ministry for Primary Industries provided data on fishing effort and protected area boundaries to Elisabeth Slooten. Government fisheries data are also available publicly, in a summarised form, at: Dragonfly.co.nz

MPI 2025a. Ministry for Primary Industries: Online data on bycatch of seabirds and marine mammals. seabirds-and-protected-marine-species-caught-by-commercial-fishers

MPI 2025b. Ministry for Primary Industries: South Island Hector’s dolphin bycatch reduction plan. Annual Report 2023/24.

Nabe-Nielsen J, Sibly RM, Tougaard J, Teilmann J, Sveegaard S. 2014. Effects of noise and by-catch on a Danish harbour porpoise population. Ecological Modelling 272: 242–251.

Nabe-Nielsen J, van Beest FM, Grimm V, Sibly RM, Teilmann J, Thompson PM. 2018. Predicting the impacts of anthropogenic disturbances on marine populations. Conservation Letters, 11(5): e12563

NIWA 2016. New Zealand Regional Bathymetry. National Institute of Water and Atmospheric Research. https://niwa.co.nz/our-science/oceans/bathymetry/

Nolan CP, Liddle GM, Elliot J. 2006. Interactions between killer whales *(Orcinus orca)* and sperm whales *(Physeter macrocephalus)* with a longline fishing vessel. Marine Mammal Science 16: 658–664.

Railsback SF, Grimm V. 2019. Agent-based and individual-based modelling. Second Edition, Princeton University Press.

Rayment W, Dawson SM, Slooten E. 2010. Seasonal changes in distribution of Hector’s dolphin at Banks Peninsula, New Zealand: implications for protected area design. Aquatic Conservation: Marine and Freshwater Ecosystems 20: 106–116.

Rayment W, Dawson SM, Slooten E, Bräger S, DuFresne S, Webster T. 2009. Kernel density estimates of alongshore home range of Hector’s dolphins *(Cephalorhynchus hectori)* at Banks Peninsula. Marine Mammal Science 25: 537–556.

Rayment W, Webster T. 2009. Observations of Hector’s dolphins *(Cephalorhynchus hectori)* associating with inshore fishing trawlers at Banks Peninsula, New Zealand. New Zealand Journal of Marine and Freshwater Research 43: 911–916.

Read AJ, Drinker P, Northridge S. 2006. By-catches of marine mammals in US fisheries and a first attempt to estimate the magnitude of global marine mammal by-catch. Conservation Biology 20:163–169.

Roberts JO, Webber DN, Roe WD, Edwards CTT, Doonan IJ. 2019. Spatial risk assessment of threats to Hector’s and Māui dolphins. New Zealand Aquatic Environment and Biodiversity Report 214, Ministry for Primary Industries, Wellington, New Zealand https://www.fisheries.govt.nz/dmsdocument/35007

Slooten E. 1991. Age, growth and reproduction in Hector’s dolphins. Canadian Journal of Zoology 69: 1689–1700

Slooten E. 2007. Conservation management in the face of uncertainty: Effectiveness of four options for managing Hector’s dolphin bycatch. Endangered Species Research 3:169–179.

Slooten E. 2013. Effectiveness of area-based management in reducing bycatch of the New Zealand dolphin. Endangered Species Research 20: 121–130.

Slooten E. 2020. Effectiveness of current protection for Māui dolphin. Journal of Cetacean Research and Management 21: 151–155.

Slooten E, Davies N. 2012. Hector’s dolphin risk assessments: Old and new analyses show consistent results. Journal of the Royal Society of New Zealand 42: 49–60.

Slooten E, Dawson SM. 2010. Assessing the effectiveness of conservation management decisions: Likely effects of new protection measures for Hector’s dolphin. Aquatic Conservation: Marine and Freshwater Ecosystems 20: 334–347.

Slooten E, Dawson SM. 2021. Delays in protecting a small endangered cetacean: Lessons learned for science and management. Frontiers in Marine Science 8: 2021. doi10.3389/fmars.2021.606547

Slooten E, Dawson SM, Rayment WJ. 2004. Aerial surveys for coastal dolphins: Abundance of Hector’s dolphins off the South Island west coast, New Zealand. Marine Mammal Science 20: 447–490.

Slooten E, Dawson SM, Rayment WJ, Childerhouse SJ. 2006. A new abundance estimate for Maui’s dolphin: What does it mean for managing this critically endangered species? Biological Conservation 128: 576–581.

Slooten E, Dawson SM, Whitehead H. 1993. Associations among photographically identified Hector’s dolphins. Canadian Journal of Zoology 71: 2311–2318.

Slooten E, Fletcher D, Taylor BL. 2000. Accounting for uncertainty in risk assessment: Case study of Hector’s dolphin mortality due to gillnet entanglement. Conservation Biology 14: 1264–1270.

Slooten E, Lad F. 1991. Population biology and conservation of Hector’s dolphin. Canadian Journal of Zoology 69: 1701–1707.

Starr P, Langley A. 2000. Inshore fishery observer programme for Hector’s dolphins in Pegasus Bay, Canterbury Bight, 1997/98. Report for Department of Conservation, Wellington

Stillman RA, Railsback SF, Giske J, Berger U, Grimm V. 2015. Making predictions in a changing World: The benefits of individual-based ecology. BioScience 65: 140–50.

Stone G, Kraus S, Hutt A, Martin S, Yoshinaga A. 1997. Reducing by-catch: Can acoustic pingers keep Hector’s dolphins out of fishing nets? MTS Journal 31: 3–7.

TMP 2019. Threat Management Plan for Hector’s and Māui dolphins. 2019. Fisheries New Zealand and Department of Conservation. June 2019.

Turek J, Slooten E, Dawson S, Rayment W, Turek D. 2013. Distribution and abundance of Hector’s dolphins off Otago, New Zealand. New Zealand Journal of Marine and Freshwater Research 47: 181–191.

United Nations Office of Legal Affairs. 2021. The Second World Ocean Assessment: World Ocean Assessment II - Volume I & II. United Nations.

van Beest FM, Kindt-Larsen L, Bastardie F, Bartolino V, Nabe-Nielsen J. 2017. Predicting the population-level impact of mitigating harbor porpoise bycatch with pingers and time-area fishing closures. Ecosphere 8: e01785

Wade PR, Long KJ, Francis TB, Punt AE, Hammond PS, Heinemann D, et al. 2021. Best practices for assessing and managing bycatch of marine mammals. Frontiers of Marine Science 8. doi: 10.3389/fmars.2021.757330

Weir CR, van Waerebeek K, Jefferson TA et al. 2011. West Africa’s Atlantic Humpback Dolphin *(Sousa teuszii):* endemic, enigmatic and soon endangered? African Zoology 46: 1–17.

Whitehead H. 1987. Social organization of sperm whales off the Galapagos: Implications for management and conservation. Reports of the International Whaling Commission 37: 195–199.

Whitehead H, Weilgart L. 2000. The sperm whale: Social females and roving males. In: Mann J, Connor RC, Tyack P, Whitehead H (eds) Cetacean Societies, University of Chicago Press, Chicago, pp 154–172.

Wickman L. 2024. Evaluating the long-term impacts of area-based protection on Hector’s dolphins at Banks Peninsula, New Zealand. PhD thesis, University of Otago, New Zealand.

Wiegand K, Henle K, Sarre SD. 2002. Extinction and spatial structure in simulation models. Conservation Biology 16: 117–128.

Wiegand T, Jeltsch F, Hanski I, Grimm V. 2003. Using pattern-oriented modeling for revealing hidden information: a key for reconciling ecological theory and application. Oikos 100: 209–222.

Wilensky U. 1999. NetLogo. http://ccl.northwestern.edu/netlogo/. Center for Connected Learning and Computer-Based Modeling, Northwestern University, Evanston, IL, USA.

Williams, H. 2022. Abundance and distribution of Hector’s dolphins off the coast of Dunedin, New Zealand, and overlap with commercial fishing. MSc thesis, University of Otago, New Zealand.

