## Supplemental material for "An agent-based approach for designing effective protection"

### Supporting information A: Additional figures

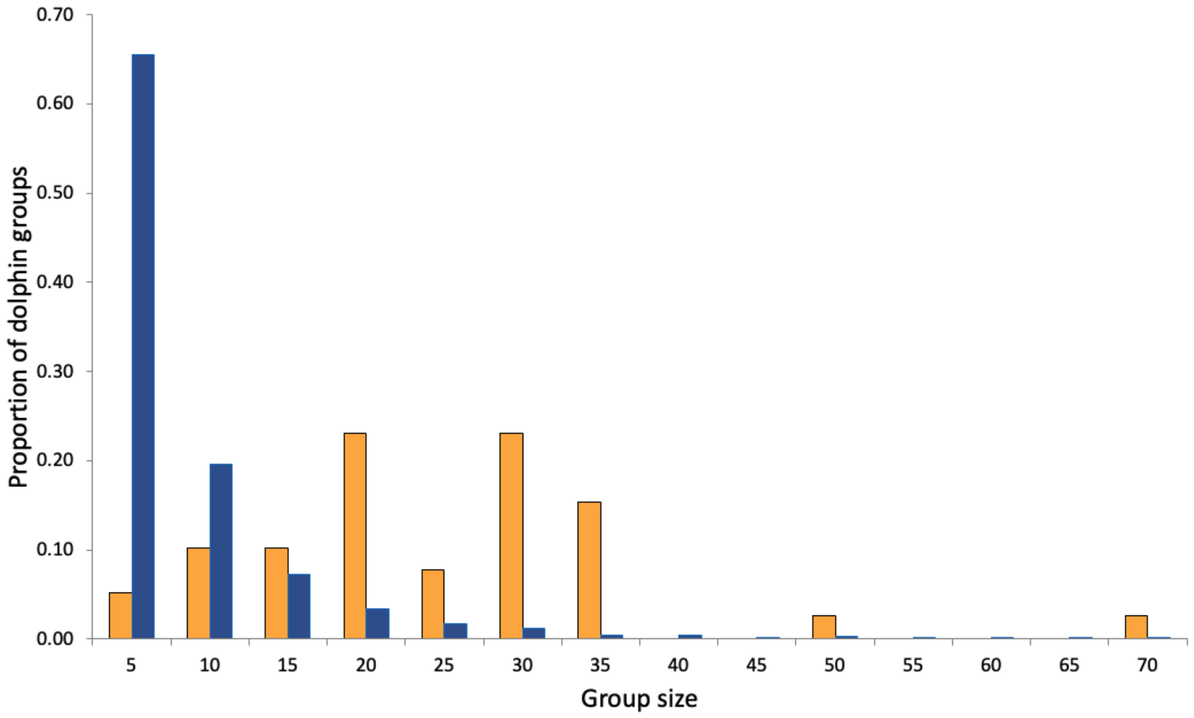

**Figure A.1:** Group size of Hector's dolphins observed in the wild between 1991 and 2020 following trawlers (orange bars, N=39) and not following trawlers (blue bars, N=9,345).

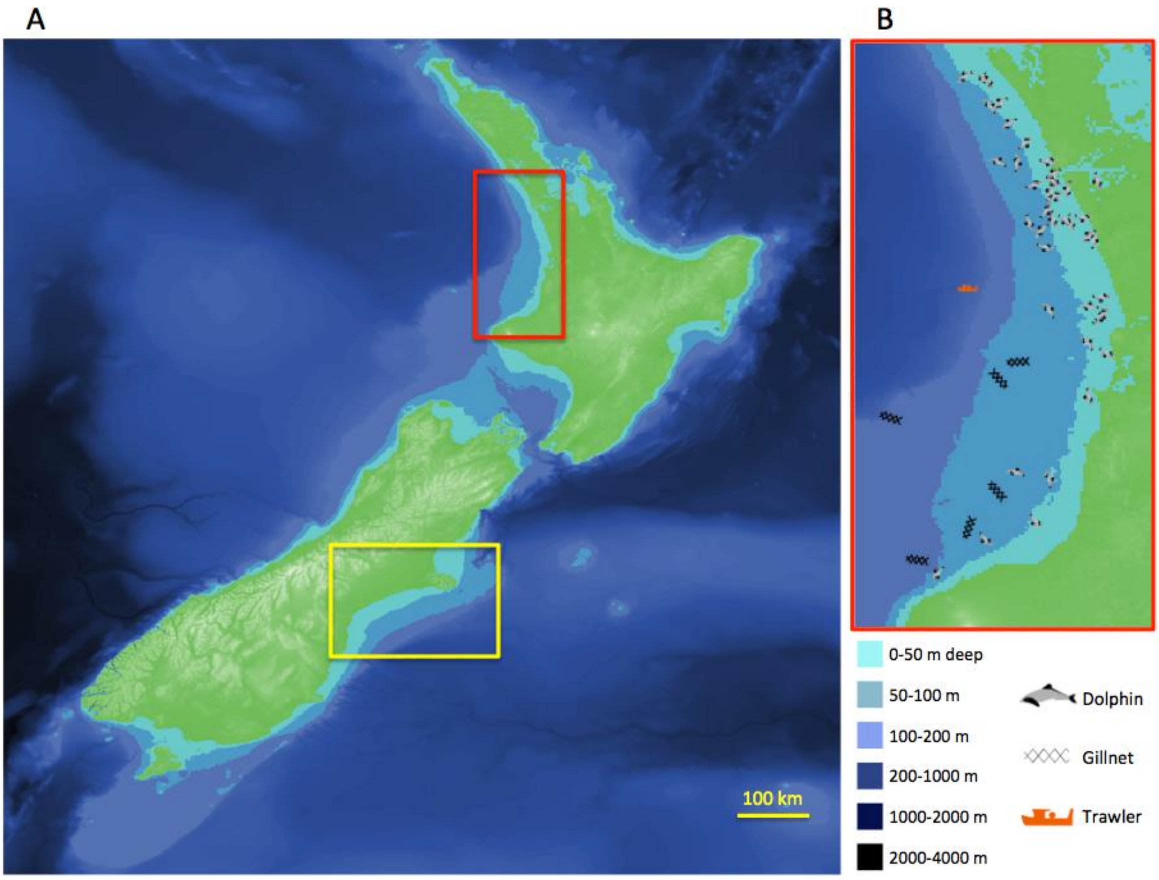

**Figure A.2** The model landscape includes New Zealand's North and South Islands (A), with the red box showing a snapshot from the simulation model for part of the west coast of the North Island (B) and the yellow box indicating the Banks Peninsula to Timaru area where high observer coverage and high dolphin densities have resulted in the vast majority of the observed dolphin catches (the size of the dolphins, gillnets and trawlers has been exaggerated to make them visible on the map).

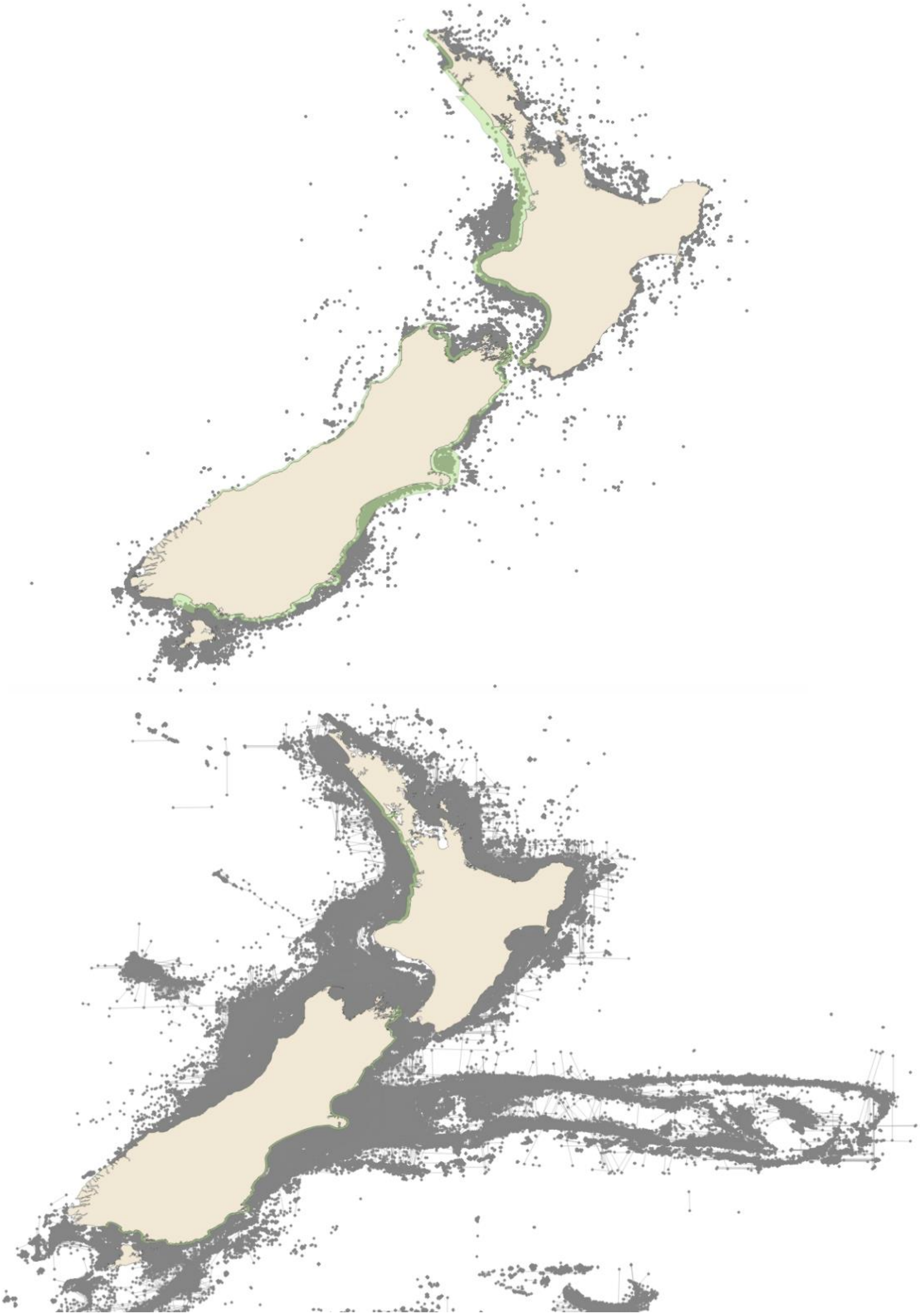

**Figure A.3** Fishing effort (grey) and current dolphin protection (green) for gillnet (above) and trawl fishing (below). Grey lines show trawls for which start and end locations were available. Grey dots show gillnet sets (above) and trawls for which only a start location was available.

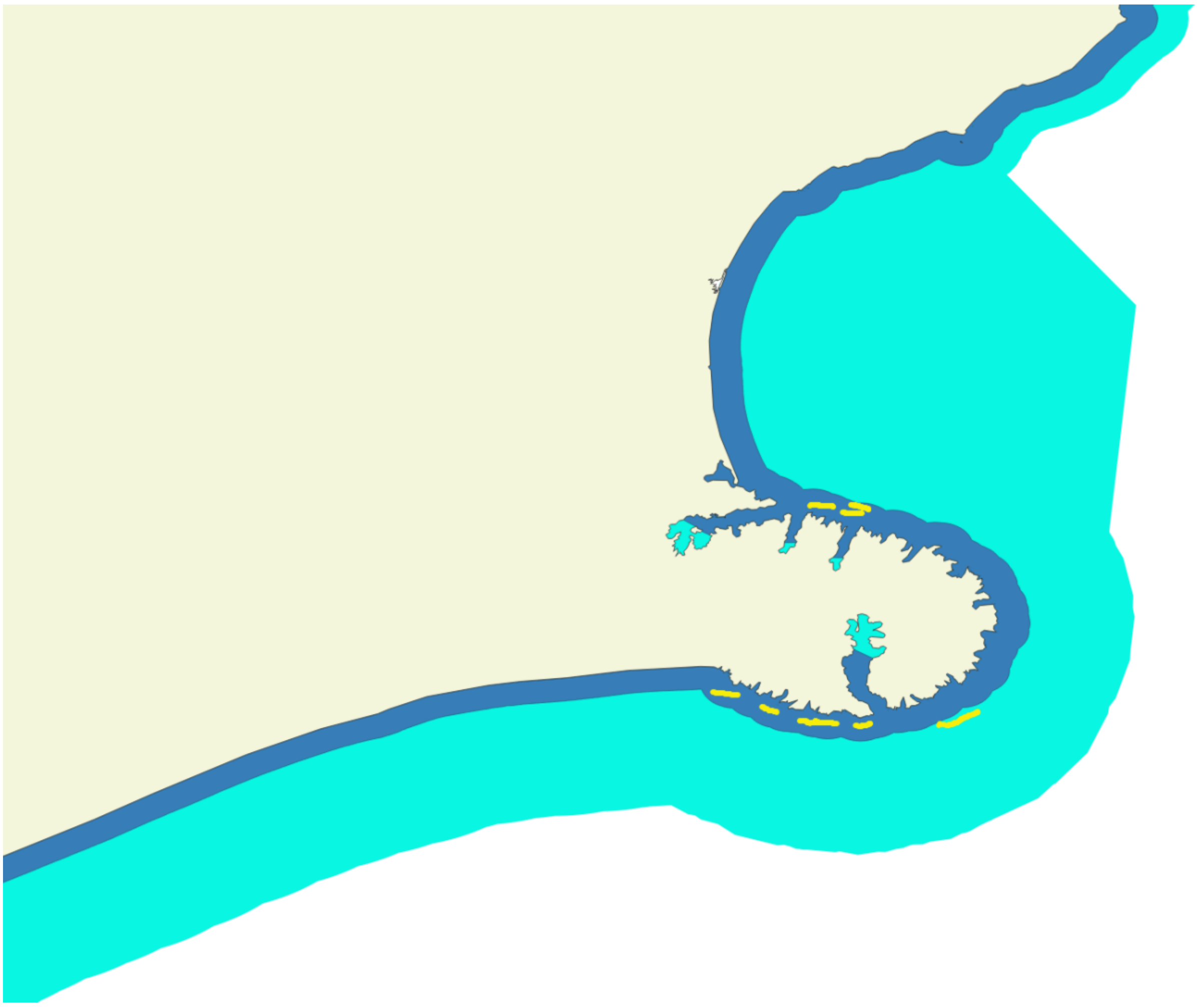

**Figure A.4** Trawlers are allowed to fish in areas where trawling is banned (dark blue) under an exemption for small trawlers. The yellow lines show the track of a research vessel following some of these trawlers. In the light blue area gillnets are banned but trawling is permitted.

### Supporting info B: ODD

The model description follows the ODD (Overview, Design concepts, Details) protocol for describing individual- and agent-based models (Grimm et al., 2006), as updated by Grimm et al. (2010, 2020).

#### 1. Purposes and Patterns

The **purpose** of our model is to predict the effects of gillnet and trawler by-catch deaths on the long-term population dynamics of Māui and Hector's dolphins around New Zealand under various protection schemes and levels of fishing effort, incorporating realistic dolphin and fishing vessel movements and densities, and considering how reduced population densities affect the likelihood of mating.

To consider our model realistic enough for its purpose, we evaluated its success at reproducing the following **patterns**: 1. dolphin movements in their home range in response to water depth and distance from shore, 2. movement of dolphins attracted to trawlers, 3. dolphin group size variation, 4. trawler movements and distribution of trawl effort, 5. bycatch rates for gillnets and trawlers.

We considered a realistic overlap in space and time between dolphins, trawlers and gillnets, combined with realistic bycatch rates, to be sufficient for producing realistic mortality patterns. We used the pattern-oriented modelling approach (POM) (Wiegand et al. 2003; Grimm et al. 2006; Kramer-Schadt et al. 2007) to evaluate the model entities' movements and confirm realistic behaviors during two phases of the modelling process: during development to parameterize the model and during evaluation to test if model outputs are realistic.

#### 2. Entities, State Variables, and Scales

The model world is 1024 x 1024 patches, each representing 1.63 x 1.63 km. The world thus covers ~ 2.786 x $10^6$ $km^2$. This area includes the entire land mass of the North and South Islands of New Zealand (~269,000 $km^2$) with the remainder representing the surrounding ocean (NIWA 2016). There is no world-wrapping. One time step in the model (tick) represents 1 hour.

The model includes the following entities: dolphins, trawlers, gillnets, ports, waypoints (navigation reference points for trawlers near ports), landscape patches (land or ocean) and placement areas based on fisheries statistical areas and dolphin surveys. Dolphins are "super-agents"; each represents a group of individual dolphins of various ages and genders. They are characterized by their sub-species (Hector's or Māui), group size, age, gender, estrus/pregnancy/post-partum status of females, velocity, traveling direction, home range, and location (see Table B.1 for details). They are also characterized by whether they move towards or flock around a trawler, or if they "wander" within their home range when trawlers are not nearby. Trawlers are characterized by their size, home port according to data provided by MPI (2023, 2024), location, speed, destination, activity (trawling, in port, etc.). They are also characterized by several variables that allow for fine-tuning where to fish, length of trawls, length of fishing trips, time in port between fishing trips, etc. Gillnets are characterized by location and soak time. Ports are characterized by name and geographic location. Waypoints are characterized by location and associated port. Table B.1 lists all entities and associated state variables.

**TABLE B.1 Entities and State Variables:**

| Entity | Type of state variable | Name of state variable | Units | Meaning | Procedure in which variable is updated | Values |
|---|---|---|---|---|---|---|
| Ports | Static | name | string | name of port | setup-ports | e.g., “Akaroa” |
| | | inbound_waypoints_list | list | list of waypoints for trawlers at this port to use | setup-ports-waypoints-list | |
| | | outbound_waypoints_list | degrees | list of waypoints for trawlers at this port to use | setup-ports-waypoints-list | |
| | | min_max_bearing_list | degrees | list of minimum and maximum navigation bearings to safely leave port (i.e., avoid land) | setup-ports-min-max-bearing-list | |
| Waypoints | Static | my_port | string | ID of associated port | setup-outbound-waypoints-list | e.g., port 2 |
| | | my_port_name | string | name of associated port; e.g., “Akaroa” | setup-outbound-waypoints-list | e.g., “Akaroa” |
| Gillnets | Static | gillnet_area | string | geographical area where gillnet is assigned | setup-gillnets | e.g., “banks-banksnorth” |
| | Dynamic | deployed? | T/F | whether gillnet is actively fishing or not | gillnets-decide-if-deployed | |
| Dolphins | Static | subspecies | string | identifies subspecies of dolphin group | setup-dolphins | “maui” or “hectors” |
| | | female? | list | list that indicates gender of each dolphin in group; e.g. T= female and F = male | setup-dolphins | T/F |
| | Dynamic | dolphin_distance_band | string | general description based on distance from shore; “near”, “middle” or “far” | dolphin-move | “near “= 0-4, “middle” = 4-12, “far” = 12-20 |

| | | | | | | |
|---|---|---|---|---|---|---|
| | | | | | | nautical miles from shore |
| | | dolphin_area | string | general location of dolphin; e.g., “south-otago” | setup-dolphins; dolphin-switch-area | |
| | | home_patch | patch ID | patch approximately in middle of dolphin’s home range; e.g., patch 22 44 | setup-dolphins; dolphin-switch-home-patch; dolphin-group-split | |
| | | home_range_shoreline_length_km | km | shoreline length of dolphin home range drawn from random normal distribution | dolphin-select-home-range-shoreline-length | μ = 50km, σ = 2.5km |
| | | home_range_limit_patch_list | ID’s (characters) | two shoreline patches at each end of home range | dolphin-setup-home-range-limit-patch-list | |
| | | home_range_orientation | | approximate orientation of shoreline | dolphin-setup-home-range-limit-patch-list | “N-S” for home ranges than run predominately north-to-south; “E-W” for those that run east-to-west |
| | | home_range_shoreline_midpoint_patch | ID (characters) | shoreline patch in the middle of home range | dolphin-setup-home-patch-and-home-range | |
| | | age | years | list of individual dolphin ages in group | dolphin-age | 0-31 |
| | | estrus_start_days | days | list of Julian days when individual dolphins begin | dolphin-setup-estrus-start-days | range is Julian days 1-52 |

| | | | | | | |
|---|---|---|---|---|---|---|
| | | | | estrus (varies with age) | | |
| | | estrus? | boolean | individual estrus status of each dolphin in group | dolphin-set-estrus-status | True or false |
| | | postpartum_timer | days | list of individual dolphins' days before they are eligible to reproduce again drawn from triangular distribution | initialize-some-dolphins-postpartum-timer; dolphin-set-postpartum-timer; dolphin-update-postpartum-timer | range is 2-4 years with peak at 3 years |
| | | calf_timer | days | list of individual dolphins' days before giving birth; drawn from random-normal distribution | initialize-some-dolphins-pregnant; dolphin-select-gestation-days; dolphin-update-calf-timer; dolphin-give-birth | $\mu$ = 330 days; $\sigma$ = 10 days |
| | | female_ready_to_mate? | boolean | indicates whether there is $\geq$ 1 female ready to reproduce in group | dolphin-set-female-ready-to-mate-status; dolphin-give-birth; dolphin-group-split | True or false |
| | | mature_male? | boolean | indicates whether there is $\geq$ 1 mature male ($\geq$ six years old) in group | dolphin-set-mature-male-status; dolphin-give-birth; dolphin-group-split | True or false |

| | | | | | | |
|---|---|---|---|---|---|---|
| | | group_size | integer | count of individual dolphins in group | dolphin-die; dolphin-give-birth; dolphin-group-split; dolphin-group-merge | Range 1-max group size |
| | | activity | string | activity dolphin group is currently performing | dolphin-decide-what's-next | “wandering”, “traveling to trawler”, or “flocking with trawler” |
| | | velocity_km_per_hour | km/hr | dolphin velocity drawn from random gamma distribution | dolphin-select-velocity; dolphin-travel-to-trawler; dolphin-flock-with-trawler | Random 0.5-7 |
| | | trawling_nearby? | boolean | indicates whether dolphin has detected a trawler fishing nearby | dolphin-detect-trawling-nearby-and-decide-to-travel; dolphin-quit-flocking; dolphin-update-target-and-flocking-trawler-trawling-complete | True or false |
| | | target_trawler | trawler ID | specific trawler dolphin is moving to | dolphin-decide-what's-next; dolphin-update-target-and-flocking- | e.g., trawler 22 (n/a otherwise) |

| | | | | | | |
|---|---|---|---|---|---|---|
| | | | | | trawler-trawling-happening; dolphin-update-target-and-flocking-trawler-trawling-complete | |
| | | flocking_trawler | trawler ID | specific trawler dolphin is flocking with | dolphin-decide-what's-next; dolphin-update-target-and-flocking-trawler-trawling-happening; dolphin-update-target-and-flocking-trawler-trawling-complete | e.g., trawler 22 (n/a otherwise) |
| | | wandering_heading | degrees | heading for dolphin group | dolphin-wander; dolphin-select-wandering-heading-biased-vector | |
| | | hours_flocking_with_current_trawler | hours | time dolphin has spent with current trawler | dolphin-decide-what's-next; dolphin-select-flocking-limit; dolphin-update-flocking-time-data; | 0-max allowed flocking time |

| | | | | | | |
|---|---|---|---|---|---|---|
| | | | | | dolphin-quit-flocking | |
| | | consecutive_flocking_hours_limit | hours | maximum time dolphin is allowed to stay with same trawler | dolphin-decide-what's-next; dolphin-select-flocking-limit | Random uniform; 3-4.99 hours |
| | | allowed_to_flock_countdown | hours | interval between dolphin flocking with trawlers | dolphin-update-allowed-to-flock-countdown; dolphin-quit-flocking | Random uniform; 6-9.99 hours |
| Trawlers | Static | my_size | string | "small" or "medium" | setup-trawlers | |
| | | my_length | meters | length of trawler | setup-trawlers | Small trawlers: 10 + random 6 (10-15) Medium trawlers: 16 + random 10 (16-25) |
| | | home_port | port ID | unique Netlogo identifier for home port; e.g., port 3 | setup-trawlers | |
| | | wake_up_time | Hours (time of day) | time of day trawler begins fishing trip | setup-trawlers | Random-uniform 04:00-07:00 |
| | | trawling_velocity_knots | knots | trawling speed | setup-trawlers | Small trawlers: 2.21 + (random-normal 0 0.5) Medium trawlers: 4.01 + (random-normal 0 0.5) |

| | | | | | | |
|---|---|---|---|---|---|---|
| | | travel_velocity_knots | knots | traveling speed | setup-trawlers | Small trawlers: 8 + (random-normal 0 1) Medium trawlers: 9 + (random-normal 0 1) |
| | | trawl_noise_dB | dB | source level noise of current trawl; attracts dolphins | trawler-select-trawl-noise | |
| | Dynamic | activity | string | current trawler activity | trawler-decide-what's-next; trawler-decide-what's-next-fishing-complete; trawler-decide-what's-next-fishing-incomplete; trawler-fishing-no-constraint; | "docked", "traveling", or "fishing" |
| | | destination | patch /waypoint/ port ID | trawler destination | trawler-decide-what's-next-fishing-complete; trawler-decide-what's-next-fishing-incomplete; trawler-select-next-waypoint; trawler- | e.g., patch 22 43 |

| | | | | | | |
|---|---|---|---|---|---|---|
| | | | | | fishing-edge-constraint; trawler-fishing-no-constraint | |
| | | velocity_knots | knots | actual trawler velocity in current tick | trawler-select-velocity; trawler-docked | |
| | | day_of_next_trip | Julian day | day trawler will leave port on next fishing trip | trawler-select-day-of-next-trip | Small trawlers: random-uniform 1-3 days Medium trawlers: random-uniform 3-5 days |
| | | trip_duration | hours | total time out of port | trawler-select-trip-duration | Small trawlers: random-normal, μ = 11 hours, σ = 0.25 hour; Medium trawlers: random-normal, μ = 96 hours, σ = 10 hours |
| | | remaining_trip_duration | hours | time remaining in current trip | trawler-select-trip-duration; trawler-reduce-remaining-trip-duration | |
| | | desired_trawl_length_km | km | desired length of trawl; could be | trawler-select-new-trawl-specs; | Small trawlers: Random- |

| | | | | | | |
|---|---|---|---|---|---|---|
| | | | | interrupted by land or protected patch | trawler-fishing-edge-constraint; trawler-fishing-no-constraint; trawler-decide-what's-next-fishing-incomplete; trawler-update-desired-trawl-length-attribute | gamma distribution; L = 5 + random-gamma 1.5 0.125 Medium trawlers: Random-gamma distribution; L = 10 + random-gamma 1.5 0.125 |
| | | actual_trawl_length_km | km | actual trawl length | trawler-decide-what's-next-fishing-incomplete; trawler-fishing-edge-constraint; trawler-fishing-no-constraint; trawler-update-actual-trawl-length; trawler-docked; trawler-select-new-trawl-specs | |
| | | target_fishing_patch | patch ID | patch where trawler is going to fish | trawler-fishing-no-constraint; trawler-docked; trawler-decide-what's-next-fishing- | |

| | | | | | | |
|---|---|---|---|---|---|---|
| | | | | | incomplete; small-trawler-select-target-fishing-patch; medium-trawler-select-target-fishing-patch | |
| | | fishing_complete? | boolean | true indicates fishing is complete and trawler must return to port; false indicates trawler can continue to fish | trawler-wake-up; trawler-decide-what's-next-fishing-incomplete; | True or false |
| | | trawl_start_coordinates | coordinates | location where trawl started | trawler-fishing; trawler-closeout-trawl | |
| | | trawl_start_tick | tick | time when trawl started | trawler-fishing; trawler-closeout-trawl | |
| | | trawl_start_depth | meters | depth when trawler started trawling | trawler-fishing; trawler-closeout-trawl | |
| | | total_dolphins_flocked_with_me | integer | count of dolphins that flocked with trawler during trawl | dolphin-update-flocking-time-data | |
| | | total_trawl_prep_time_hrs | hours | total time spent in-between trawls preparing for next trawl | trawler-prep-for-next-trawl | |
| | | total_trawl_preps | integer | total count of trawl preparations (hauling net, | trawler-prep-for-next-trawl | |

| | | | | | | |
|---|---|---|---|---|---|---|
| | | | | removing fish, cleaning net, etc.) | | |
| | | avg_trawl_prep_time_minutes | hours | average time spent preparing for trawls | trawler-calculate-average-trawl-prep-time | |
| | | fishing_days_this_year | days | count of days spent fishing; one trawl any time during a day counts as one fishing-day | trawler-increment-fishing-days-this-year; trawler-reset-fishing-days-this-year | |
| | | last_fishing_day | Julian day | last day trawler fished | trawler-increment-fishing-days-this-year | |
| | | total_dolphins_flocking_with_me_list | count | number of total individual dolphins flocking with trawler during each trawl | trawler-count-dolphins-flocking-with-me | |
| | | total_dolphin_groups_flocking_with_me_list | count | number of total dolphin groups flocking with trawler during each trawl | trawler-count-dolphins-flocking-with-me | |
| Patches | Static | elevation | meters | height above sea level | setup-world | |
| | | depth | meters | ocean depth | setup-world | |
| | | trawler_protected_2008 | boolean | whether patch was protected from trawling in 2008 until next protection update or not | update-protection | True or false |
| | | trawler_protected_2020 | boolean | whether patch was protected from trawling in 2020-present or not | update-protection | True or false |
| | | gillnet_protected_1998 | boolean | whether patch was protected from | update-protection | True or false |

| | | | | | | |
|---|---|---|---|---|---|---|
| | | | | gillnets in 1998 until next protection update or not | | |
| | | gillnet_protected_2003 | boolean | whether patch was protected from gillnets in 2003 until next protection update or not | update-protection | True or false |
| | | gillnet_protected_2008 | boolean | whether patch was protected from gillnets in 2008 until next protection update or not | update-protection | True or false |
| | | gillnet_protected_2013 | boolean | whether patch was protected from gillnets in 2013 until next protection update or not | update-protection | True or false |
| | | gillnet_protected_2020 | boolean | whether patch was protected from gillnets in 2020 or not | update-protection | True or false |
| | | gillnet_protected_IUCN_plus | boolean | whether patch was protected from gillnets in IUCN + protection scheme | update-protection | True or false |
| | | is_shoreline? | boolean | whether land patch is adjacent to ocean | setup-world | True or false |
| | | distance_to_shore | patches | distance from ocean patch to shore patch | setup-world | |
| | | dolphin_gradient | n/a | heatmap of dolphin distribution based on population surveys | setup-world | range 0-1 |
| | | land? | boolean | land patch or not | setup-world; cleanup-lake-patches | True or false |
| | | lake? | boolean | indicates inland body of water | setup-world; cleanup-lake-patches | True or false |
| | | patch_area | string | placement area that this patch belongs to | setup-world | e.g., “north-east” |

| | | | | | | |
|---|---|---|---|---|---|---|
| | | patch_port | port ID | if patch has a port in it, identifies the port | setup-ports | e.g., port 3 |
| | | patch_port_name | string | if patch has a port in it, identifies the port name e.g., "Akaroa" | setup-ports | e.g., "Akaroa" |
| | | runway_numbers_list | list of headings | list of two numbers that approximate the shoreline heading at this patch (see text for description) for shoreline patches and ocean patches on 50m and 100m depth contours | setup-shoreline-patches-runway-numbers-list | Range 0-359 |
| | | hr_orientation | string | indicates home range orientation for any dolphin that picks it as home range midpoint shoreline patch | setup-shoreline-patches-home-range-orientation | "N-S" (North-South orientation) or "E-W" (East-West) |
| | Dynamic | trawler_protected | boolean | whether patch is currently protected from trawling or not | update-protection | True or false |
| | | gillnet_protected | boolean | whether patch is currently protected from gillnets or not | update-protection | |
| | | gillnet_gradient | n/a | heatmap of gillnet fishing effort, based on MPI effort data; higher number means more gillnet soak time in patch | setup-lake-patches, update-gillnet-gradient | range 0-1 |
| | | trawler_gradient | n/a | heatmap of trawl fishing effort, based on MPI effort data; higher number means more trawling | setup-lake-patches, update-trawler-gradient | range 0-1 |
| | | temporary_gradient | n/a | relative value used when trawlers & gillnets are selecting most attractive patch | trawler-select-next-fishing-patch-normalized- | |

| | | | | | |
|---|---|---|---|---|---|
| | | | | | random; gillnet-select-next-fishing-patch-normalized-random |

#### 3. Process Overview and Scheduling

*Processes*: Figure B.1 (parts 1 and 2) is a top-level flowchart showing an overview of the model processes that run every time step (hour). The number of trawlers and gillnets in the simulation are adjusted on January 1. New patch protection schemes are introduced at the start of the fishing year on October 1. At startup, dolphins are placed in their home range and offshore distribution (i.e., alongshore distribution and distance from shore) that matches population survey data. The simulation terminates when it has run for the desired number of years.

**Trawlers** leave port when scheduled to do so (Figure B.1.1), deploy their nets to fish (which attracts dolphins and can cause bycatch), and return to port at the end of their trip. Because trawlers take multi-hour fishing trips, no trawler performs every procedure in Figure B.1.1 every hour. Each trawler has unique parameters for when it leaves port, trip duration, length of trawls, residence time in port between trips, velocity, etc. They remain in their home port until it is time to leave on a fishing trip. Then they select the duration of the fishing trip, velocity, and destination, with areas that have historically high fishing effort more likely to be selected. Medium trawlers (16-25m long) take multi-day fishing trips and cannot fish in patches protected from trawling, while small trawlers (< 15m) only take day trips and can fish anywhere. They travel to their selected fishing spot and pick a trawl length (km) and trawl direction which tends to follow the shoreline or a depth contour. Average trawl speed is 2.2 knots for small and 4 knots for medium trawlers. Average travel speed is 8 knots for small and 9 knots for medium trawlers. As they trawl, they attract dolphins within detection range while their nets are in the water. The detection of trawlers by dolphins is determined by trawler noise (150 dB for small and 171 dB for medium trawlers; Daly and White 2021) and the attenuation of trawler noise with distance (see below for more detail). After each completed trawl, the model records any dolphin bycatch and the trawler evaluates whether it continues fishing or returns to port based on remaining trip length and projected travel time. Those that keep fishing can either stay in their current location and start a new trawl, or select a different fishing area before they resume fishing. Trawlers ultimately return to port to

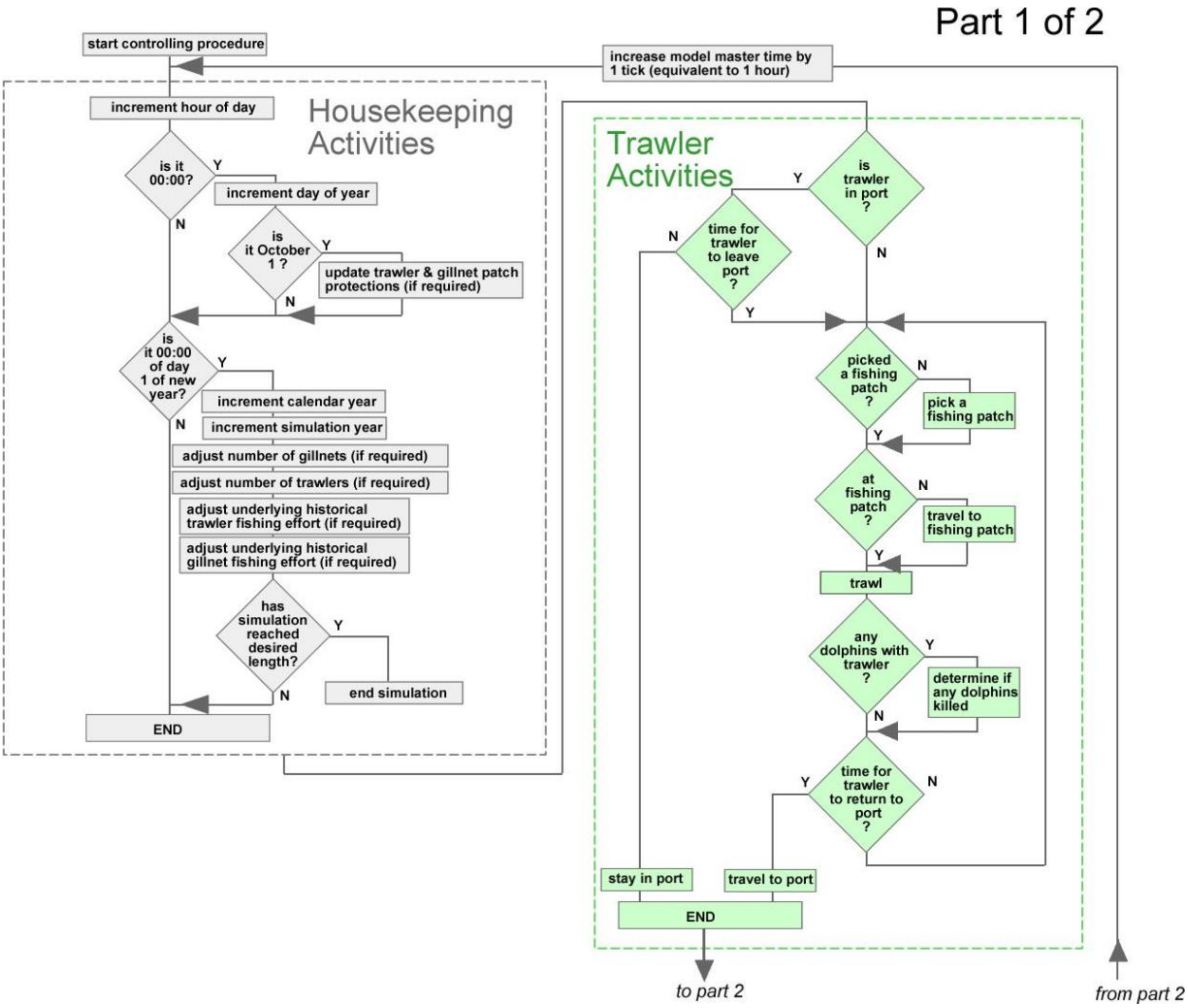

**Figure B.1.1** Main Controlling Procedure Flowchart part 1 (the gray section is run by the model as part of its 'housekeeping' obligations, the green section contains trawler activities).

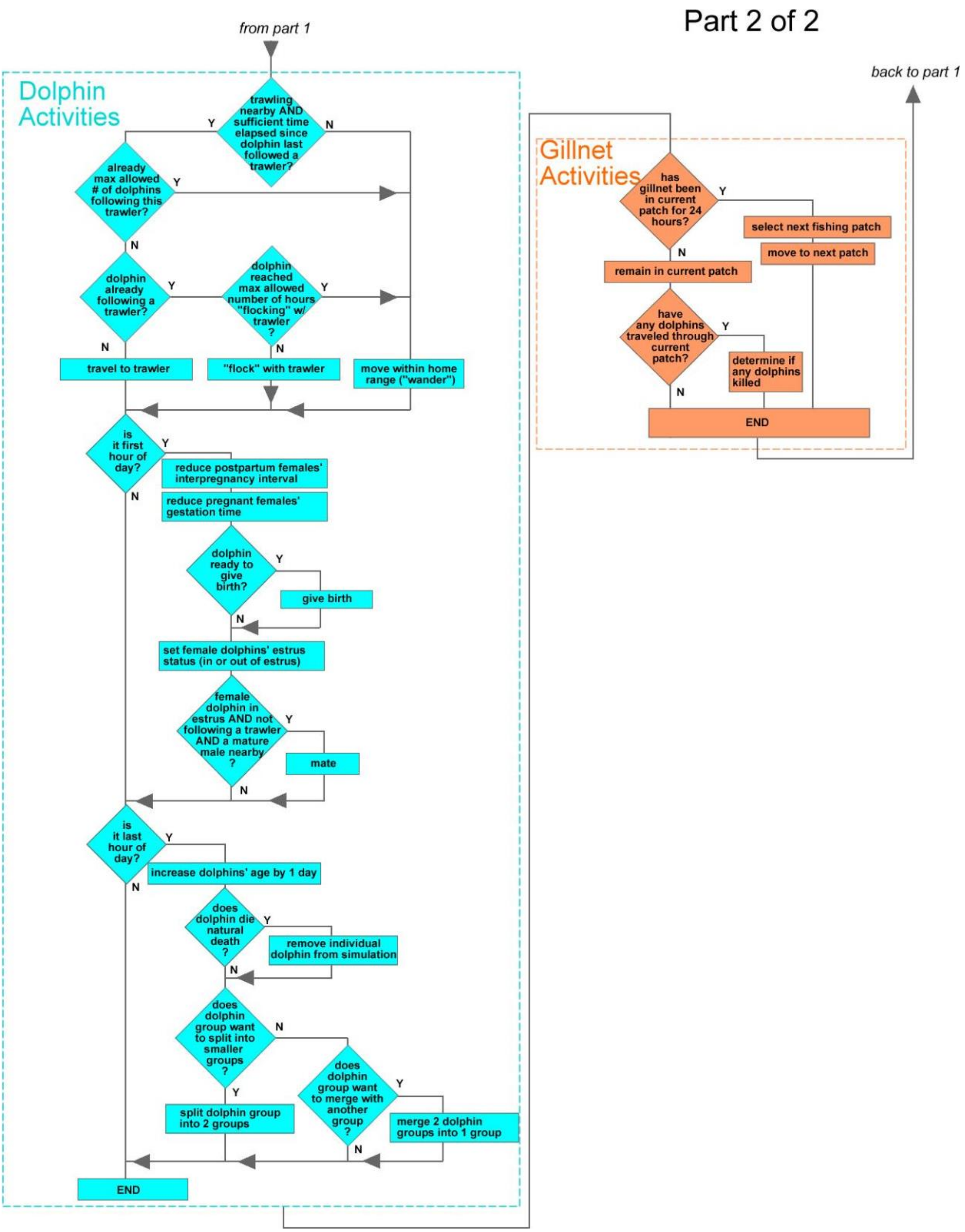

**Figure B.1.2** Main Controlling Procedure Flowchart part 2 (blue section are dolphin activities, and orange are gillnet steps).

conclude their trip, and remain there until it is time to depart on their next trip (intervals based on historical data and are longer for medium trawlers). Because there can be multiple trawlers in a simulation, a single hour in the simulation could find some trawlers in port, while others are traveling to a fishing patch, fishing (potentially generating dolphin bycatch) or traveling home.

**Dolphins** decide on their next activity and velocity based on: whether there is trawling nearby, whether they detect it, whether they are already flocking with the trawler, or if not flocking, whether they choose to travel to it. "Flocking" dolphins travel adjacent (< 10m) to trawlers that are actually fishing and are susceptible to death from bycatch. If not travelling to or flocking with a trawler, dolphins remain in their home range and "wander", which is a catch-all term for time spent in non-trawler focused movement and behavior. Home ranges are individual to each dolphin, and are assigned at model start-up based on field data for observed Hector's dolphin home ranges. Dolphins "wander" via a vector-based random walk that keeps dolphins within their home range and within the maximum allowed water depth. Each time a dolphin selects a new wandering heading, it uses its current heading plus a random normal value with mean zero and a Standard Deviation (wiggle). This initial, unbiased heading is then modified, based on the depth bias parameter (k). The strength of the dolphins' response to water depth is controlled by a logistic curve, with k determining the steepness of the curve. With high values of k, dolphins only respond when they are close to the maximum water depth (100m) and then respond strongly by changing their direction away from deep water. At low values of k, dolphins respond more gradually, turning away from deeper water when still in relatively shallow water and showing a relatively weaker response in terms of turning angle.

Dolphins reproduce on a mating/gestation/calving schedule that mimics field data and age-at-maturity constraints. The model includes an Allee effect which allows mature females to become pregnant only if there is a mature male within 5km. This causes the reproductive rate to be lower in when population densities are low. Field data from different New Zealand dolphin populations indicate that the proportion of calves is lower in smaller populations. The

5km range used in our model results in similar differences in the proportion of calves among populations of different sizes. Dolphins adjust group size by merging into larger groups and splitting into smaller groups ("fission-fusion" behavior), resulting in the same group sizes observed in the field. Their probability of dying from old age increases as they approach the maximum age of 30 years.

**Gillnets** stay in the same patch for a 24-hour "soak time" where they might be encountered by dolphins, potentially resulting in bycatch.  When soak time is complete, they relocate to another patch to soak for 24 hours.

*Scheduling*: The order of procedures is the same for each time step, however not all procedures run every time step; e.g., dolphin age is incremented only at the end of every day. The order in which specific dolphins, trawlers, and gillnets execute their steps is random for each time step, and their dynamic state variables are updated as soon as a new value is calculated.

#### 4. Design Concepts

##### Basic principles

This model addresses the effects of trawler and gillnet bycatch on the population dynamics of Hector's and Māui dolphins.  The model builds on the notion that bycatch probabilities are related to the spatial and temporal overlap in the presence of dolphin and fishing gear, but takes into account that dolphins are attracted to actively fishing trawlers and may remain by trawlers even in areas where dolphins are scarce.  The model incorporates positive density dependence at low population sizes, which provides a degree or realism that cannot be incorporated in non-dynamic models.  The model is comparable with other agent-based models that incorporate marine mammal bycatch as population stressors (Cervin et al. 2020; Gende et al. 2018; Nabe-Nielsen et al. 2014; van Beest et al. 2017).  The dolphin, trawler, and gillnet behaviors are customized to reflect field data on dolphins and historical fishing effort in New Zealand waters.  The model incorporates historical and current trawler-protected and gillnet-protected areas so it can quantify the effect of protection zones in conservation efforts.

##### Emergence

Trawler fishing patterns emerge from the number and type of trawlers, length of fishing trips, and individual trawler decisions on fishing locations, trawl lengths, and whether the trawlers encounter land or a protected patch during a trawl. Gillnet soak time patterns emerge from their placement and soak time. Gillnet mortality patterns emerge from the interaction of dolphin movement patterns and gillnet quantities, soak time, and location. Dolphin movement patterns while “wandering” emerge from their velocity, home range shoreline length, maximum depth limit, turning angles, and how strongly dolphins react as they approach the limits of their home ranges and maximum depth. Dolphin movement patterns near trawlers emerge from the number and location of trawlers, length of trawls, and the density of dolphins within the distance trawlers can be detected. Trawler bycatch mortality emerges from the dolphins’ attraction to trawlers, which is influenced by variations in both vessel and dolphin densities in space and time. The annual dolphin population pattern emerges from initial population size and demographics, births, and deaths via bycatch and natural causes (age).

##### Adaptation

Trawlers adapt their fishing decisions based on their size, location, travel limits, water depth constraints, and historical fishing effort. Medium trawlers adapt to changes in patch protection schemes; medium trawlers will not start a trawl in a protected patch, and if they encounter one mid-trawl, will end the trawl rather than proceed. Small trawlers do this but only under the “IUCN” and “IUCN Plus” protection schemes; otherwise, they ignore protected patches. Dolphins adapt to changed trawler densities; more trawlers mean more opportunities to flock and take fish. This also increases dolphin-trawler encounters and bycatch mortality.

#### Objectives

The objective of the trawlers and gillnets is to fish at locations, durations, and effort levels that match historical data. Therefore, there is no direct "payoff" to evaluate before each trawler or gillnet decision. For example, trawler trip duration and trawl length decisions result from random draws within a distribution based on the trawler size (small or medium). Trawlers select their fishing patch based on how far they are willing to travel (estimated from field data) and a weighted random draw derived from historical fishing effort data contained within each patch; gillnets also select their fishing patches from a weighted random draw based on patch-specific historical gillnet effort data. The dolphins have two objectives: move within their home range and flock with an actively fishing trawler when one is detected under the imposed constraints. Dolphins cannot always flock with a trawler or stay indefinitely since there are constraints that limit the number of dolphins with a trawler, the amount of time a dolphin can stay with a trawler, and an interval timer that prevents the dolphin from immediately moving to a different trawler nearby.

#### Learning

Learning is not implemented.

#### Prediction

None of the entities use implicit or explicit prediction to drive adaptive behaviors or estimates of future conditions.

#### Sensing

Female dolphins in estrus sense mature male dolphins of the same subspecies within dolphin detection range. Dolphins sense other dolphins of the same subspecies nearby which initiates a potential group merge. Dolphins sense when they are adjacent to another placement area; this triggers the dolphin to decide whether to switch its placement area and home range. Dolphins sense trawlers when they are actively trawling based on emitted trawl noise, transmission losses, and detection probability (see dolphins-detect-trawling-nearby-and-decide-to-travel in section 7). Dolphins that attempt to flock with a trawler sense how many dolphins are already there and if joining those already there would exceed the maximum limit of dolphins with a single trawler.  Dolphins sense when they pass through a patch with a gillnet.  Dolphins sense and respond to the limits of their home range, maximum depth limit and avoid land patches.  Trawlers sense how many dolphins are flocking with them when they complete a trawl to compute bycatch.  Trawlers sense and avoid land. Trawlers sense the orientation of shoreline patches and ocean patches on the 50m and 100m depth contour and use them to select a trawl heading.  Medium trawlers sense and avoid protected patches; gillnets do likewise for gillnet-protected patches.  Trawlers and gillnets base their fishing location decisions on depth and historical fishing effort data contained within each ocean patch.

##### Interaction

Dolphins interact with others of the same subspecies to mate and occasionally merge groups.  Dolphins interact with trawlers and gillnets resulting in bycatch deaths.

##### Stochasticity

Most of the procedures controlling agent movements and decisions are stochastic. Distributions and probabilities were selected based on field data (dolphins) and industry data (trawlers and gillnets). Model-generated patterns were compared to corresponding field and industry data via Pattern Oriented Modeling (Wiegand et al. 2003; Grimm and Railsback 2012; Railsback and Grimm 2019).

For dolphins, stochastic processes include their initial placement, group size, ages, genders, home range dimensions, fraction of females pregnant, and fraction of females still within their post-partum interval.  Female dolphins randomly select their estrus start days, probability of successful mating, length of gestation, and length of post-partum interval. Dolphins randomly select their initial heading when not pursuing a trawler.  The gender of calves is randomly selected.  Stochastic processes also control number of dolphins splitting off to form a new group and migrating to new placement area and selecting new home range dimensions.

Trawlers are initialized with a randomly assigned boat length and hour of day when they nominally leave port to start a trip. Trawlers use stochastic processes to: select the day and duration of its next trip, the next fishing patch to start a trawl, the heading to take during the trawl, and the length of the trawl (km).

Gillnets randomly select their next fishing patch.

#### Collectives

Dolphins are the only collectives in the model; each dolphin represents multiple individual dolphins and has its own state variables. It was more efficient to model dolphins as groups rather than thousands of individual dolphins and force collective movement (e.g.; moving cohesively, flocking with trawler, etc.) and decisions.

#### Observation

The model offers multiple ways to observe, save, and visualize data. Trawler, dolphin, and gillnet movement can be observed on the user interface, including tracing movement if desired. There are histograms of multiple entities' state variables to check if model distributions match distributions based on field data.

The following external data files can be populated during a simulation (user choice) for post-processing/analysis/visualization:

- dolphin demographics – every month the model records: number of dolphin individuals by subspecies, five-year age bucket and gender, and average group size
- dolphin location – any time user asks, model exports: dolphin with unique identifying id number, subspecies, group size, coordinates, latitude, longitude, distance from shore (nmi), and patch depth (m)
- age of females when they give birth – each time a female delivers a calf, model records: model revision, simulation run, day of year, year, subspecies, and mother's age

- trawl data -  each time a trawler completes a trawl it records: model revision, simulation run, day of year, year, trawler, home port, trawler size (small or medium), trawler length (m), trawl start coordinates, trawl end coordinates, trawl start latitude and longitude, trawl end latitude and longitude, trawl length, trawl start time, trawl end time, trawl duration, trawl start depth, trawl end depth, total trawls for this trawler, total trawling hours for this trawler, trawling velocity, and any comment to clarify why a trawl ended; e.g., "return to port"
- fishing days data – at midnight on the first day of each year, each trawler records: model revision, simulation run, day of year, year, trawler, home port, trawler size (small or medium), trawler length (m), and total fishing days from the previous year; these data are used to compare fishing effort with government data.
- trawler bycatch data - every time a dolphin dies in a trawl, we record the following: model revision, simulation run, day of year, year, trawler, home port, trawler size, trawler length, coordinates, latitude, longitude, total dolphins flocking, number of Hector's dolphins killed, and number of Maui dolphins killed
- gillnet bycatch data - every time a dolphin dies in a gillnet, we record the following: model revision, simulation run, day of year, year, gillnet, gillnet area, coordinates, latitude, longitude, patch depth, total dolphins in the patch, number of Hector's dolphins killed, and number of Maui dolphins killed

As the model runs, it also records data within the patches that support observations of entity behaviors.  They can be visualized immediately on the user interface as heat maps, or can be exported to csv files.  They are:

- dolphin wandering – each ocean patch tracks how many hours dolphins spend wandering there
- dolphin flocking with trawler – each ocean patch tracks the total time dolphins spend flocking with trawlers in units of dolphin-hours; e.g., 10 dolphins flocking for 1 hour = 10 dolphin-hours

- dolphin births – each ocean patch tracks total calves born there
- dolphin-trawler bycatch – each ocean patch tracks the number of dolphins killed by trawlers
- gillnet soak time – each ocean patch tracks total hours gillnets have been there
- gillnet-dolphin encounters – each gillnet keeps a running total of dolphin groups that have been in the same patch
- dolphin-gillnet bycatch – each ocean patch tracks the total dolphins killed by gillnets
- trawl start – each ocean patch tracks the total number of trawls that start in the patch

#### 5. Initialization

Figure B.2 shows the setup procedure and sub-procedures that it initiates. Setup runs in full to initialize a simulation for the first time; subsequent simulations can initialize with only a subset of procedures as desired (see Section 7 for detailed description of setup and sub-procedures).

**Time and day:** The simulation begins at midnight on January 1 of the desired initial calendar year (*setup-time*). *Setup-world* does everything involved with getting the environment ready to run and begins by reading in GIS files that determine which patches are land, ocean, or lake. GIS attributes include elevation for land patches and depth for ocean patches. The model checks the initial year and refers to internal data that identifies patches protected from gillnet and/or trawler fishing by year and sets the patch protection attributes. Each ocean patch is loaded with year-based gradient data (number from 0-1) that indicates how likely it is to find a gillnet, dolphin, or trawler fishing in that patch. The model uses that information to place dolphins and gillnets in the ocean at startup; trawlers start out in their home port but use their gradient data to pick fishing patches. The model adds guide numbers to shoreline patches and ocean patches at 50m and 100m depth that trawlers use to help set initial trawl headings (see *setup-world* in Section 7 for details). The remaining initialization steps occur no matter what the geographical selection is; just the number and location of entities differ.

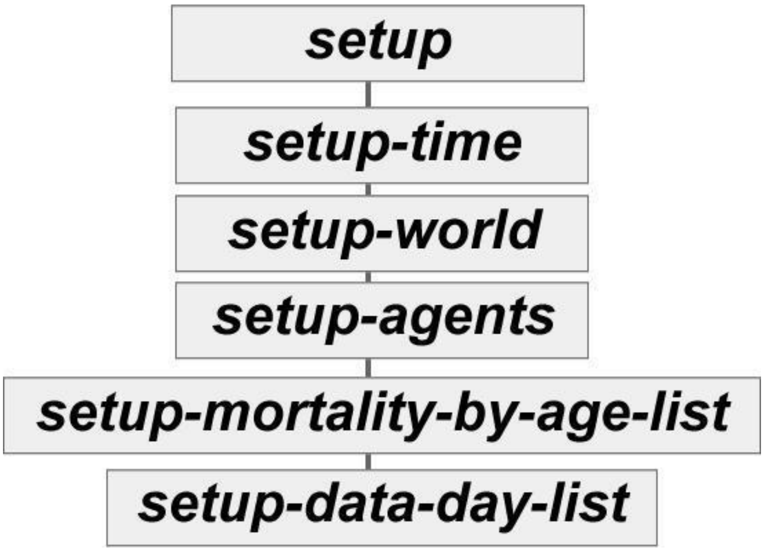

**Figure B.2.** Setup Procedure.

The *setup-agents* procedure initializes the quantity and location of agents including:

- **Ports & Waypoints**: Ports and waypoints are created and placed at specific coordinates (*setup-ports*). They have no processes or submodels.
- **Trawlers**: The model creates an initial number of trawlers based on the simulation start year and geographic area. See Section 7 for the *setup-trawlers* procedure.
- **Gillnets**: The model creates an initial number of gillnets based on the simulation start year and geographic area. See section 7 for the *setup-gillnets* procedure.
- **Dolphins**: The model creates and positions an initial number of Maui and Hector's dolphins based on UI settings. See *setup-dolphins* details in Section 7.

The *setup-mortality-by-age-list* procedure builds the list of 31 age-based dolphin mortality probabilities and *setup-data-day-lists* build a list of 12 Julian days, one for the last day of each month that the model uses to determine when to export monthly data.

#### Globals Table

Table B.2 shows the global variables, which entities they affect, initial values, and the data source. Note that most of these are on the User Interface (UI).

**Table B.2 Global Variables**

| Scope | Global Variable | Meaning | Value | Data Source |
|---|---|---|---|---|
| Generic | protection (UI) | trawler and gillnet patch protection scheme | 2020. Other options are 1988, 2003, 2008, 2013, "none", "IUCN", "IUCN Plus", and "year" | Historical data for protection schemes by effective year (Dawson and Slooten 1993; Gormley et al. 2012; Slooten 2013; MPI 2023 shapefiles for current protected areas) |
| | input_starting_year (UI) | calendar year to start simulation if not using current calendar year | n/a | n/a |
| Dolphins | initial_hectors_banks_peninsula (UI) | initial dolphin population in this area | 6953 | MacKenzie and Clement 2014, 2016, 2019 |
| | initial_hectors_south_west (UI) | initial dolphin population in this area | 5807 | MacKenzie and Clement 2014, 2016, 2019 |
| | initial_hectors_south_north (UI) | initial dolphin population in this area | 2146 | MacKenzie and Clement 2014, 2016, 2019 |
| | initial_hectors_south_east (UI) | initial dolphin population in this area | 384 | MacKenzie and Clement 2014, 2016, 2019 |
| | initial_māui (UI) | initial dolphin population in this area | 73 | Cooke et al., 2019; IWC 2023 |
| | hector_mating_range (UI) | range where female dolphins that are ready to mate can detect eligible males | 5 km | Slooten 1991; Slooten and Lad 1991; Slooten et al. 1993; Turek et al. 2013; Harvey 2021; Constantine et al. 2021; Williams 2022; Bennington 2025 |
| | maui_mating_range (UI) | range where female dolphins that are ready to mate can detect eligible males | 5 km | Slooten 1991; Slooten and Lad 1991; Slooten et al. 1993; Turek et al. 2013; |

| | | | | |
|---|---|---|---|---|
| | | | | Harvey 2021; Constantine et al. 2021; Williams 2022; Bennington 2025 |
| | reproduction_probability (UI) | odds that mating event results in pregnancy | 0.02 | Slooten 1991; Slooten and Lad 1991; Gormley 2009 |
| | fraction_dolphins_migrating_per_year (UI) | fraction of dolphin groups that migrate to new area | 0 | Fletcher et al. (2002) estimated migrating fraction, but for this analysis it was set to 0 |
| | dolphin_max_velocity (UI) | maximum dolphin swimming velocity | 12 km/hr | Slooten 1994 and additional field observations; see histogram in *dolphin-select-velocity* above. |
| | dolphin_wandering_wiggle (UI) | maximum change in heading when dolphin selects next heading while "wandering" | 20 deg | Calibration |
| | max_dolphin_depth (UI) | dolphins avoid patches at this depth and greater | 100 m | Slooten et al. 2006b; MacKenzie and Clement 2014, 2016, 2019 |
| | home_patch_min_offshore_distance (UI) | minimum offshore distance for dolphin home patch | 0 km | Calibration |
| | home_patch_max_offshore_distance (UI) | maximum offshore distance for dolphin home patch | 5 km | Calibration |
| | allow_dolphin_switches? (UI) | allows dolphins to switch areas or not | off | n/a |
| | mean_home_range_shoreline_length | mean shoreline length of dolphin home range | 50 km | Rayment et al. 2009 |
| | std_dev_home_range_shoreline_length | standard deviation of dolphin home range shoreline length | 2.5 km | Rayment et al. 2009 |

| | | | | |
|---|---|---|---|---|
| | max_dolphins_allowed_to_flock_with_trawler (UI) | maximum number of individual dolphins allowed to flock with trawler at once | 200 | Calibration |
| | allow_dolphin_births? (UI) | allows dolphins to give birth or not | off | n/a |
| | allow_pre_sim_dolphin_pregnancy? (UI) | determines whether some dolphins are initialized as pregnant and some unable to get pregnant because they are waiting for their postpartum interval to expire | off | n/a |
| | allow_dolphin_splits_and_merges? (UI) | allows dolphin groups to split into smaller groups or merge with another group | off | n/a |
| | dolphin_min_interval_between_flocking (UI) | minimum time dolphin group has to wait after a flocking session before it can flock with a trawler again | 6 hrs | field data and calibration |
| | dolphin_max_interval_between_flocking (UI) | maximum time dolphin group has to wait after a flocking session before it can flock with a trawler again | 10 hrs | field data and calibration |
| | dolphin_min_flocking_time (UI) | minimum time dolphin group spends flocking with a trawler | 3 hrs | field data and calibration |
| | dolphin_max_flocking_time (UI) | maximum time dolphin group spends flocking with a trawler | 5 hrs | field data and calibration |
| | hectors_min_merge_prob (UI) | minimum probability that a Hector's dolphin group will merge with another Hector's group | 0.4 | field data and calibration |
| | hectors_merge_prob_L (UI) | factor that controls merge probability; as potential merged group size increases, probability of merging decreases | 0.1 | field data and calibration |
| | hectors_min_split_prob (UI) | minimum probability that a Hector's dolphin group will split into two groups | 0.2 | field data and calibration |

| | | | | |
|---|---|---|---|---|
| | hectors_split_prob_L (UI) | factor that controls split probability; as group size increases, probability of splitting increases | 0.1 | field data and calibration |
| | maui_min_merge_prob (UI) | minimum probability that a Maui dolphin group will merge with another Maui group | 0.8 | field data and calibration |
| | maui_merge_prob_L (UI) | factor that controls merge probability; as potential merged group size increases, probability of merging decreases | 0.1 | field data and calibration |
| | maui_min_split_prob (UI) | minimum probability that a Hector's dolphin group will split into two groups | 0.0001 | field data and calibration |
| | maui_split_prob_L (UI) | factor that controls split probability; as group size increases, probability of splitting increases | 0.2 | field data and calibration |
| Trawlers | trawler_fishing_patch_selection_method (UI) | controls how trawlers select fishing patches | Used "historical probability"; other options are "random" and "normalized random" | derived from a heat map of fishing effort provided by MPI (2023, 2024) |
| | trawler_kill_probability (UI) | probability that an individual dolphin dies if it is flocking with a trawler when a trawl finishes | 1.1e-4 | Calibration based on MPI estimate of total dolphins caught per year |
| | small_trawler_max_initial_travel_time_home_port (UI) | maximum travel time from home port to initial fishing patch | 3.5 hrs | Calibration (AIS fishing effort data from GFW 2025, direct observations and interviews with fishers) |
| | small_trawler_max_travel_time_next_patch (UI) | maximum travel time to next fishing patch if trawler decides to relocate | 2 hrs | Calibration (AIS fishing effort data from GFW 2025, direct |

| | | | | |
|---|---|---|---|---|
| | | | | observations and interviews with fishers) |
| | small_trawler_max_start_trawl_depth (UI) | maximum patch depth to start a trawl | 30 m | Fishing effort data from MPI 2023, 2024 and GFW 2025 |
| | small_trawler_switch_trawl_patch_odds (UI) | probability a trawler will relocate to another patch after completing a trawl | 0.5 | Calibration (AIS fishing effort data from GFW, 2025, direct observations and interviews with fishers) |
| | medium_trawler_max_initial_travel_time_home_port (UI) | maximum travel time from home port to initial fishing patch | 6 hrs | Calibration (AIS fishing effort data from GFW 2025, direct observations and interviews with fishers) |
| | medium_trawler_max_travel_time_next_patch (UI) | maximum travel time to next fishing patch if trawler decides to relocate | 4.5 hrs | Calibration (AIS fishing effort data from GFW 2025, direct observations and interviews with fishers) |
| | medium_trawler_min_start_trawl_depth (UI) | minimum patch depth to start a trawl | 20 m | Fishing effort data from MPI 2023, 2024 and GFW 2025 |
| | medium_trawler_max_start_trawl_depth (UI) | maximum patch depth to start a trawl | 100 m | Fishing effort data from MPI 2023, 2024 and GFW 2025 |
| | medium_trawler_switch_trawl_patch_odds (UI) | probability a trawler will relocate to another patch after completing a trawl | 0.5 | Calibration (AIS fishing effort data from GFW 2025, direct observations and interviews with fishers) |

| | | | | |
|---|---|---|---|---|
| | small_trawler_mean_trip_duration (UI) | mean trip duration (time out of home port on a fishing trip) | 11 hrs | Calibration (AIS fishing effort data from GFW 2025, direct observations and interviews with fishers) |
| | small_trawler_sd_trip_duration (UI) | standard deviation for fishing trip time | 0.25 hr | Calibration (AIS fishing effort data from GFW 2025, direct observations and interviews with fishers) |
| | med_trawler_mean_trip_duration (UI) | mean trip duration (time out of home port on a fishing trip) | 96 hrs | Calibration (AIS fishing effort data from GFW 2025, direct observations and interviews with fishers) |
| | med_trawler_sd_trip_duration (UI) | standard deviation for fishing trip time | 10 hrs | Calibration (AIS fishing effort data from GFW 2025, direct observations and interviews with fishers) |
| | small_trawler_min_days_between_trips (UI) | minimum number of days in port between fishing trips | 1 day | Calibration (AIS fishing effort data from GFW 2025, direct observations and interviews with fishers) |
| | small_trawler_days_between_trips_alpha (UI) | factor in the random gamma draw that determines when trawler leaves on next trip | 2.0 | Calibration (AIS fishing effort data from GFW 2025, direct observations |

| | | | | |
|---|---|---|---|---|
| | | | | and interviews with fishers) |
| | small_trawler_days_between_trips_lambda (UI) | factor in the random gamma draw that determines when trawler leaves on next trip | 0.5 | Calibration (AIS fishing effort data from GFW 2025, direct observations and interviews with fishers) |
| | med_trawler_min_days_between_trips (UI) | minimum number of days in port between fishing trips | 4 days | Calibration (AIS fishing effort data from GFW 2025, direct observations and interviews with fishers) |
| | med_trawler_days_between_trips_alpha (UI) | factor in the random gamma draw that determines when trawler leaves on next trip | 6.0 | Calibration (AIS fishing effort data from GFW 2025, direct observations and interviews with fishers) |
| | med_trawler_days_between_trips_lambda (UI) | factor in the random gamma draw that determines when trawler leaves on next trip | 0.5 day | Calibration (AIS fishing effort data from GFW 2025, direct observations and interviews with fishers) |
| | sm_trawler_mean_trawl_noise (UI) | mean emitted noise during trawl | 150 dB | Daly & White 2021 |
| | med_trawler_mean_trawl_noise (UI) | mean emitted noise during trawl | 171 dB | Daly & White 2021 |
| | TL_equation_coefficient (UI) | coefficient in spreading transmission loss equation; TL = TL_equation_coefficient * logR; used to calculate received level of trawl noise at dolphin | 15 | Lynch & Newhall 2017 |

| | | | | |
|---|---|---|---|---|
| | alpha_broadband | broadband noise absorption coefficient in seawater; used to calculate trawl noise transmission loss | 1.27e-05 dB/m | Richardson et al. 2013 |
| Gillnets | gillnet_patch_selection_method (UI) | method gillnets use to select fishing patches | Used “historical probability”; other options are “random”, and “normalized random” | derived from a heat map of fishing effort provided by MPI 2023, 2024 |
| | gillnet_kill_probability (UI) | probability of an individual dolphin’s death when dolphin group encounters gillnet | 0.005 | Calibration based on MPI 2023, 2024 estimate of total number of dolphins caught per km of gillnet |
| Patch sets | unprotected_trawler_patches | patches eligible for fishing by trawlers | set based on protection scheme | Dawson and Slooten 1993; Gormley et al. 2012; Slooten 2013; MPI 2023 shapefiles for current protected areas |
| | protected_trawler_patches | patches protected from trawler fishing | set based on protection scheme | Dawson and Slooten 1993; Gormley et al. 2012; Slooten 2013; MPI 2023 shapefiles for current protected areas |
| | unprotected_gillnet_patches | patches eligible for fishing by gillnets | set based on protection scheme | Dawson and Slooten 1993; Gormley et al. 2012; Slooten 2013; MPI 2023 shapefiles for current protected areas |
| | protected_gillnet_patches | patches protected from gillnet fishing | set based on protection scheme | Dawson and Slooten 1993; Gormley et al. |

| | | | | |
|---|---|---|---|---|
| | | | | 2012; Slooten 2013; MPI 2023 shapefiles for current protected areas |
| Patch dimension | km | convert patches to kilometers | 0.613 patch/km | n/a |
| | patch_size | length of sides of each patch | 1.63 km/patch | n/a |
| Distance bands | near_band | area closest to shore where dolphins are located | 0-4 nautical miles offshore | Survey strata from MacKenzie and Clement 2014, 2016, 2019 |
| | middle_band | middle area where dolphins are located | 4-12 nautical miles offshore | Survey strata from MacKenzie and Clement 2014, 2016, 2019 |
| | far_band | area furthest from shore where dolphins are located | 12-20 nautical miles offshore | Survey strata from MacKenzie and Clement 2014, 2016, 2019 |

#### 6. Input Data

The model uses multiple GIS files to setup the world. GIS data identify land, lake, and ocean patches, as well as which ocean patches are protected from gillnet and trawler fishing by year. Patch elevation and depth are included along with the shapes of the placement areas. Dolphin survey data and historical trawling and gillnet effort are read in from three .asc files and used as patch attributes; the trawler and gillnet fishing effort data are used by those entities to measure the relative attractiveness of potential fishing patches. The number, size, and home port of trawlers is initialized and updated annually from an external data file of the number of trawlers by size and port. At the start of every year, it refers to that file and adjusts the number of trawlers as required.

#### 7. Submodels

*7.1 Setup*

Setup prepares the simulation to run per the flowchart in Figure B.3; detailed descriptions of sub-procedures are below.

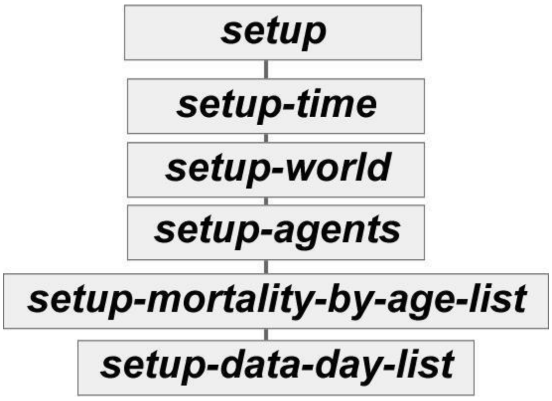

**Figure B.3** Setup Procedure Flowchart.

*7.1.1 setup-time*

Model default start time is January 1 at midnight. This procedure sets the initial time variables: year_of_sim = 1, hour_of_day = 0, day_of_year = 1, day_of_sim = 1. It sets the starting_year (the calendar year we want to start in) based on a UI pulldown. If there is a year entered, it begins in that year; otherwise, it starts in the current calendar year. It sets the current_year (which is the year the model uses and increments every 365 days) to match the starting_year.

*7.1.2 setup-world*

Setup-world manages all the procedures that create the environment or the dolphins, trawlers, and gillnets (Figure B.4). Sub-procedures are indicated with italics and dashes; e.g., setup-shoreline-patches.

*7.1.3 setup-shoreline-patches*

This procedure identifies land patches adjacent to ocean patches and sets the is_shoreline? attribute to true.

*7.1.4 setup-shoreline-patches-home-range-orientation*

Because the shoreline contour varies around the islands and we needed a way to measure the maximum distance a dolphin can travel within its home range as part of the random walk algorithm, we classified the dolphins by home range orientation; "North-South" home ranges for those dolphins with home ranges centered along a stretch of shore that runs predominantly north-south and "East-West" for those with home ranges on shorelines that run mostly east-west. This procedure assigns the "NS" or "EW" to each shoreline patch. When the dolphin picks its home range, it refers to the shoreline patch in the middle of its range and adopts the "NS" or "EW" orientation. See dolphin-setup-home-patch-and-home-range procedure for more details.

*7.1.5 setup-depth-contour-patches*

This procedure identifies which ocean patches are on the 50m and 100m depth contours.

*7.1.6 setup-placement-areas*

The model classifies groups of patches as "placement areas" to simplify some of the entities' initial location assignments and track output data on a larger scale; e.g., trawler kills off the South Island, south of Banks Peninsula area (abbreviated as "south-bankssouth"). All 12 placement areas are shown in Figure B.5.

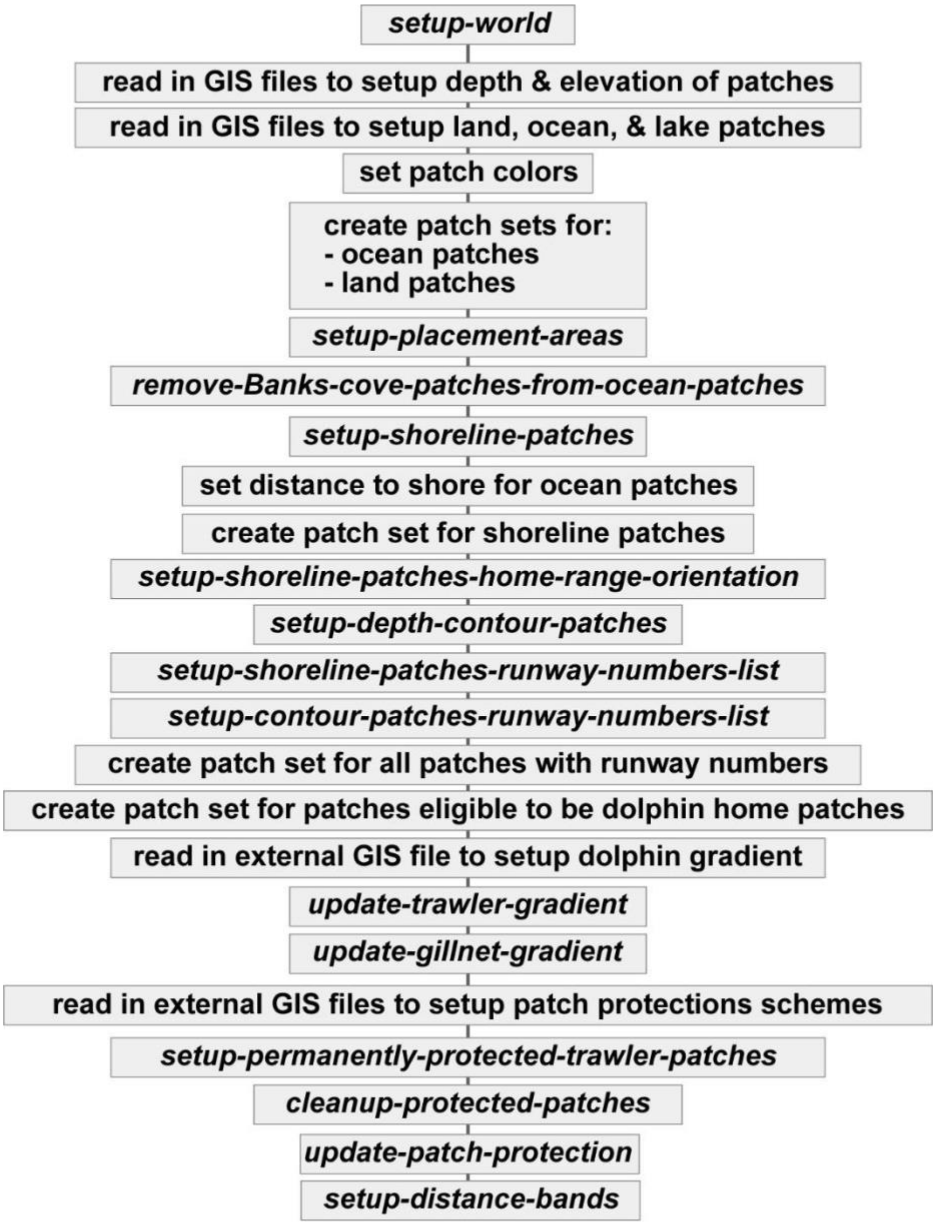

**Figure B.4** setup-world procedure flowchart.

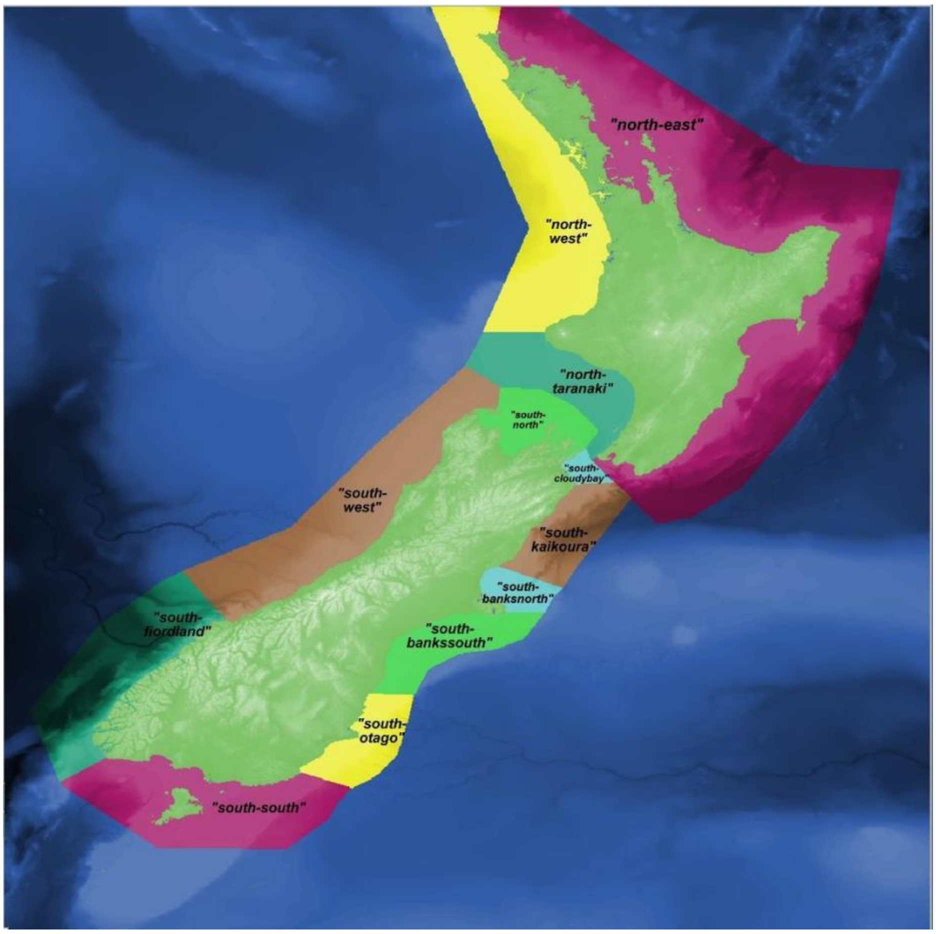

**Figure B.5** Placement areas.

*7.1.7 setup-shoreline-patches-runway-numbers-list*

AIS data and government fishing effort data show trawlers tend to trawl on a heading that roughly parallels depth contours (GFW 2025). Therefore, shoreline patches and ocean patches at 50m and 100m depth have “runway” numbers that trawlers use as references when picking initial trawl headings. Runway numbers indicate a rough estimate of the compass heading of several adjacent shoreline or ocean patches. To establish the runway numbers, the model uses a temporary agent to act as a surveyor that does the following:

1. build a patch-set of acceptable patches (e.g., if model is working on shoreline runway numbers, the acceptable patch set is all shoreline patches; if working the 50m depth contour it picks other patches at 50m depth)
2. start in the center of the patch we are choosing runway numbers for

3. have the surveyor pick the initial heading chosen from 0, 45, 90, 135, 180, 225, 270, or 315 degrees based on which one has the maximum number of acceptable patches that are within a semi-circle in front of the surveyor 10 patches deep
4. the surveyor faces and moves to one of the neighboring acceptable patches (e.g., shoreline patches)
5. surveyor repeats step 4 until it has moved five patches away from the initial patch
6. calculate the initial runway number using *atan* function from the initial patch to surveyor coordinates
7. surveyor moves back to initial patch and chooses its initial heading to go opposite of the heading from step 3
8. repeat the face-neighbor-and-move steps until five patches away from the initial patch
9. calculate second runway number using *atan* function

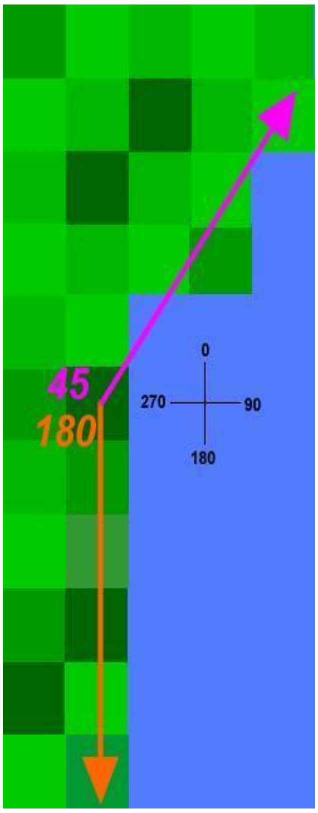

**Figure B.6** Example of patch runway numbers.

Figure B.6 is an example of runway numbers for a shoreline patch. The shoreline patch in dark green with the two arrows has runway numbers of 45 (pink arrow) and 180 (orange arrow). Trawlers choose the runway number closest to their current heading to use as an initial reference when they pick a new trawl heading.

*7.1.8 setup-contour-patches-runway-numbers-list*

The process for 50m and 100m depth ocean patches is the same as described above for shoreline patches, just with a different set of acceptable patches to move through.

*7.1.9 update-trawler-gradient*

Imports the GIS file of trawler effort for each ocean patch based on current year

*7.1.10 update-gillnet-gradient*

Imports the GIS file of gillnet effort for each ocean patch based on current year

*7.1.11 setup-permanently-protected-trawler-patches*

This procedure declares specific ocean patches (e.g., inside harbors) as permanently protected from trawling.

*7.1.12 setup-distance-bands*

The model groups ocean patches by distance from shore. The "near" band is within 4 nautical miles (7.41 km) from shore, the "middle" band is between 4 and 12 nmi (7.42 - 22.22 km) from shore, and the "far" band is between 12 and 20 nmi (22.23 and 37.04 km) from shore as shown in Figure B.7.

*7.1.13 update-patch-protection*

This procedure applies the protections used based on the UI input. Each patch has a true/false attribute that identifies whether or not it is a gillnet or trawler protected patch; e.g., gillnet_protected_2020 and trawler_protected_2020. The number of protected patches increases with later protection schemes as shown in Figure B.8, with "IUCN+" being the most protective (the IUCN and IUCN+ protections are recommended protection levels and have not been implemented in New Zealand).

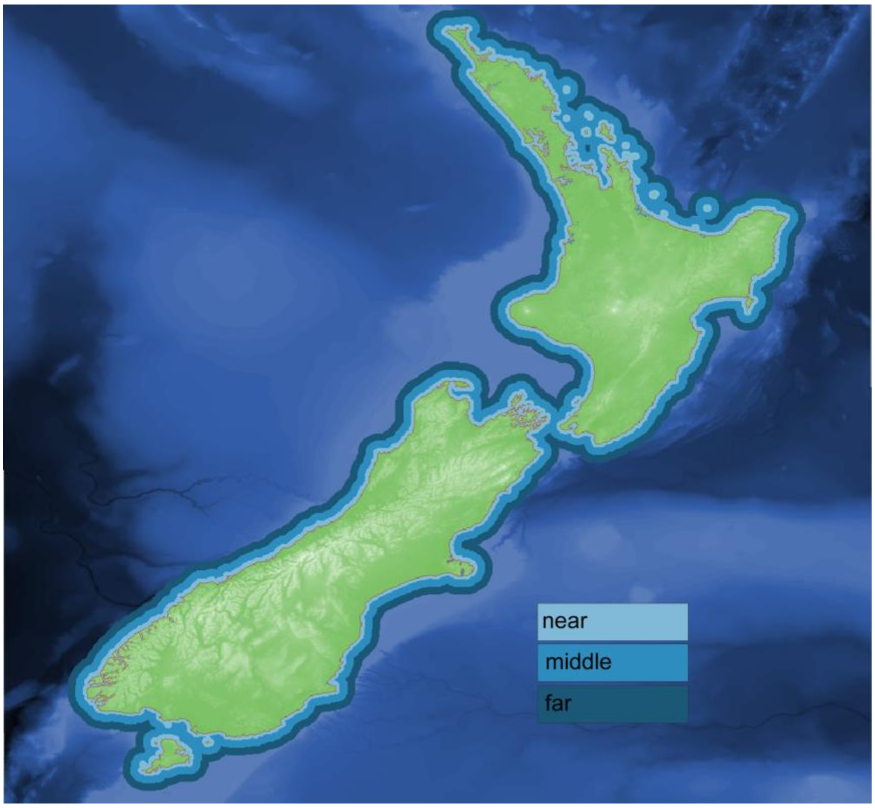

**Figure B.7** Distance bands.

Patch protections can either be constant (e.g., run an entire simulation under the 2013-2019 protection scheme for a certain number of years), or it can be configured so the protection scheme changes as the simulation year changes. For example, if the model started in 2015 and ran for 10 years, it would use the Oct 1, 2013-Sep 30, 2020 protections initially and switch to the 2020 protections on Oct 1 of that year. Since IUCN and IUCN+ are recommended protection levels, they must be manually selected.

*7.1.14 setup-mortality-by-age-list*

This procedure converts the age-based annual probability of an individual dolphin dying to daily probabilities and stores them in a list. The daily mortality probabilities are shown in Figure B.9. Estimates of annual survival are available from research using photographic identification of individual dolphins (Gormley 2009; Gormley et al. 2012; Wickman 2024).

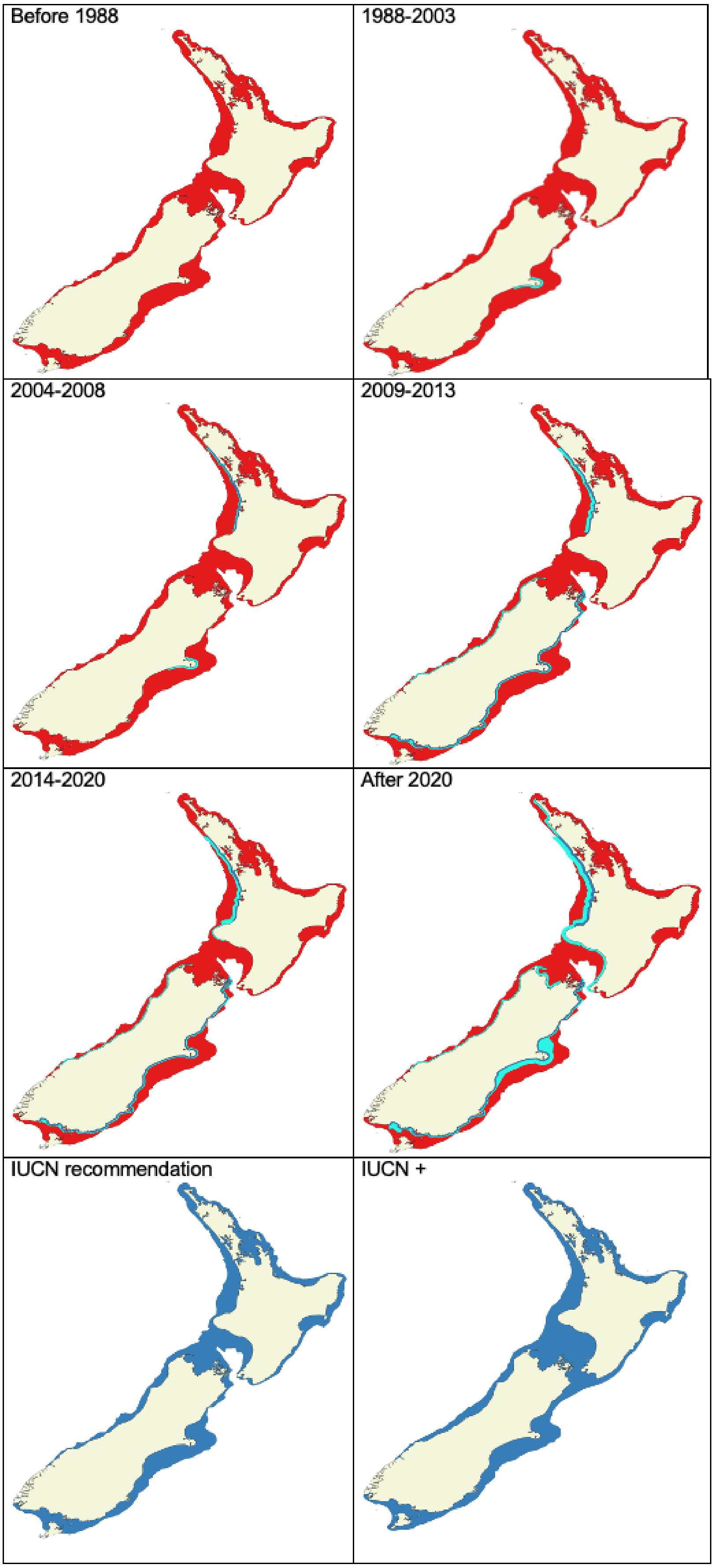

**Figure B.8** Patch protection schemes.

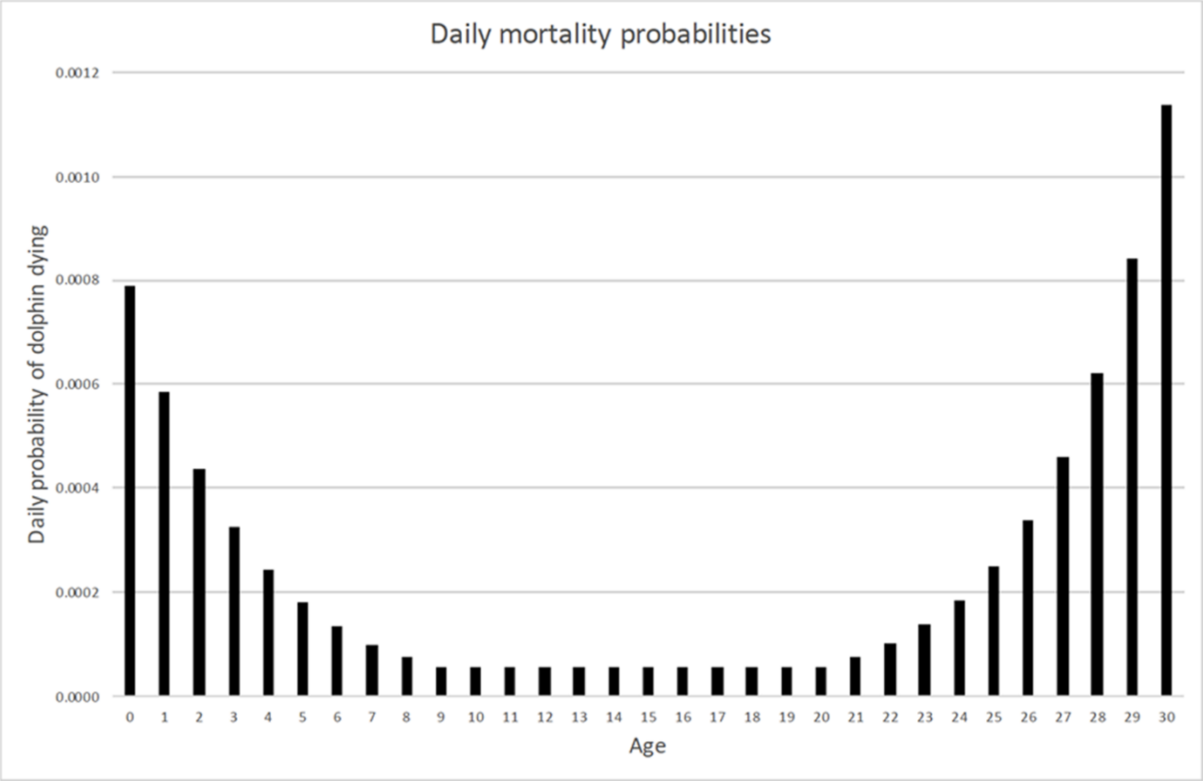

**Figure B.9** Daily probability of dolphin dying depending on age from 0 (newborn to 1 year old) to 30 (maximum age).

*7.1.15 setup-data-day-list*

This procedure creates a list of the Julian last days of the month which trigger exporting dolphin population data:

$$data\_day\_list \quad [31\ 59\ 90\ 120\ 151\ 181\ 212\ 243\ 273\ 304\ 334\ 365]$$

*7.1.16 setup-agents*

Setup-agents is called during initial model setup and any time the operator wants to reset for another simulation without re-loading all the GIS data via a button on the UI. A flowchart setup-agents is shown in Figure B.10.

*7.1.17 check-min-max-sliders*

The UI has paired sliders that set the minimum and maximum limits for some parameters; this procedure checks to make sure the minimum is < the maximum. If not, it displays an error.

*7.1.18 build-port-locations-table*

This procedure builds the global port_locations_table which holds the name of the port (e.g. "akaroa") and the coordinates (e.g., patch 31 -157) for all 32 ports.

*7.1.19 setup-trawl-data-file*

This procedure creates an empty shell file to hold output data

*7.1.20 setup-fishing-days-data-file*

This procedure creates an empty shell file to hold output data

*7.1.21 setup-dolphin-data-files*

This procedure creates an empty shell file to hold output data

*7.1.22 setup-trawler-bycatch-data-file*

This procedure creates an empty shell file to hold output data

*7.1.23 setup-gillnet-bycatch-data-file*

This procedure creates an empty shell file to hold output data

*7.1.24 setup-gillnet-fishing-days-data-file*

This procedure creates an empty shell file to hold output data

*7.1.25 setup-settings-file*

This procedure creates an empty shell file to hold model settings data

*7.1.26 setup-dolphins*

Figure B.11 is the flowchart for the setup-dolphins procedure.

Sliders on the UI determine the initial number of dolphins and which regions around New Zealand to simulate. Table B.3 shows the UI slider, the regions, and the corresponding dolphin placement areas included in that region (reference Figure B.5 for dolphin placement areas).

This procedure treats the sliders as separate populations; i.e. if *initial_maui* is 0, no Māui dolphins will be created. If the *initial_hectors_banks_peninsula* slider is > 0, dolphins will be created and placed in south-banksnorth and south-bankssouth areas, etc. The *generate-dolphins* procedure creates the dolphin groups, selects the number of individual dolphins in the group (group_size) and creates the dolphins. Māui dolphins are placed in specific areas around the North Island to match local densities from field data (Slooten et al. 2006a; Roberts et al. 2019) while Hector's are placed around the South Island per their own local distributions (MacKenzie and Clement 2014, 2016, 2019; Turek et al. 2013). For example, 12% of all Māui dolphins start in the "north-east" area, 19% start in "north-taranaki" and 69% are in the "north-west." Dolphins are also initialized at specific distances from shore within their placement area based on sighting data; e.g., for Hector's dolphins placed around Banks Peninsula, 43% are placed in the "near" distance band, 46% are in the "middle" distance

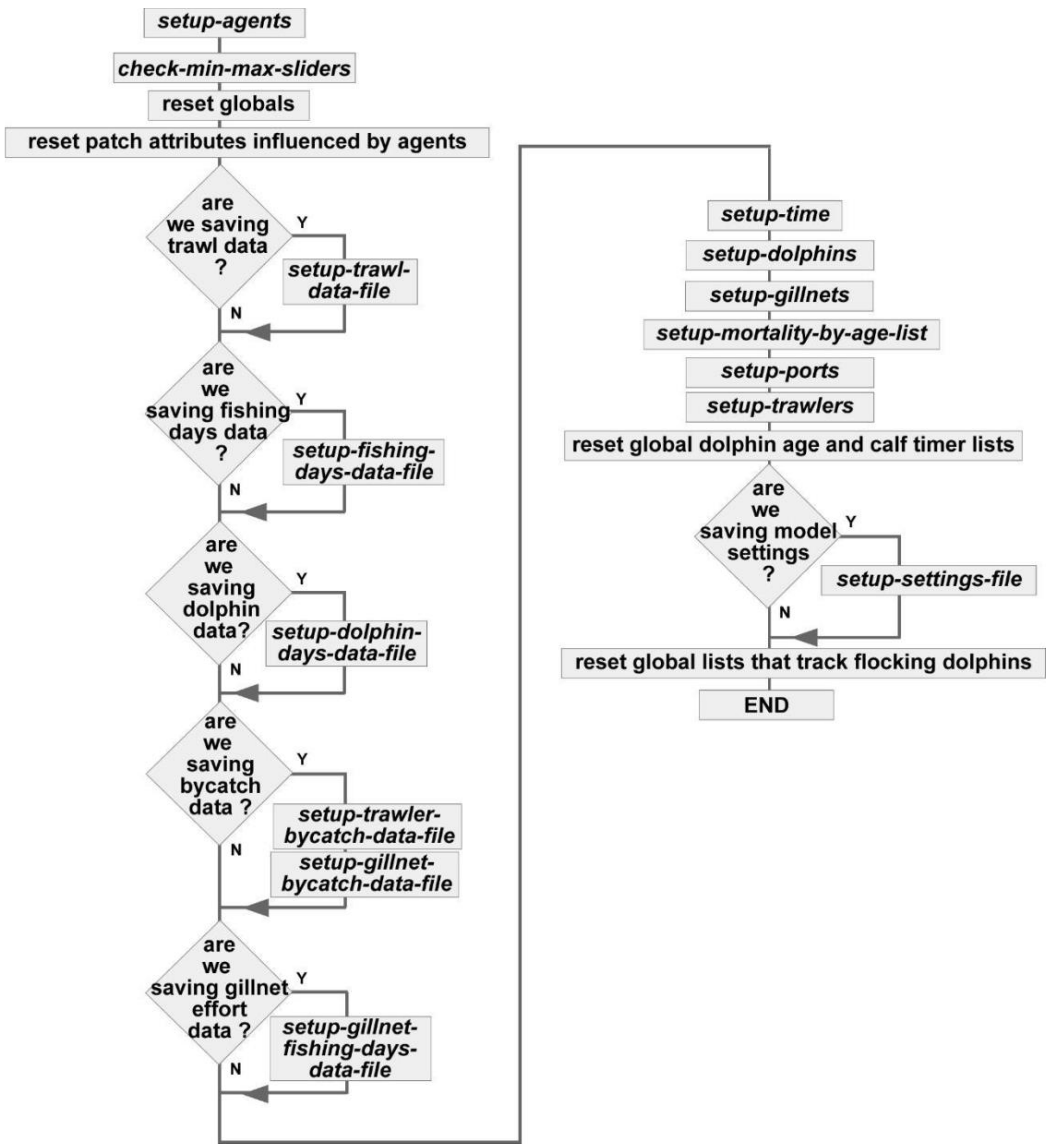

**Figure B.10** Flowchart for setup-agents procedure.

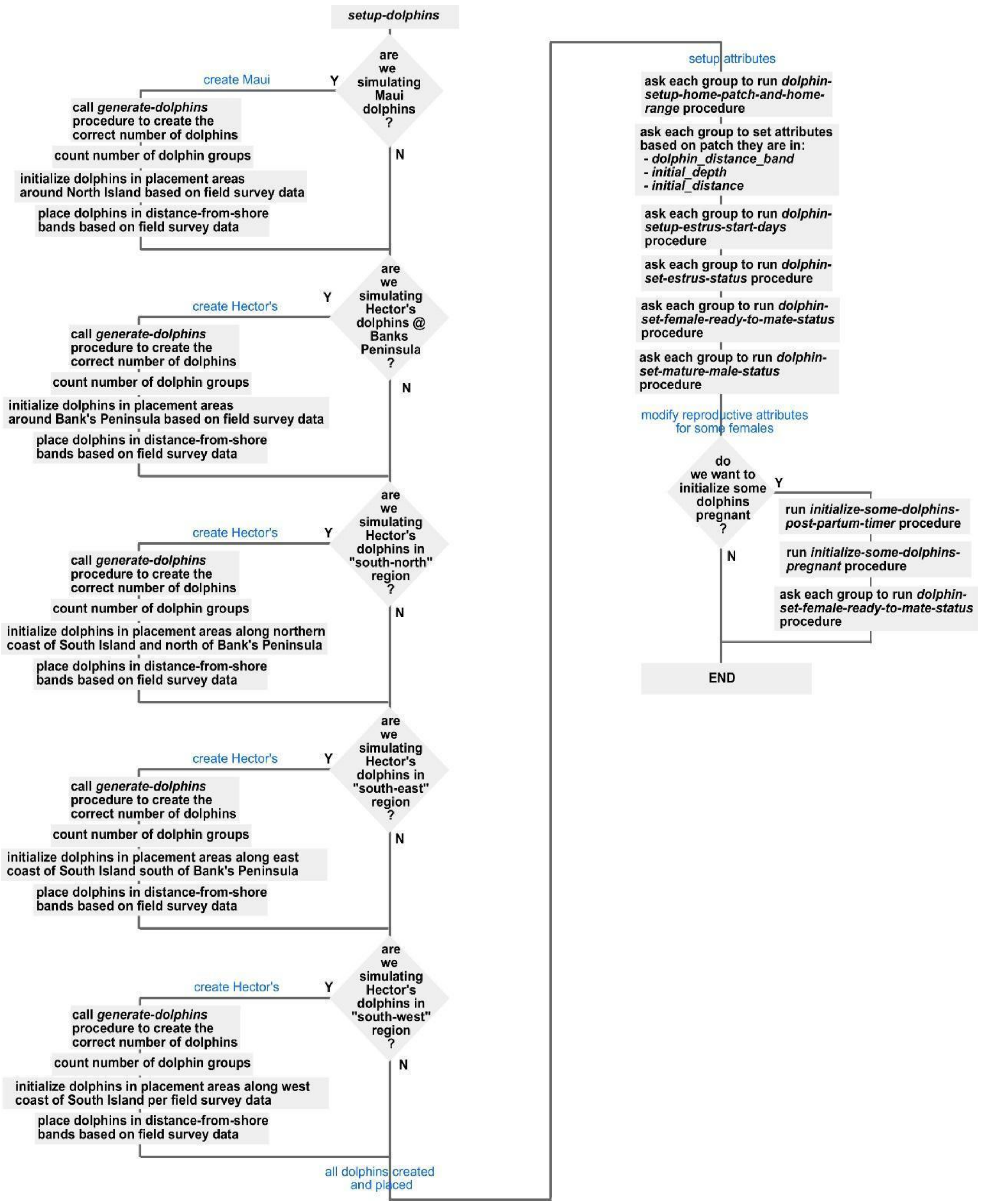

**Figure B.11** Flowchart for setup-dolphins procedure.

**Table B.3** Sliders for Initial Dolphins and Placement Areas:

| UI slider | NZ region | Dolphin Subspecies | Dolphin Placement Areas |
|---|---|---|---|
| initial_maui | all of North Island | Māui | north-west, north-east, north-taranaki |
| initial_hectors_banks_peninsula | South Island around Banks Peninsula | Hector's | south-banksnorth, south-bankssouth |
| initial_hectors_south_north | South Island, northern coast and east coast north of Banks Peninsula | Hector's | south_cloudybay, south_kaikoura, south_north |
| initial_hectors_south_east | South Island, east coast south of Banks Peninsula and most of southern coast | Hector's | south_otago, south_south |
| initial_hectors_south_west | South Island, west coast | Hector's | south_west, south_fiordland |

band, and the remaining 11% are in the "far" band (Slooten et al. 2006b; MacKenzie and Clement 2014, 2016, 2019). The procedure continues with adding more details for dolphin attributes. The *dolphin-setup-home-patch-and-home-range* procedure establishes each dolphin group's home range details. Female dolphins > six years old select their estrus start days. Those that happen to start estrus on day one of the year will start the simulation in estrus and are ready to mate; remaining mature females will have to wait before they enter estrus (*dolphin-set-estrus-status*). Dolphins run *dolphin-set-female-ready-to-mate-status* and *dolphin-set-mature-male-status* procedures to identify whether there are females ready to mate (not pregnant, in estrus, and postpartum timer has expired) and/or mature males (> six years old). Finally, if the user wants to initialize some dolphins as pregnant and other dolphins still in their postpartum recovery (via a switch on the UI), the model runs three procedures. In the *initialize-some-dolphins-post-partum-timer* procedure, the model assumes 50% of females > seven years old gave birth the previous year and sets their postpartum timer so they start the simulation with some postpartum time remaining before they can mate. The *initialize-some-dolphins-pregnant* procedure assumes 20% of eligible females ≥ seven years old successfully mated last year, are currently pregnant, and adjusts their gestation time to deliver calves randomly between days 1-59 to match field observations (Slooten 1991; Gormley 2009). It reruns the *dolphin-set-female-ready-to-mate-status* procedure to remove those females who were just declared pregnant or now have a non-zero postpartum timer.

*7.1.27 generate-dolphins*

This procedure creates the dolphin groups and the individual dolphins within them per Figure B.12. The procedure is passed the number of total individual dolphins and the subspecies. Each dolphin entity is a group, so the model first creates a dolphin group, then uses a random inverse gaussian distribution based on field data for each subspecies to pick the desired group size. As each group is created, the model tracks the total individual dolphins in those groups and keeps adding new groups until the total individual number of dolphins matches the desired number of dolphins. The gender of an individual dolphin is decided by a 50-50 random draw. Ages are selected from a random procedure that uses a custom distribution based on estimated age distributions of dolphin populations (see *select-dolphin-initial-age*).

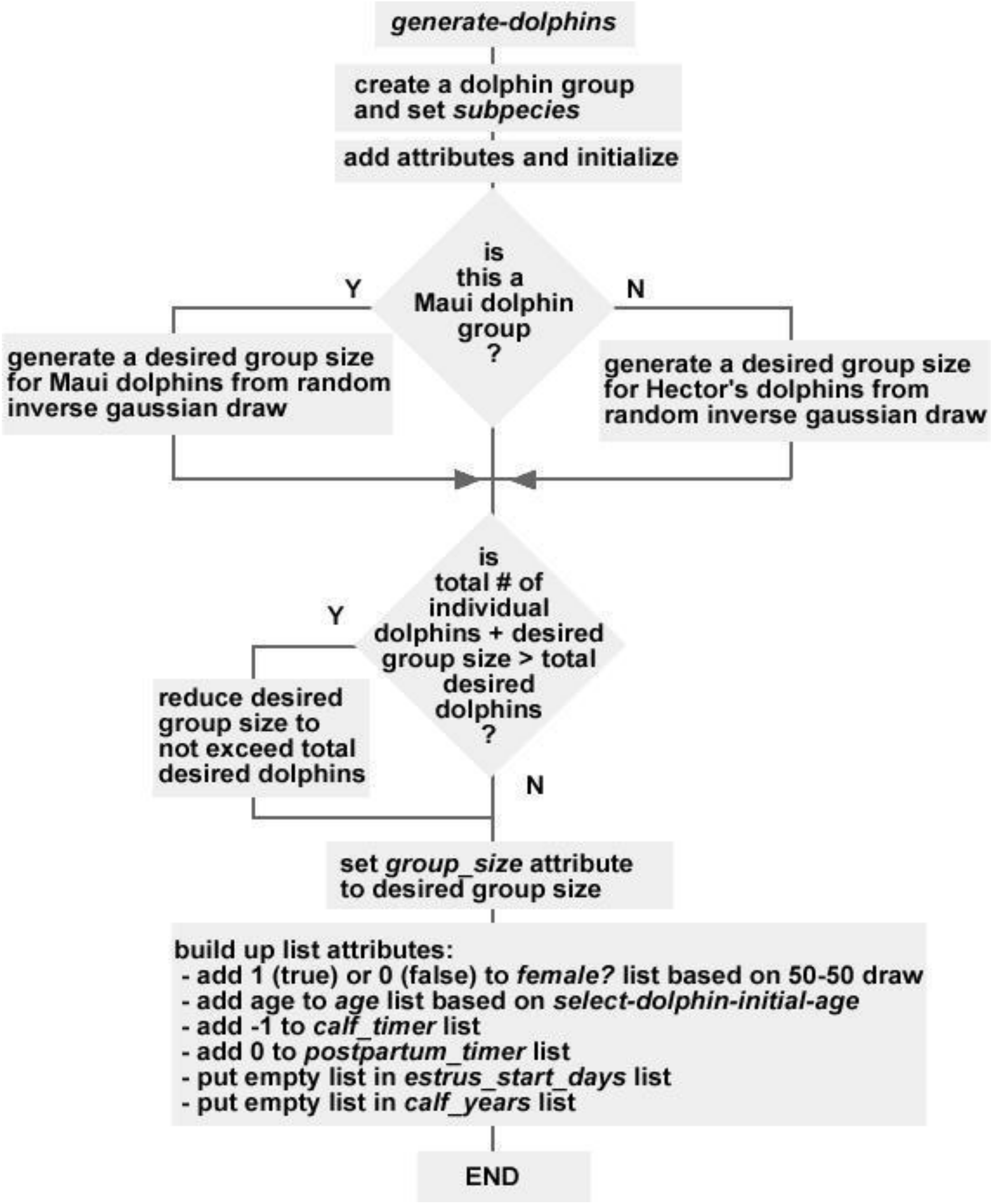

**Figure B.12** Flowchart for *generate-dolphins procedure.*

*7.1.28 dolphin-setup-home-patch-and-home-range*

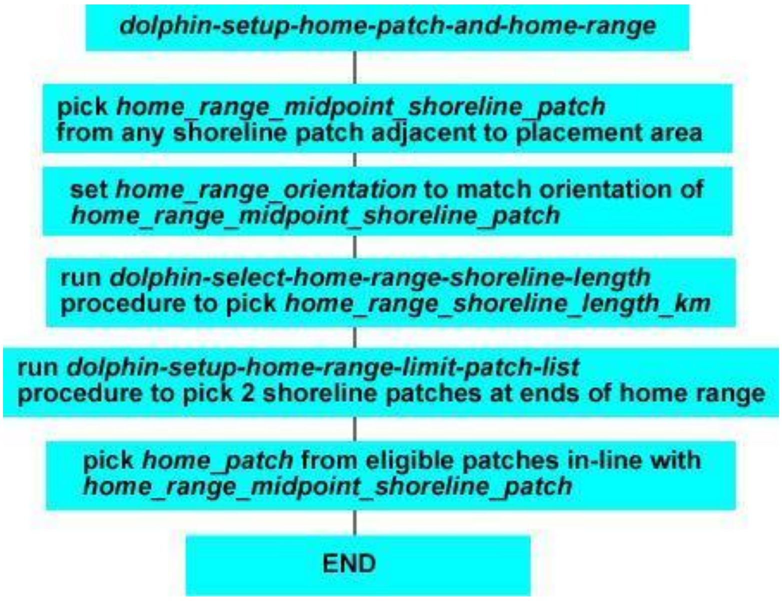

**Figure B.13** Flowchart for *dolphin-setup-home-patch-and-home-range* procedure.

Figure B.13 details the *dolphin-setup-home-patch-and-home-range* procedure. Dolphins randomly pick a home range midpoint shoreline patch from shoreline patches adjacent to their placement area. This midpoint patch defines the center of the home range. They select the home range shoreline length from a random-normal distribution based on a mean shoreline length of 50 km (Rayment et al. 2009) and standard deviation of 2.5 km. This allows them to pick the two patches on the shoreline that define the limits of shoreline travel and still be within the home range. They next pick a random home patch from ocean patches that are: between 0 and 5 km offshore, $\leq$ 60 m deep, and in-line with the home range midpoint shoreline patch (so home patch is centered relative to the length of the shoreline home range). The home range orientation is either “NS” (North-South) or “EW” (East-West) and is inherited from the shoreline midpoint patch’s orientation attribute (set in the *setup-shoreline-patches-home-range-orientation* procedure). Figures B.14 and B.15 show each orientation with the actual home range (light blue) bordered by shoreline patches (green) and the max depth limit (dark blue). The figures also show the home range midpoint shoreline patch, the home patch, and home range distance values used in dolphin movement procedures.

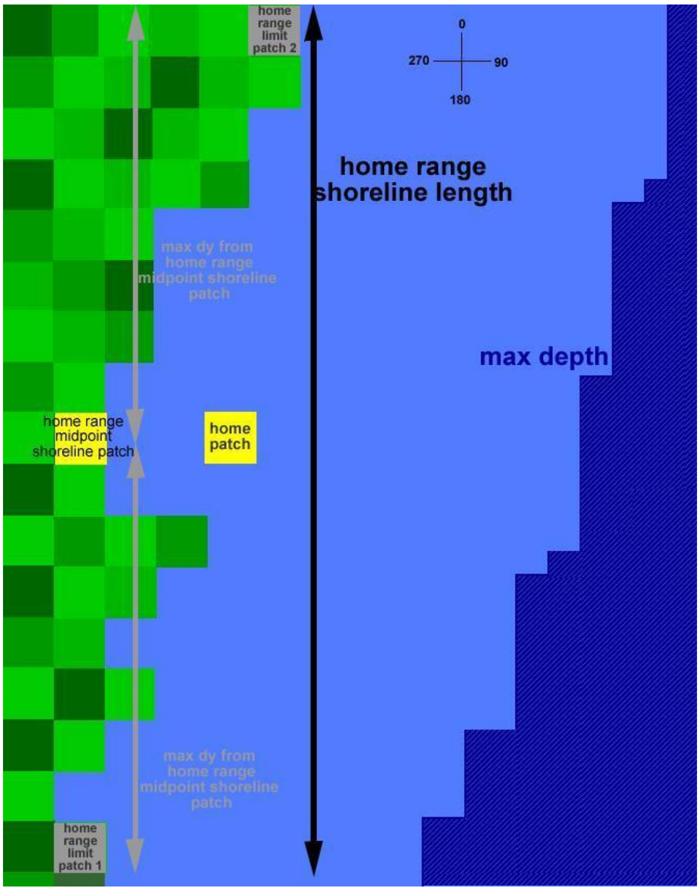

**Figure B.14** Example of “North-South” Dolphin Home Range (not to scale).

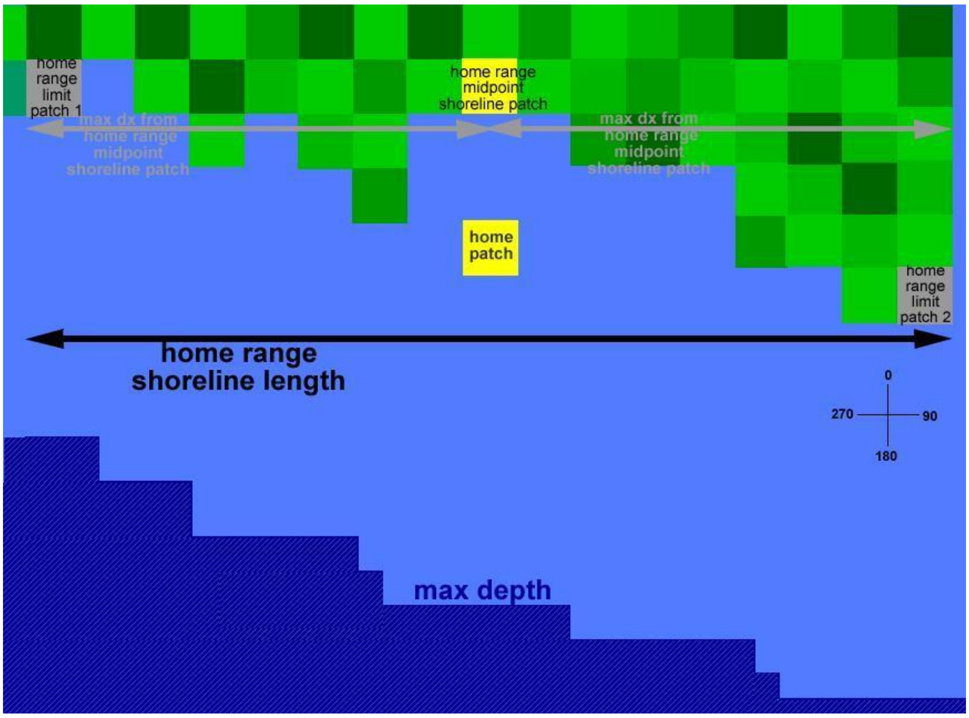

**Figure B.15** Example of "East-West” Dolphin Home Range (not to scale).

*7.1.29 select-dolphin-initial-age*

This procedure returns the initial age of a dolphin based on a random number draw from 0-1. The random number is passed to a table built from a stable age distribution derived from a Leslie Matrix model with the same survival and reproductive rates as the Netlogo model (Slooten et al. 2000). The table returns the age of the dolphin. An additional random number draw from 0-1 is added to age to represent fractional years. E.g., if the first random draw returns 0.32, the dolphin is 5 years old; if the second random draw returns 0.83, the final dolphin age is 5.83 years. Figure B.16 shows the resulting age distribution.

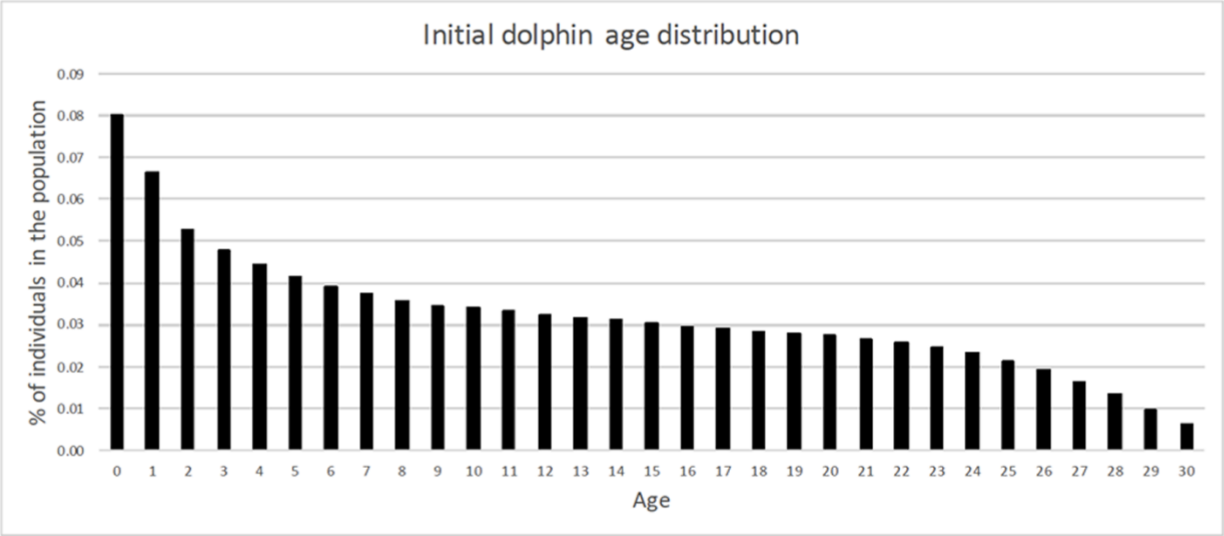

**Figure B.16** Initial Dolphin Age Distribution.

*7.1.30 dolphin-select-estrus-start-days*

Each mature female (minimum six years old) has at least one seven-day estrus period occurring within the first two months of the year so that the nominal gestation of 10-12 months will result in births in November-February (Slooten 1991; Gormley 2009; Bennington 2025). Six-year-olds have three estrus periods, seven-year-olds have two, and those eight and older have just one. These frequencies were chosen to replicate age-based ovulation rates (Slooten 1991; Gormley 2009). The actual days that estrus starts are stored in each dolphin's *estrus_start_days* list. When a five-year-old female turns six, it runs a procedure to pick three estrus start days from Julian day 1 to day 52 to endure the last estrus period finishes no later than day 59 (Feb 28). The first day is the only random day; it is chosen from a random uniform draw from days 1-22, with the second day 15 days after the first, and the third day 15 days after the second. For example, if the first estrus day is day 5, the second will be day 20, and the third will be day 35. When the six-year-old turns seven, she picks two new estrus start days with the first being from day 1-22 and the second 30 days later. When she turns eight years old, she picks a single estrus start day from day 1-52 and she retains that start day for the rest of the simulation.

*7.1.31 setup-gillnets*

This procedure is called at model startup, as well as the first day of each simulation year. Gillnets are only placed in areas where dolphins have been allocated (which is controlled by UI sliders), so the first thing this procedure does is determine which areas should have gillnets. It calls the *setup-goal-gillnet-fishing-days-by-area-table* procedure to build the table that stores the desired number of gillnet fishing days for each area. Gillnet effort is measured in gillnet fishing days; one gillnet fishing day is one gillnet deployed in a patch for 24 hours. It calls the *setup-actual-gillnet-fishing-days-table* procedure to build the data table that tracks the actual gillnet fishing days. The model loops through each eligible area and calls the *calculate-number-of-gillnets-in-area* procedure; this is how many gillnets should be in the area in the current year. It also counts the number of gillnets already in each area. If the actual gillnets are less than the goal, it creates enough gillnets via the *generate-gillnets* procedure to make up the difference. If the number of actual gillnets is more than desired, the model randomly removes existing gillnets until it matches the goal. Next, the model uses the *setup-gillnet-deployment-probability-table* to calculate and store the odds a gillnet is actually deployed each day. Finally, each gillnet runs the *gillnets-decide-if-deployed* procedure to actually do the probability check and set their *deployed?* attribute to true or false. Gillnets that are deployed count towards actual gillnet fishing days and can kill dolphins via bycatch.

*7.1.32 setup-goal-gillnet-fishing-days-by-area-table*

This procedure opens an external csv and extracts the historical (1970-2023) and forecast (> 2023) annual gillnet fishing days for each placement area for the current simulation year. It inserts the data into the *goal_gillnet_fishing_days_by_area_table* to be used as a reference in other procedures.

*7.1.33 setup-actual-gillnet-fishing-days-by-area-table*

This procedure builds an empty table that stores the actual gillnet fishing days for each area. Every day, the model counts up the deployed gillnets by area and adds them to the value in this table. At the end of the year, it can export the table data to confirm gillnet fishing effort matches the desired effort.

*7.1.34 calculate-number-of-gillnets-in-area*

This procedure refers to *goal_gillnet_fishing_days_by_area_table* to get the number of desired gillnet fishing days (e.g. 2000), divides by 365, then rounds up to the next integer (2000/365 = 5.47 = 6) to determine the number of gillnets needed in the area for the year. It does this for all areas that will have gillnets.

*7.1.35 generate-gillnets*

This procedure generates the actual gillnet entity, which then uses the *gillnet-select-next-patch* procedure to pick its patch, and moves to it.

*7.1.36 setup-gillnet-deployment-probability-table*

This procedure creates the *gillnet_deployment_probability_by_area_table* to store the probability each gillnet will be deployed that day in their placement area. Each area has its own calculation. For example, assume 500 gillnet fishing days is the target for the area. The model has already calculated and created the minimum integer number of gillnets needed to get 500 deployed days in a year (500/365 = 1.37 which requires 2 gillnets minimum). Since there are more gillnets than needed, the model calculates the deployment probability for this area as:

$$500\ gillnet \frac{days}{yr} / (2\ gillnets\ *\ 365 \frac{days}{yr}) =\ 0.68$$

It stores the result in the *gillnet_deployment_probability_by_area_table*. This process is repeated for all placement areas with gillnets.

*7.1.37 gillnet-decide-if-deployed*

This procedure reads the *gillnet_deployment_probability_by_area_table* to get the probability each gillnet will be deployed that day in their placement area. Each gillnet in the area performs a random uniform draw and if the number is ≤ the probability for its area, the gillnet is deployed.

*7.1.38 setup-ports*

The model refers to the UI settings for initial dolphins in regions around New Zealand (Table B.2), and builds a list of ports that are in placement areas that have dolphins. The model then generates those ports and places them at pre-set coordinates per the *port_locations_table*. The model also creates the waypoints (navigational aids used by trawlers to get out of the port) via the *setup-ports-waypoints-lists* procedure.

*7.1.39 setup-ports-waypoints-lists*

This procedure creates the *outbound_waypoints_list* and *inbound_waypoints_list* for each port. Each port reads an external csv to build a list of patches that will hold its outbound waypoints, in order (closest to port first). Then it calls *setup-outbound-waypoints-list* procedure to actually create the waypoints in those patches and put them in the *outbound_waypoints_list*. Finally, it reverses the outbound list to create the *inbound_waypoints_list.*

*7.1.40 setup-trawlers*

During initialization, the model refers to the simulation start year (e.g. 2024) and an external file provided by MPI (2023) that stores historical trawler counts by size, port, and calendar year. Trawlers are created and placed at each port to match data in that file. Trawlers select the day of their first trip. Trawler length, trawling velocity, travel velocity, and wake up time are assigned as follows.

- Small Trawlers:

$$length\ (m) = 10 + random\ 6$$

$$trawling\ velocity\ (knots) = 2.21 + random - normal\ 0\ 0.5$$

$$travel\ velocity\ (knots) = 8 + random - normal\ 0\ 1$$

$$wake\ up\ time = 4 + random\ 4$$

- Medium Trawlers:

$$length\ (m) = 16 + random\ 10\ m$$

$$trawling\ velocity\ (knots) = 4.01 + random - normal\ 0\ 0.5$$

$$travel\ velocity\ (knots) = 9 + random - normal\ 0\ 1$$

$$wake\ up\ time = 4 + random\ 4$$

At the start of every year the model refers to the same external file to compare actual trawlers vs. desired trawlers at each port. If the actual trawlers are less than the desired, it creates enough trawlers to make up the difference per the setup process described above. If the number of current trawlers is more than desired, the model randomly removes existing trawlers until it matches the goal. This procedure is called during setup where it refers to the initial year to create the historically correct number of trawlers by size and home port. It is also called at the start of each year to add/remove trawlers to match historical counts and ports.

#### 7.2 Housekeeping Submodels

The housekeeping processes manage model time information: hour, day of year, year of simulation, etc. They also update numbers of trawler and gillnet entities to match historical fishing effort, update patch protections as required, and eventually end the simulation. Figure B.17 is a more detailed flowchart of the housekeeping schedule and processes. All procedures have the opportunity to run every hour, but many of them run only on specific dates and times.

- *update-time:* This procedure controls how and when the model's time, day, and year are updated.
    - The time of day runs from hour 0 to 23; the *update-hour-of-day* procedure increments it by one.
    - When the time of day goes to 0, the *update-day-of-year* procedure increments the Julian day (day 1-365). When *update-day-of-year* sets the day = 274 (October 1), it runs *update-patch-protection*, which refers to the protection pulldown on the UI:
        - If *protection* = "year", it refers to calendar year and a table that identifies which patches are protected from gillnet and trawler fishing and sets protected patches to match; this model allows the protection scheme to change with the calendar year.
        - If *protection* is anything else, e.g., "2008", or "IUCN", the protected patches are set to match that specific protection scheme and will not change
    - When time of day is 0 and day of year is 1, the model runs *update-year-of-sim.*
- *update-year-of-sim:* If it is hour 0 of day 1 of the new year, this procedure increases the year of simulation and calendar year variables by one. It also kicks off multiple tasks that only happen at midnight on the first day of the year.
    - It calls the *setup-trawlers* procedure which refers to the new calendar year and may adjust the number and size of trawlers in the simulation to match historical effort.
    - The model calls the *setup-gillnets* procedure to adjust number and location of gillnets in each of the 12 placement areas. The model creates and places the gillnets depending on the current year and the goal gillnet effort (measured in

gillnet fishing days), which is based on historical data up to 2023 and then estimated future effort for subsequent years.

- If the year = 2009, the model runs *update-trawler-gradient* and *update-gillnet-gradient* procedures which update the trawler and gillnet gradient patch attributes. Trawler gradients indicate the historical fishing effort in each ocean patch and influence where trawlers choose to fish; gillnet gradients reflect historical gillnet effort and influence where gillnets are placed. The probability that a trawler and gillnet choose a specific patch increases as the gradient increases. There are two data sets for each gradient; data prior to 2009 and then a more recent set used from 2009 onward.
- Trawlers record their total fishing days for the previous year in *trawler-write-total-fishing-days* procedure. Then they immediately clear out the data in *trawler-reset-fishing-days-this-year*.
- Finally, the observer compares the year of the simulation (e.g. year 20) to the desired number of years the simulation should run and ends the simulation when required.

- resets global lists that track calf timers: every day at midnight, the observer resets the global list that stores the calf timers of Māui dolphins and another one for Hector’s dolphins. The lists are used to feed histograms on the UI.

- resets global lists that track dolphin ages: every day at 23:00, the observer resets the global list that stores the ages of Māui dolphins and another one for Hector’s dolphins. The lists are used to feed histograms on the UI.

- calls update-actual-gillnet-fishing-days procedure which counts up all the gillnets that are deployed in each placement area to compare with the goal gillnet fishing days

- calls write-dolphin-population-data procedure: on the last day of the month at 23:00, the model records dolphin population data to a csv for post-processing and data visualization.
- tick; model advances to the next hour in the simulation

These processes also:

- calls *count-mature-females* procedure: model counts the total mature (≥ six years old) female Māui and Hector's dolphins; data is used on the UI
- calls *count-pregnant-females* procedure: model counts the total pregnant Māui and Hector's dolphins; data is used on the UI

#### 7.3 Trawler Submodels

##### *7.3.1 Overview of Trawler Processes*

Trawler processes, shown in Figure B.18, are grouped within dashed boxes to represent the major trawler activities: leave port, trawl for fish, and return to port. The actual trawler procedures are indicated by italics; e.g., *trawler-select-fishing-patch*, *trawler-decide-what's-next*, etc.

Trawlers are classified as "small" (length ≤ 15m) or "medium" (16-25m). Medium trawlers cannot fish in patches protected from trawling; they can travel through them but not actually fish there. As long as the protection scheme is not "IUCN" or "IUCN Plus", small trawlers are exempt and can fish anywhere they choose with depths ≤ max trawl depth limit. Under the "IUCN" and "IUCN Plus" protections, all trawlers avoid fishing in protected patches. Large trawlers (> 25m) rarely fish in waters used by Hector's and Maui dolphins, and are not considered in our model.

Trawlers are created at model startup and assigned to a home port based on the calendar year and an external csv that sets the number of small and medium trawlers at ports around New Zealand. The csv has historical trawler counts and home port data from 2008-2022; from 2022 onward, the model uses the 2022 values.

Trawlers leave from and return to their home ports after each fishing trip. Small trawlers leave and return to port on the same day (mean trip duration is 11 hours) while medium trawlers are out of port for multiple days (mean trip duration is 96 hours).

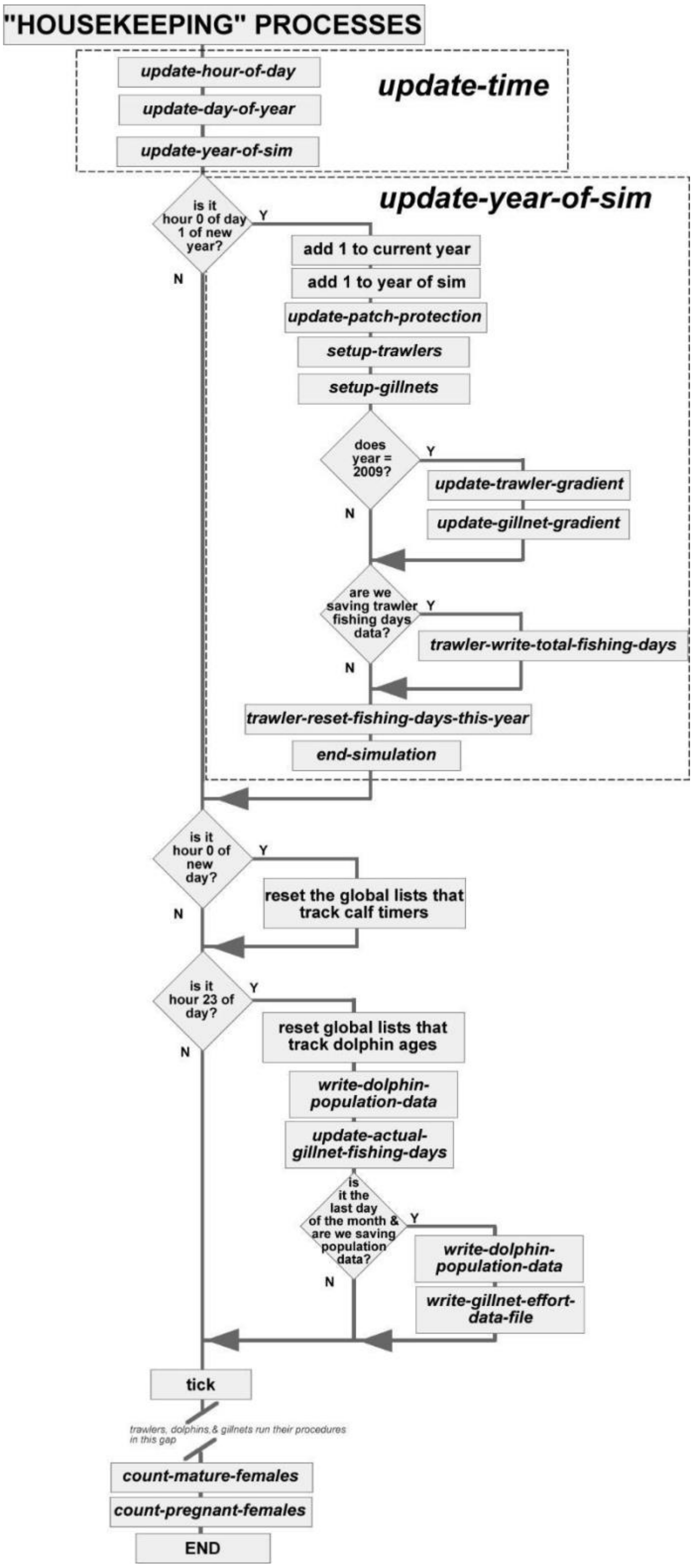

**Figure B.17** Housekeeping Procedures.

Trawlers go through the following steps:

a) *trawler-wake-up*: This procedure runs every hour, and it controls whether a trawler stays in its home port or leaves on a new trip. It compares the simulation's day of year and time of day to the trawler's scheduled next trip day and start time; when those match, it's time for the trawler to "wake up" and start a trip. The trawler sets a flag variable *fishing_complete?* to false to indicate fishing is **not** complete, which spurs the trawler to start fishing. Otherwise, it leaves *fishing_complete?* true, which keeps the trawler in port for at least one more hour.

b) *trawler-decide-what's-next*: The trawler decides what activity it will undertake before each time step, and sometimes within the same time step if it completes an activity with time remaining in the hour; e.g., traveling to a fishing patch and having time left to start a trawl. The decision process splits based on the *fishing_complete?* variable;
   - *trawler-decide-what's-next-fishing-complete*: Trawler is done fishing. If it is in port, it remains. If it is away from port, it will travel back to port and continue traveling to port until it arrives and ends the trip.
   - *trawler-decide-what's-next-fishing-incomplete*: The trawler can begin a fishing trip (if in port) or keep fishing if it is already out on a trip. When it is time to depart on a new trip, the trawler selects the length of the trip in hours (*trawler-select-trip-duration*) and the fishing patch to start in (*trawler-select-fishing-patch*). Every hour the trawler is out on a trip, the first thing it does is estimate the travel time back to port based on distance and travel velocity, and compare that with the time remaining to make it back to port on schedule. If the travel time to port is $\geq$ remaining trip duration, the trawler decides to return to port and will continue traveling to port until it arrives and ends the trip. Otherwise, with time left to fish, the trawler evaluates where it is and works through the steps to start or continue a trawl. If the trawler is not in its desired fishing patch, it will travel to the patch. If the trawler has arrived, it will set its activity to fishing.

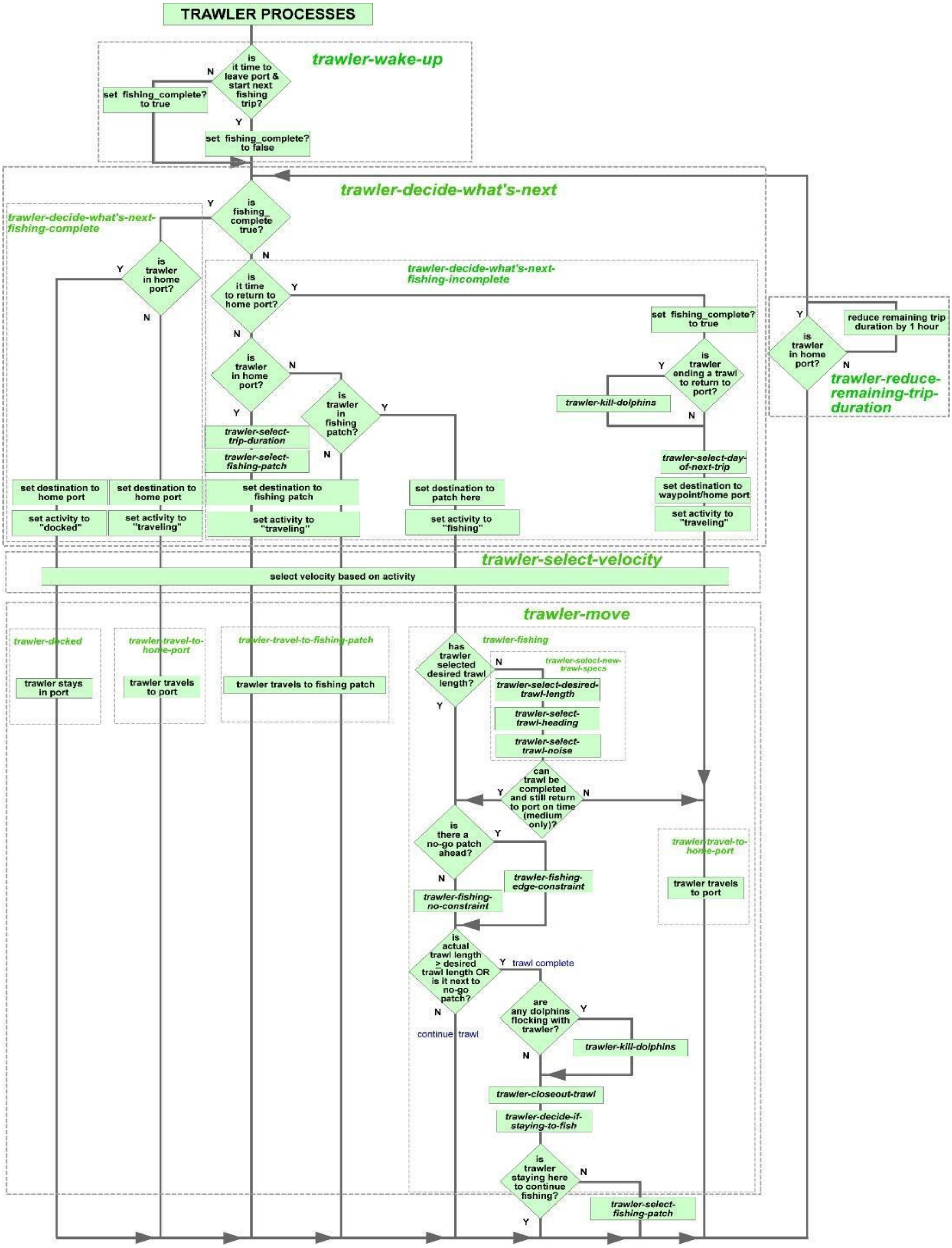

**Figure B.18** Trawler Processes Flowchart.

c) *trawler-select-fishing-patch*: Trawler selects a specific patch to start fishing in. This could occur while trawler is still in port picking the initial patch or out in the open ocean. See *trawler-select-fishing-patch* procedure for detailed description.

d) *trawler-select-velocity*: The trawler chooses a velocity depending on activity. If the trawler is docked in port, velocity is 0. If fishing, it chooses its trawling velocity. All other activities involve traveling somewhere, so it chooses travel velocity.

e) *trawler-select-day-of-next-trip:* At simulation startup and every time a trawler ends a fishing trip, it chooses the day it will leave for its next trip based on fishing effort data for small and medium trawlers. See submodel for specifics.

f) *trawler-move*: The trawler executes the chosen activity. It will use one of the following subprocedures to actually move:

  - *trawler-docked*: This procedure keeps the trawler in port until its next fishing trip.
  - *trawler-travel-to-home-port:* Trawler travels back to home port at travel velocity; once it makes this decision it will continue to travel to port until it arrives.
  - *trawler-travel-to-fishing-patch*: Trawler travels to chosen fishing patch. See *trawler-travel-to-fishing-patch* for detailed description.
  - *trawler-fishing*: This procedure controls the trawlers when they are actually fishing, whether starting a new trawl or continuing one from the previous hour. The actual trawling process occurs in "mini-steps", which is the trawling velocity in patches/hour broken up into 10 segments. Because of patch geometry and the way Netlogo "vision" works, we found mini-steps prevented the trawler from ending up or traveling through a no-go patch. Each trawler:
    - Runs *trawler-select-new-trawl-specs.* When the trawler arrives at a new fishing patch, or decides to stay in a patch and start a new trawl, it:

- Selects a desired trawl length (km) via *trawler-select-desired-trawl-length* procedure
- Runs *trawler-select-trawl-heading* where trawler evaluates the surrounding area and picks a heading for the trawl. See *trawler-select-trawl-heading* for detailed description
- Runs *trawler-select-trawl-noise* procedure to select the actual trawl's emitted noise level in dB; this is what attracts the dolphins to the trawler

- (Medium trawler only) calculates the estimated time the trawl will finish based on trawl length and trawling velocity versus the time it has to stop fishing and return to port. If completing the trawl pushes it past return time, it will select a shorter trawl length to stay on schedule. If the shortened trawl length is < 5km, it will not start the trawl and head to port a little early.
- Performs a "look-ahead" to check for "no go" patches on the current heading a short distance ahead. Small trawlers are looking out for land and protected patches **only** when the protection scheme is IUCN or IUCN Plus, while medium trawlers look for land or protected patches under all protection schemes. If there are "no go" patches ahead the trawler runs *trawler-fishing-edge-constraint* to trawl as far as it can without entering the "no go" patch. If the path forward is clear, it uses the *trawler-fishing-no-constraint* procedure. These procedures also alert dolphins within the detection range when trawling is starting and ending.
- When the trawl ends, from either reaching the desired trawl length or encountering a no-go patch, the trawler:
  - Runs *trawler-kill-dolphins*: Trawlers have an opportunity to kill dolphins that are flocking with them every time a trawl ends

which represents when the net comes out of the water. See details in *trawler-kill-dolphins* procedure.

- Runs *trawler-closeout-trawl:* This officially ends the trawl with the trawler saving detailed trawl data (trawl length, start and end coordinates, etc.) that can be exported to an external file. It also updates global variables that track total trawls, length of trawls, etc.
- Runs *trawler-decide-if-staying-to-fish:* Decides whether it will stay in the same patch for its next trawl or relocate.

g) *trawler-reduce-remaining-trip-duration:* Trawlers that are not in port run this procedure to reduce the remaining trip time by one hour

*7.3.2 Detailed Trawler Submodels*

*7.3.2.1 trawler-select-day-of-next-fishing-trip*

The day of a trawler's next fishing trip depends on how many days they decide to stay in port. Trawlers choose the maximum of either a fixed minimum number of days between trips or a randomly generated value based on a gamma distribution. All parameters are size-based; small trawlers have fewer days between trips compared to medium trawlers (GFW 2025; MPI 2023). The formula for all trawlers is:

*Days between trips = max(minimum days between trips, random-gamma alpha, lambda)*

Parameters are:

| Trawler Size | Min Days Between Trips | Days Between Trips *alpha* | Days Between Trips *lambda* |
|---|---|---|---|
| Small Trawler | 1 | 1 | 0.5 |
| Medium Trawler | 4 | 6 | 0.5 |

The actual day of the next trip is calculated as:

$$Day\ of\ next\ trip\ =\ simulation\ day + days\ between\ trips$$

*7.3.2.2 trawler-select-trip-duration*

Trawlers select the length of each trip (hours) with the same formula; the parameters differ since small trawlers leave and return to port on the same day while medium trawlers take multi-day trips.

| Trawler Size | Mean Trip Duration (hrs) | Trip Duration Std Dev (hrs) |
|---|---|---|
| Small Trawler | 11 | 0.25 |
| Medium Trawler | 96 | 10 |

$$Trip\ Duration\ (hrs) = round(random\ normal \quad mean\ trip\ duration \quad std\ dev\ trip\ duration)$$

*7.3.2.3 trawler-select-fishing-patch*

The trawler method to select a fishing patch is slightly different depending on location.

**From home port**: A trawler in its home port, first selects a group of ocean patches within the trawler's maximum initial travel time limit from home port; 3.5 hours for small or 6 hours for medium trawlers. They reduce that group to include only ocean patches that: satisfy minimum and maximum trawl depth limits; are unprotected (medium trawlers only); are within the home port's min-max bearing limits (each port has a minimum and maximum bearing that the trawlers use to prevent them picking a fishing patch that would cause the trawler to travel across land). For example, the eligible initial fishing patches for a medium trawler in the blue-starred port considering travel time and **without** considering intervening land are shown in black in Figure B.19.

Notice patches south and south-east of the port are initially eligible, but the trawler would cut across Banks Peninsula to get there. In Figure B.20, the patches in pink are eligible patches when we consider where the port is in relation to surrounding land.

From this subset of patches, it picks one via a weighted random draw based on the historical use of eligible patches; each ocean patch has an attribute derived from patch-specific historical fishing effort based on New Zealand government data (MPI 2023, 2024). Higher historical fishing data in a patch leads to greater odds of being selected. Finally, trawlers check the candidate patch to confirm it is "reasonable" to start the trawler's fishing trip. The "reasonable" check is a brief evaluation of the patches between where the trawler is and where it wants to go to prevent trawlers indiscriminately crossing land; the trawler makes sure the percentage of land patches in its path is $\leq$ 25%. *Note - This model does not do detailed navigation around land patches to avoid crossing* ***any*** *land; we traded the inaccuracy of travel paths for increased CPU performance from not having to plot accurate patch-by-patch directions to a fishing patch.*

If the first patch selected is not "reasonable", the trawler picks another from the subset of eligible patches and does the "reasonable" check again. It will repeat this process until an acceptable patch is chosen.

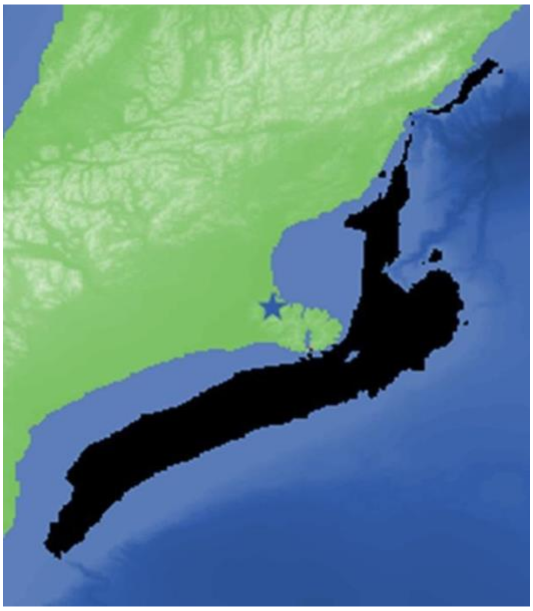

**Figure B.19** Potential fishing patches from home port before downselect.

**In open ocean:** If the trawler is out on the open ocean and has completed a trawl, it does a random uniform draw based on the probability that the trawler continues fishing at the current patch or a different patch. Both small and medium trawlers have a 50% chance of staying. If the trawler is staying, it waits until the start of the next time step before starting a new trawl (this delay replicates the time it takes for the trawler to prepare for its next trawl).

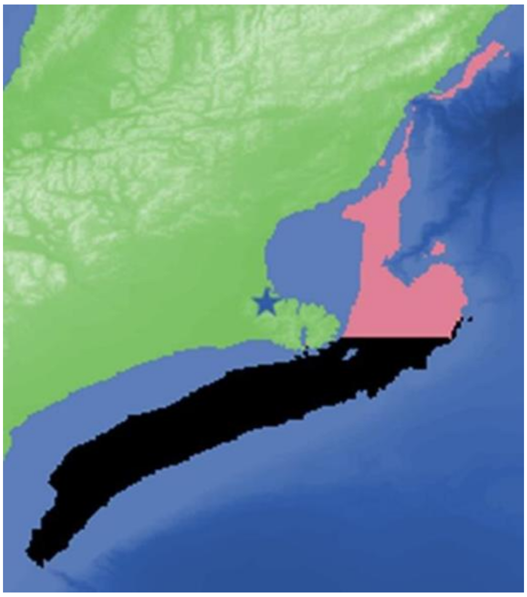

**Figure B.20** Potential fishing patches from home port after downselect.

If it wants to relocate, it considers all the ocean patches that are: within the maximum allowed travel time between trawls (small trawlers are allowed 2 hours; medium trawlers can travel 4 hours), within minimum and maximum trawl depth limits, and unprotected (medium trawlers only). It chooses the next fishing patch via the same historically-weighted random algorithm and “reasonable” test as it does from the home port. There is not an explicit delay for trawlers that relocate since it is assumed any prep work would be done in transit.

*7.3.2.4 trawler-travel-to-fishing-patch*

Trawlers run this procedure to actually move to their fishing patch no matter where they start. If they are in port, they use the port’s waypoints to navigate out into open water. In the open ocean they face the fishing patch and calculate the distance to the patch. Because the *distance* primitive in Netlogo measures to the center of the patch, the trawler deducts a random 200-500m from that distance to ensure it will make it within the patch, but not travel all the way to the center before it officially “arrives” and can start a trawl. If the distance to the patch is $\geq$ trawler’s traveling velocity (patches/hour), it will move the entire velocity in one tick. If not, the trawler moves only as far as it needs to, which means it has some velocity “left over” that it can still use within this hour to trawl. For example, assume trawler velocity is 8 knots which converts to 9.26 patches/hour (8 knots *1.852 km/hr/knot * 1 patch/1.6 km = 9.26 patches/hour) and the distance to get within the fishing patch is 3.81 patches. The trawler moves 3.81 patches, but still has 5.45 patches it **could** use in this hour at traveling velocity. It does not need to keep traveling, so the trawler will compute how much of the hour it has remaining to use for trawling as:

$$tick\ remaining = \ remaining\ velocity/full\ velocity$$

In our example, the trawler still has 5.45/9.26 = 0.59 of the hour remaining (~35 minutes) that it could use to trawl instead of travel. This rather complicated velocity tracking process ensures that the trawler fully utilizes each hour rather than take the whole hour to move the remaining 3.81 patches; that implies the trawler slows to a crawl for the last hour of travel which is not realistic. Since this example trawler is transitioning from traveling to trawling, the trawler will take the remaining fraction of the hour and convert it into a new remaining velocity in patches/tick by multiplying it by the trawler's nominal trawl velocity; i.e., the trawler will slow down and trawl for those last ~35 minutes. The trawler does this math every time it transitions between activities mid-time step. Other examples include switching from trawling to traveling to port, or ending a trawl and traveling to another fishing patch.

*7.3.2.5 trawler-select-desired-trawl-length*

Trawl lengths are selected to match field data (MPI 2023, 2024; GFW 2025). Using the formula:

$$Trawl\ length\ (km)\ =\ 5\ +\ random - gamma\ 1.5\ 0.125$$

The same gamma distribution of desired trawl length, regardless of trawler size, resulted in an excellent match to field data (Figures B.21 and B.22). Medium trawlers accomplished longer trawls because they work in open waters, further from the coast. Small trawlers completed shorter trawls because they operate closer to shore and therefore had to cut some of their trawls short when they approached the shoreline. Small trawlers spend fewer hours per day trawling, which also results in shorter trawls.

Medium trawlers only do an additional check before starting a new trawl; they check if there is enough time to complete the trawl (based on desired trawl length selected above) and still make it back to port on time. If the timing works, the trawler commits to the trawl. If not, it computes the maximum length trawl it could complete under the time constraints. If that new trawl length is ≥ 5 km, it commits to the shortened trawl; otherwise, it does not even start the trawl and heads back to port.

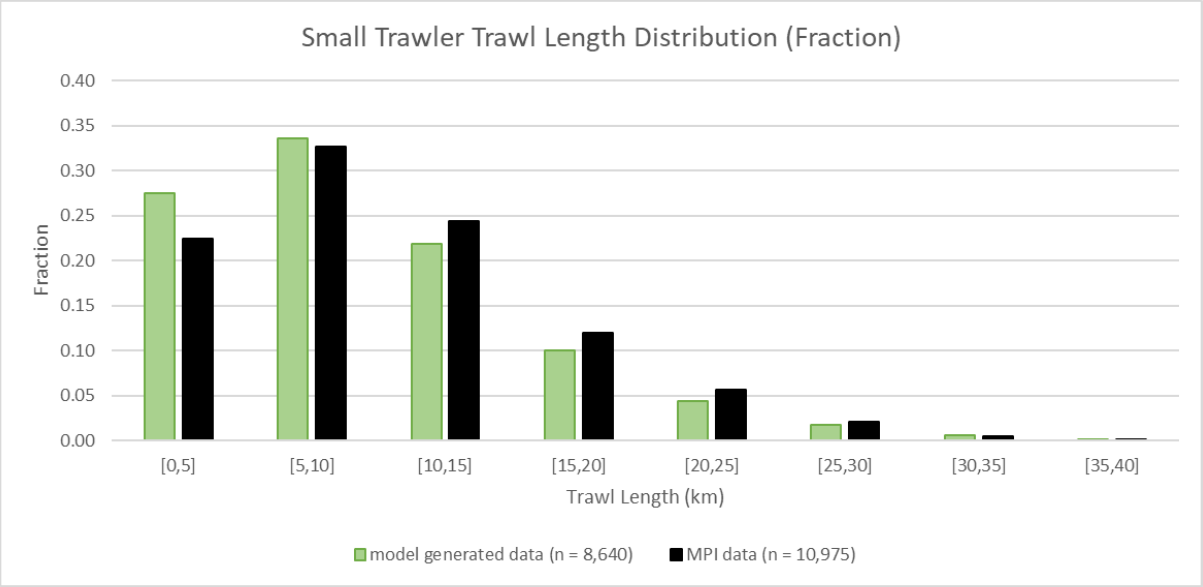

**Figure B.21** Proportion of trawls of different lengths for small trawlers.

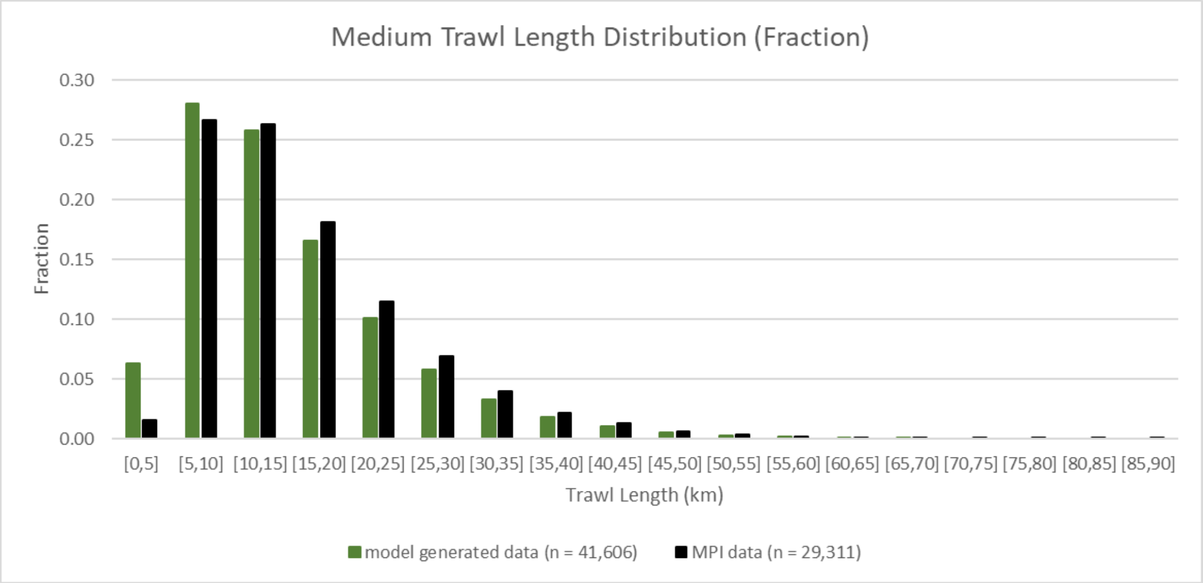

**Figure B.22** Proportion of trawls of different lengths for medium-sized trawlers.

*7.3.2.6 trawler-select-trawl-heading*

Government and AIS data show small and medium trawlers tend to follow the shoreline or a depth contour (MPI 2023, 2024; GFW 2025). The first thing a trawler does is identify the nearest patch with "runway numbers" which are attributes of patches along the shoreline, the 50m depth contour, and the 100m depth contour that indicate to the trawler the general heading of the contour. Next it determines which runway number is closest to its current heading and sets its initial heading to match plus a randomization factor based on the distance to the patch; i.e., the closer the trawler is to the patch, the closer its heading will be to the runway number. Before it starts to fish, the trawler does a brief look ahead to make sure the initial heading does not have it pointed into land or a protected patch (for medium trawlers or all trawlers under the IUCN or IUCN Plus protection scenarios). If so, it turns away from the no-go patch in small increments, checking after each micro-turn and stopping when the path forward is clear. The trawler only runs this procedure when it is starting a new trawl.

*7.3.2.7 trawler-select-trawl-noise*

The trawler select's the noise that the trawl will emit; this is what attracts dolphins to the trawler. Trawl noise depends on the size of the trawler and selected via a random normal draw based on the mean trawl noises set by UI sliders (Daly and White, 2021):

Small trawler trawl noise -> random normal with mean = sm_trawler_mean_trawl_noise dB and sd = 0.65 dB

Medium trawler trawl noise -> random normal with mean = med_trawler_mean_trawl_noise dB and sd = 0.65 dB

*7.3.2.8 trawler-fishing-no-constraint*

Trawlers alert nearby dolphins that trawling is underway via trawler-alert-dolphins-trawling-happening. This alerts dolphins to run procedures to determine if they detect the trawl noise and want to flock with this trawler. The trawler compares the remaining trawl length to the length of a "mini-step", which is 0.1 * trawling velocity. If the remaining trawl length is > than the mini-step, it moves the entire mini-step; it keeps moving in mini-steps until the remaining trawl length is < goal trawl length. Once that occurs, it moves forward only as far as it needs to achieve the desired trawl length, ends the trawl, and records the final trawl data (trawler-closeout-trawl). With the trawl complete, the trawler does a 50-50 draw to decide if it wants to start another trawl in the current patch on a fresh heading or relocate to another fishing patch. It also notifies nearby dolphins that trawling is complete via trawler-alert-dolphins-trawling-complete.

*7.3.2.9 trawler-fishing-edge-constraint*

A trawler is using this procedure when its next mini-step will take it into a no-go patch (land or a protected patch). The trawler moves only as far as it safely can, ends the trawl, and records final trawl data (trawler-closeout-trawl). We force the trawler to end the trawl because trawlers cannot turn with nets in the water. It does a 50-50 draw to decide if it wants to start another trawl here on a fresh heading or relocate to another fishing patch. It also notifies nearby dolphins that trawling is complete via trawler-alert-dolphins-trawling-complete.

*7.3.2.10 trawler-alert-dolphins-trawling-happening*

This procedure alerts nearby dolphins that the trawler is starting a new trawl or continuing a trawl from the previous hour. The trawler first counts how many dolphins are flocking with it; if there are already the maximum number of dolphins allowed, it will not send out an alert. Otherwise, it calculates the alert radius via the trawler-calculate-dolphin-alert-radius procedure and sends an alert to those dolphins within the radius and allowed to flock because their interval between flocking events has been satisfied. Rather than have all dolphins constantly listening for trawling nearby, we chose to have the trawling boats send out a signal to only those eligible to detect and act on it. The dolphins have a separate process that determines if they actually detect the trawl noise and whether or not they will move to the trawler (dolphin-detect-trawling-nearby-and-decide-to-travel).

*7.3.2.11 trawler-calculate-dolphin-alert-radius*

The trawler calculates the maximum distance where a dolphin has at least a 50% chance of detecting the trawl noise. The trawler starts by calculating the maximum transmission loses

(TL) allowed until received level of trawl noise matches the estimated minimum detection threshold:

$$TL\ (dB) = trawl\ noise\ (dB) - dolphin\ minimum\ detection\ threshold\ (dB)$$

Audiograms for Hector's and Maui dolphins are unavailable; therefore, the minimum detection threshold for broadband noise was estimated at 100 dB based on harbor porpoise audiograms (Kastelein et al., 2017). The trawler uses the TL equation to calculate the distance where that TL will occur:

$$TL = xlogR$$

$$R = 10^{TL/x}$$

$$where\ x = TL\ coefficient$$

Traditionally, 20 is used for x in deep water (spherical spreading) and 10 is used in shallow water (cylindrical spreading). In our case, trawling is carried out in intermediate depths so we used 15 for x, also known as "intermediate" spreading (Lynch and Newhall 2017).

*7.3.2.12 trawler-alert-dolphins-trawling-complete*

Every trawler that finishes a trawl sends out a signal to alert all the dolphins within the dolphins' trawler detection radius that trawling has ended. Rather than have all dolphins constantly listening for the conclusion of each trawl, we chose to have the trawling boats send out a signal only to those eligible to detect it.

*7.3.2.13 trawler-kill-dolphins*

At the conclusion of a trawl, whether it reached the desired trawl length, encountered a no-go patch, or had to end a trawl to return to port, the trawler has the opportunity to kill dolphins. First, the trawler totals up the number of dolphins in the flocking groups. It then applies a random uniform probability that a trawler-dolphin interaction results in mortality to each individual dolphin and totals up the number that will be killed. For example, if there are 50 flocking dolphins, and the probability of a trawler bycatch death is 0.01, the trawler does 50 random-uniform draws from 0-1 and those that come back < 0.01 count as a death and the total dolphins killed increases by one. Then it randomly removes enough individual dolphins from the flocking groups until it reaches the total kills calculated. It also records all the relevant mortality data for post processing analysis.

*7.3.2.14 trawler-decide-if-staying-to-fish*

This procedure determines if the trawler will start its next trawl in the same patch the last trawl ended or relocate to a new patch. Trawlers have different odds of relocating based on size so the formulas are:

$$small\ trawler\ probability\ of\ staying = 1 - small\_trawler\_switch\_trawl\_patch\_odds$$

$$medium\ trawler\ probability\ of\ staying = 1 - med\_trawler\_switch\_trawl\_patch\_odds$$

If a random number from 0-1 is < probability of staying, the trawler will stay in the same patch. Otherwise, it will choose a new patch for its next trawl.

*7.3.2.15 trawler-select-day-of-next-trip*

Trawlers run this procedure to select the day they will leave port on their next fishing trip. The model variable day_of_sim is a continuous count of days from 1 to the last day of the sim; e.g., if the simulation lasts 10 years, day_of_sim would run from 1-3650. Trawlers have different parameters that control the interval between trips based on size: minimum number of days between trips, and alpha and lambda parameters for a gamma distribution. The trawler first does a random gamma number draw with the alpha and lambda parameters to set an initial interval between trips. If that interval is > the minimum interval, that is the interval that will be used; otherwise, the trawler selects the minimum interval. The actual day of the next trip is based on the current day_of_sim:

$$day\ of\ next\ trip = day\_of\_sim + interval\ between\ trips$$

*7.4 Dolphin Submodels*

*7.4.1 Overview of Dolphin Processes*

Figure B.23 shows the detailed dolphin processes flowchart grouped by dashed boxes and top-level procedures. Note that not all procedures run every hour (e.g., dolphin-age only runs once per day).

They are:

*dolphin-decide-what's-next:* Dolphins decide what activity they will do next in this procedure. Dolphins are attracted to trawlers that are fishing, but they must clear two constraints before they can flock with a trawler. The first constraint is the dolphin's time between flocking events has expired (when a dolphin leaves a trawler because it has reached the maximum continuous amount of time it can stay with a trawler, there is an interval timer *(allowed_to_flock_countdown)* that prevents the dolphin from rejoining this or any other trawler until that interval has expired). The second is the number of dolphins flocking with this particular trawler is less than the max allowed *(max_dolphins_allowed_with_trawler)*. If both constraints are satisfied, the dolphin evaluates the distance to that trawler. If the dolphin is >10m away from the trawler it decides to travel to that trawler. If within 10m, it will stay close and flock with the trawler. If there is no trawler fishing within the *max_dolphin_trawling_detection_distance,* or the dolphin reaches its maximum allowed consecutive flocking time limit or the trawler stops fishing, the dolphin will decide to "wander" around within their home range.

a) *dolphin-select-velocity:* Dolphins select their velocity depending on the activity decision. Dolphins flocking with a trawler set their velocity to match the trawler's. Dolphins traveling to trawlers swim at maximum velocity to arrive at the trawler as soon as possible. Wandering dolphins set their velocity from a random-gamma distribution to approximate field data (see *dolphin-select-velocity* submodel for details).
b) *dolphin-move:* The dolphin actually moves in these procedures. Choices are:
   - *dolphin-travel-to-trawler:* Dolphin sets its heading for the trawler and moves toward it at maximum velocity.
   - *dolphin-flock-with-trawler:* Dolphin stays close to an actively fishing trawler and matches the trawler's speed and heading. Maximum flocking time limit is randomly selected between *dolphin_min_flocking_time.*

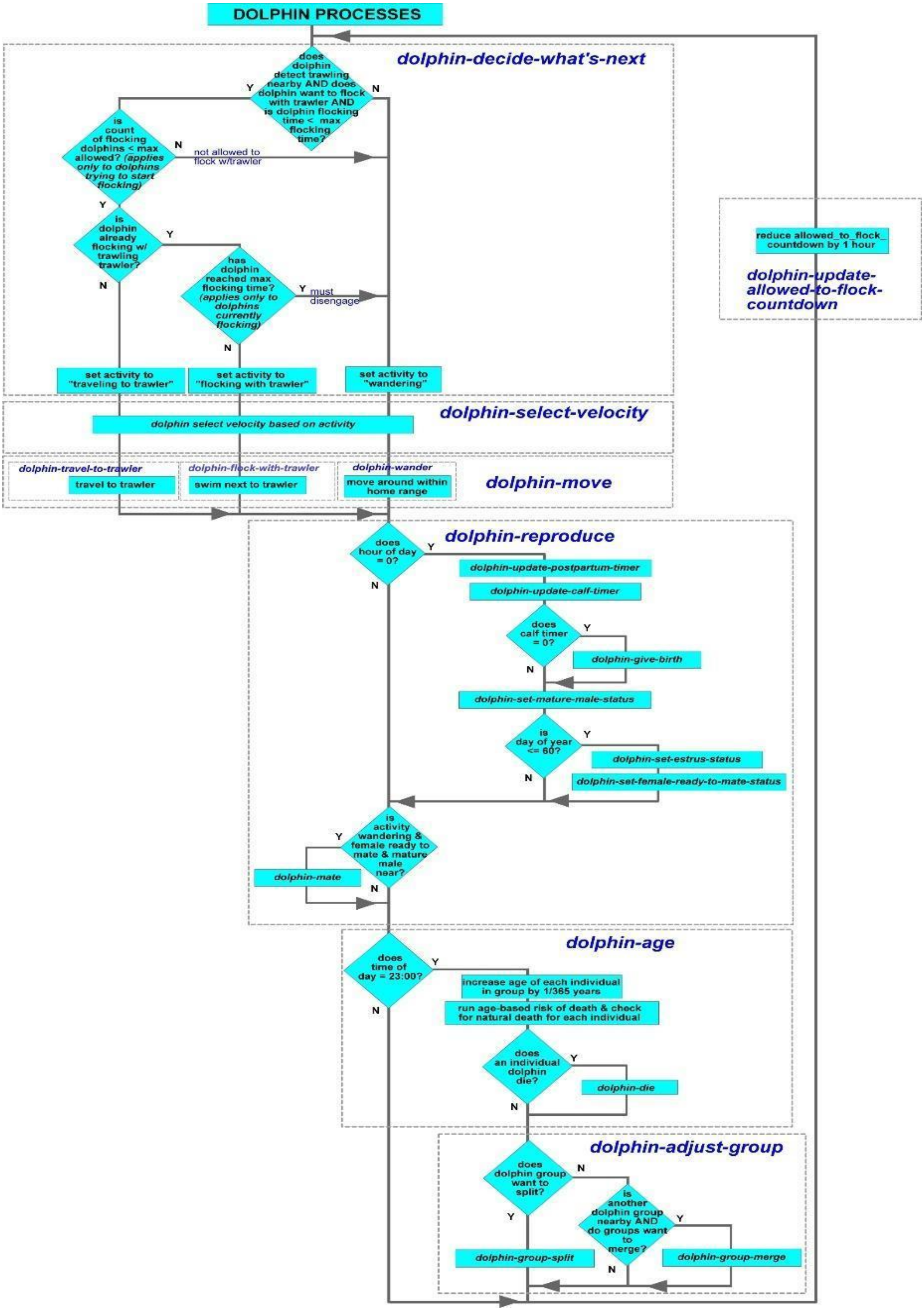

**Figure B.23** Dolphin Processes Flowchart.

- *dolphin_max_flocking_time* (hours) sliders on the UI. Once a dolphin reaches that limit, it moves away from the trawler and returns to wandering. It selects a countdown interval that prevents it from flocking with another trawler randomly between the *dolphin_min_interval_between_flocking* and *dolphin_max_interval_between_flocking* (hours) sliders.
- *dolphin-wander:* Random vector-biased walk that replicates the dolphin swimming within its home range defined by the range's shoreline length and maximum depth limit. See detailed description in *dolphin-wander.*

c) *dolphin-reproduce:* This procedure oversees all the dolphins' reproductive processes. Every day at midnight the dolphins do three things:

- *dolphin-update-postpartum-timer:* decrement each female dolphin's postpartum timer by one day
- *dolphin-update-calf-timer:* decrement each pregnant dolphin's gestation countdown timer by one day and manage birth when the calf timer reaches 0 (*dolphin-update-calf-timer*)
- *dolphin-set-mature-male-status:* updates whether there are any males > six years old in the group

If within the first 60 days of the year, these procedures run to manage female dolphins' reproductive status:

- *dolphin-set-estrus-status:* female dolphins enter and leave estrus according to their estrus start days
- *dolphin-set-female-ready-to-mate-status:* updates whether there are females in the group that are ready to mate which requires at least one female that is: not pregnant, in estrus, and satisfied their postpartum interval. Model female dolphins do not experience reproductive senescence since there is no evidence from field data (Slooten and Lad 1991).

- if there is a female dolphin ready to mate and her activity is wandering, she has the opportunity to mate (mating is blocked while traveling to or flocking with a trawler). See Section 7.2.4.14 for detailed description of dolphin-mate.

d) *dolphin-age* (and natural death): Every day at 23:00 each individual dolphin's age in every group increases by one day. This is also the time the model runs an age-based probability of an individual dolphin dying a natural death. The model has a list of 31 daily mortality probabilities (Gormley 2009; Wickman 2024); one for each dolphin age from <1-31 years, with dolphins that turn 31 having a mortality probability of 1. If a random uniform draw between 0-1 comes back as < daily probability, that specific dolphin in the group dies. When that happens, that individual is removed from the group and group size is updated. If it's the last dolphin in that group, the dolphin group is removed completely from the simulation.

e) *dolphin-adjust-group:* Field observations show groups of dolphins periodically merge to form a new, larger group and also split up into smaller groups in "fission/fusion" behavior (Slooten et al. 1993; Slooten 1994; Dawson et al. 2004; Rayment et al. 2009; Turek et al. 2013). Model dolphin groups get that same opportunity once per day. The odds of a group splitting increase as the group size increases (see *dolphin-group-split* procedure). If the group does not split and there is another dolphin group of the same subspecies nearby, it has the opportunity to merge with that group to form a larger group with probability based on the combined group size *(dolphin-groups-merge).*

f) *dolphin-update* allowed to flock countdown: This procedure runs after all the other activities in the hour have completed. For those dolphins prevented from flocking with a trawler because the time between flocking events has not expired (one of the constraints in *dolphin-decide-what's-next),* this procedure reduces that time by one hour until it reaches zero.

*7.4.2 Detailed Dolphin Submodels*

*7.4.2.1 dolphin-select-velocity*

Dolphins select their velocity depending on their next activity. Dolphins flocking with trawlers set their velocity to match the trawler's. Dolphins traveling to trawlers swim at maximum velocity to arrive at the trawler as soon as possible. If the dolphin decides there are no trawlers around to flock with, it picks its wandering velocity based on a gamma distribution approximated from field data (Slooten et al. 1993; Slooten 1994; Rayment et al. 2009) as shown in Figure B.24. Final velocity is the maximum of either 0.45 km/hr or the result of the random-gamma draw formula.

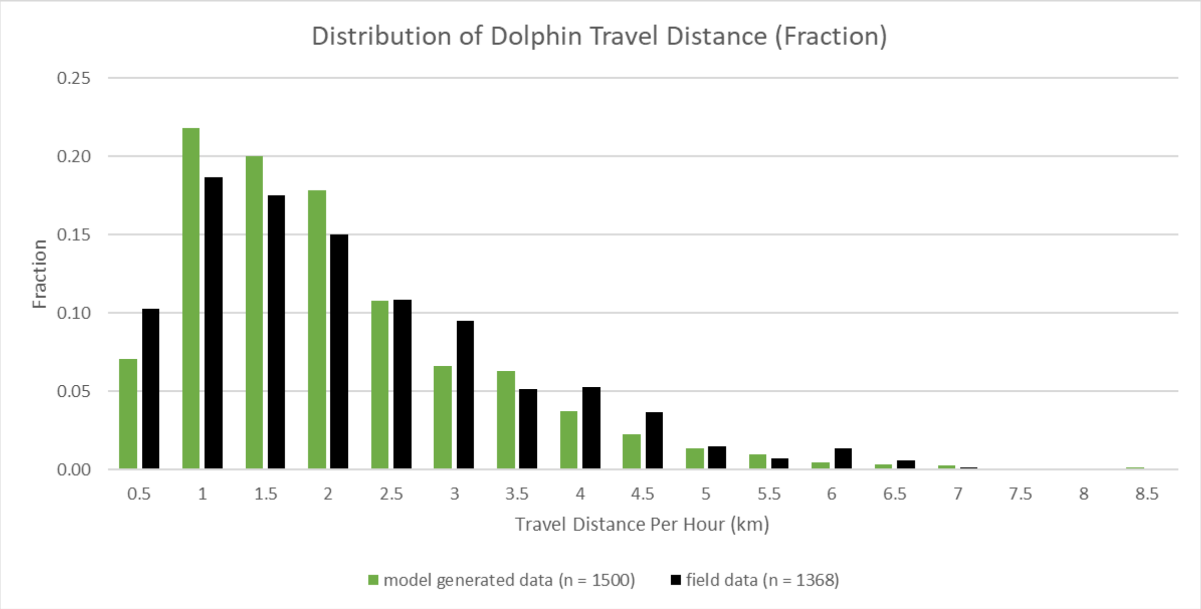

**Figure B.24** Dolphin Travel Distance Distribution.

The formula for selecting the velocity for a full hour of wandering is:

$$velocity\_km\_per\_hour = max\ (list\ 0.45\ precision(\ (1.75\ +\ (random - gamma\ 1.5\ 1.0)\ -\ 1.5))\ 2)$$

The dolphin converts this from km/hr to patches/tick for the actual movement phase. If a dolphin enters this procedure mid-tick, it inherits the velocity it is allowed to use in the time remaining in the tick.

*7.4.2.2 dolphin-wander*

These steps describe the dolphin wandering (any time not traveling to or flocking with a trawler):

1. Select Wandering Heading: The dolphin selects a new wandering heading each time it enters this procedure which could be at the start of a tick or somewhere in the middle if it is flocking with a trawler and the trawler stops fishing. It is a multi-step process that involves creating two biasing vectors that steer the dolphin to keep it within its home range.
   a. It picks the initial heading as current heading plus a random normal value ($\mu$ = 0 and $\sigma$ = 20 degrees).
   b. Next it projects the actual xcor and ycor it will end up in if it moved based on the initial heading and wandering velocity (in patches/tick).
   c. It calculates the difference in xcor and ycor from current to projected coordinates
   d. It takes the delta xcor and delta ycor and builds an unbiased vector $\upsilon_{unbiased}$ to the projected coordinates; this is how the dolphin would travel without the influence of the biasing vectors.
   e. Next it builds the first biasing vector $\upsilon_{biasing\_midline}$:
      i. First it finds the closest ocean patch that runs through the midline of the home range perpendicular to the home range orientation. Shown below in Figure B.25 is an example of midline patches for an east-west home range (gray patches):
      ii. Compute heading back to the closest midline patch from projected xcor and ycor
      iii. Convert that heading into an angle in Netlogo geometry
      iv. Compute distance back to closest midline patch using either the projected xcor (for dolphins with home ranges that run predominantly east to west) or ycor for dolphins with north-south

home ranges (see *setup-shoreline-patches-homerange-orientation* procedure for more details)

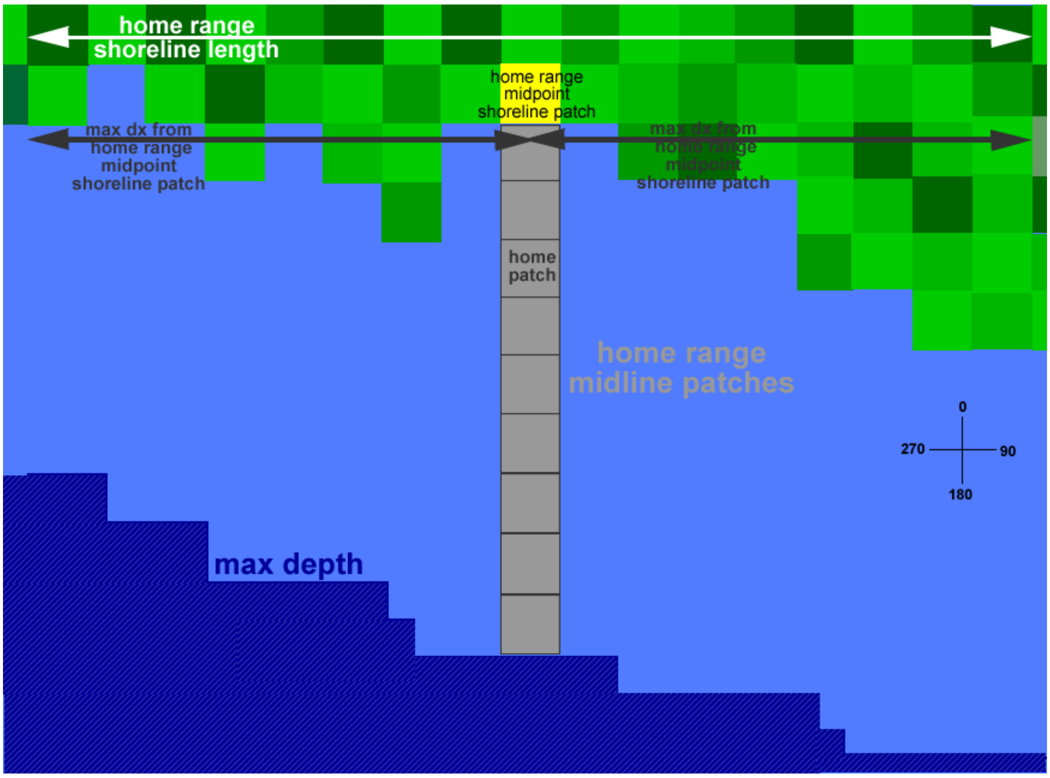

**Figure B.25** Midline patches for east-west dolphin home range.

v. Calculate length of midline patch biasing vector as:

$$L = 1/(d - \ distance\ to\ midline\ patch)$$

$$Where\ d = \frac{1}{2}\ the\ home\ range\ shoreline\ length$$

Note that if the dolphin wanders outside of its home range, the distance to midline patch would be > d and L would be negative. If that happens, the calculated L is replaced by 1.0, which will steer the dolphin back towards the midline patch

vi. Construct biasing vector υbiasing_midline based on L and heading back to nearest midline patch

$$\upsilon\_biasing\_midline \ = (L \ * \ cos\ angle, L \ * \ sin\ angle)$$

f. The dolphin follows a similar process using the projected depth to build a depth biasing vector that prevents it from exceeding maximum depth. As depth increases the strength of the bias toward shallow water increases:

   i. Record depth of patch that contains projected xcor and ycor
   ii. Compute heading to dolphin's home range midpoint shoreline patch from projected patch and convert to Netlogo angle; we assume dolphin would head towards shore (and shallow water) if it found itself in deep water
   iii. Calculate depth ratio as:

$$Depth\ ratio\ =\ projected\ depth\ /\ max\ allowed\ depth$$

   iv. Calculate length of depth biasing vector using logistics curve as shown in Figure B.26 (chosen with a very steep k factor so shallow depths have no influence on final heading, but projected depths close to maximum exert very strong influence):

$$L\ = value\ of\ curve\ /(1 +\ \ exp\ (-k)(depth\ ratio - midpoint)))$$

Where: max value = 1.1, k = 15, midpoint = 0.95

   v. Construct biasing vector ʋbiasing_depth based on L and angle back to shore patch:

$$\upsilon\_biasing\_depth\ \ =\ (L\ *\ cos\ angle, L\ *\ sin\ angle)$$

g. Finally, takes the two biasing vectors ʋbiasing_midline and ʋbiasing_depth and adds them to ʋunbiased to get the final biased vector ʋbiased , and uses the atan primitive to convert the angle to a final heading.

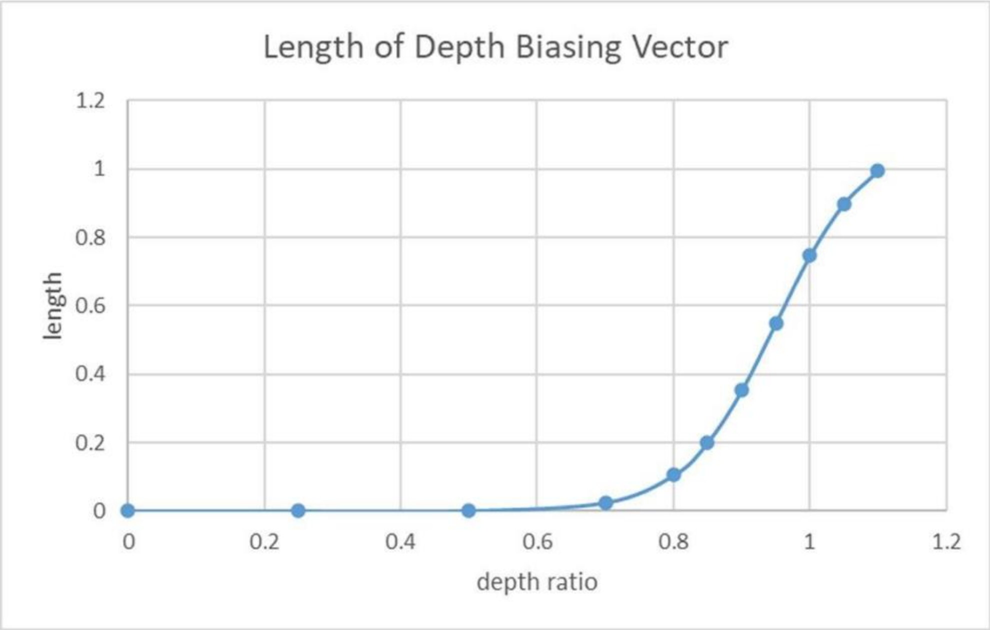

**Figure B.26** Length of depth biasing vector.

Two examples of the biasing vector calculation follow. Figure B.27 shows a dolphin (black object) near shore, close to the midline patches (light gray) within its home range (light blue patches).

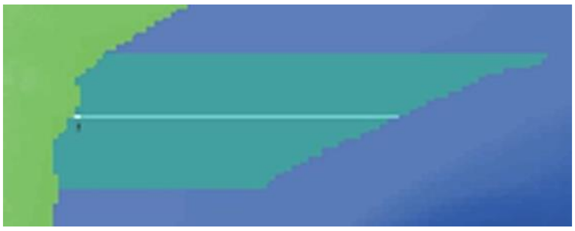

**Figure B.27** Dolphin location in home range for biasing vector example 1.

The dolphin is asked to pick a wandering heading and results are:

- initial (unbiased) heading = 25.52 deg
- υunbiased = [0.2973 0.6227]
- distance of projected patch from closest midline patch dmidline patch if dolphin moved based on unbiased heading and already-selected velocity = 1.3 patches
- maximum distance from middle of home range to travel and stay within home range d = 14.52 patches
- Heading from projected patch to closest midline patch = 347.82 deg
- L for midline patch biasing vector = 0.076

- υbiasing_midline = [-0.016 0.074]
- depth of projected patch = 14.19 m
- ratio of projected depth to max allowed depth (depth ratio) = 0.1419
- heading from projected patch to shoreline midline patch = 300.95 deg
- L for depth biasing vector = 0.0038
- υbiasing_depth = [-0.0033 0.002]
- υbiased = [0.278 0.699] which converts to final heading = 21.68 deg

Figure B.28 shows the various vectors. The υbiasing_midline vector wants to steer the dolphin to the NNW, but since this dolphin is very close to the midline of its home range, its influence is small. The υbiasing_depth vector is even less influential because the dolphin is already in shallow water. The final heading reflects the slight influence of both steering vectors as the dolphin makes a very minor adjustment from 25.52 deg (NNW) to a 21.68 deg heading

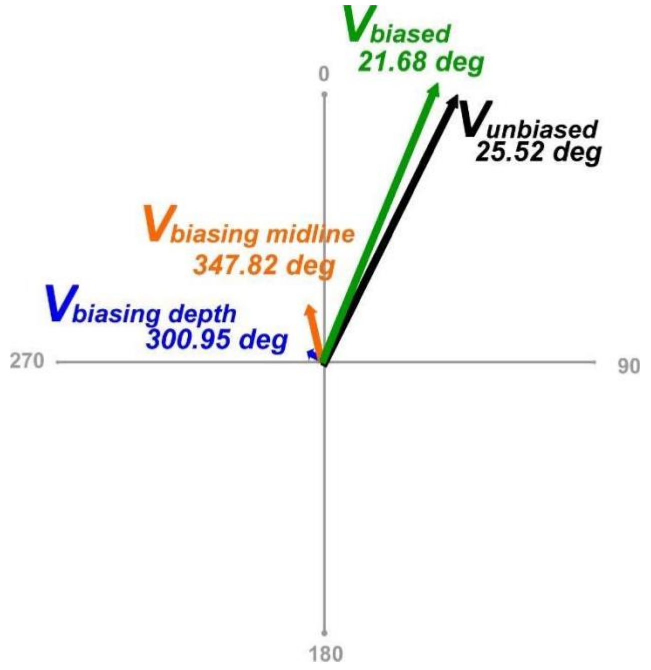

**Figure B.28** Example 1 of unbiased vector, biasing vectors, and final vector.

The second example is shown in Figure B.29; the dolphin is still within its home range, but near the depth limit and slightly more than halfway to its northernmost home range travel limit.

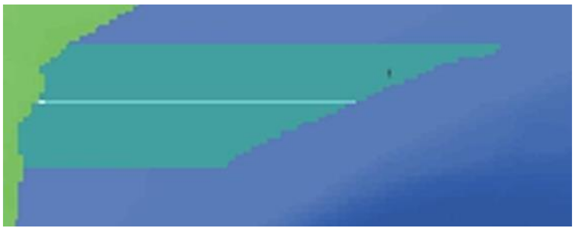

**Figure B.29** Dolphin location in home range for biasing vector example 2.

The dolphin is asked to pick a wandering heading and results are:

- initial (unbiased) heading = 29.83 deg
- υunbiased = [0.34327 0.5985]
- distance of projected patch from closest midline patch dmidline patch if dolphin moved based on unbiased heading and already-selected velocity = 7.6 patches
- maximum distance from middle of home range to travel and stay within home range d = 14.52 patches
- Heading from projected patch to closest midline patch = 182.59 deg
- L for midline patch biasing vector = 0.144
- υbiasing_midline = [-0.00652 -0.1443]
- depth of projected patch = 81.93 m
- ratio of projected depth to max allowed depth (depth ratio) = 0.8193
- heading from projected patch to shoreline midline patch = 264.85 deg
- L for depth biasing vector = 0.314
- υbiasing_depth = [-0.3133 -0.0282]
- υbiased = [0.0235 0.426] which converts to final heading = 3.16 deg

Figure B.30 shows the various vectors. Since this dolphin is past half way on distance from the midline of its home range, the υbiasing_midline vector does want to steer dolphin to the south, but it is not exerting a strong steering input. The υbiasing_depth vector is more influential, steering the dolphin west and back to shallow water. The final heading reflects the influence of both steering vectors as the dolphin turns away from 29.83 deg (NNW) to a 3.16 deg heading.

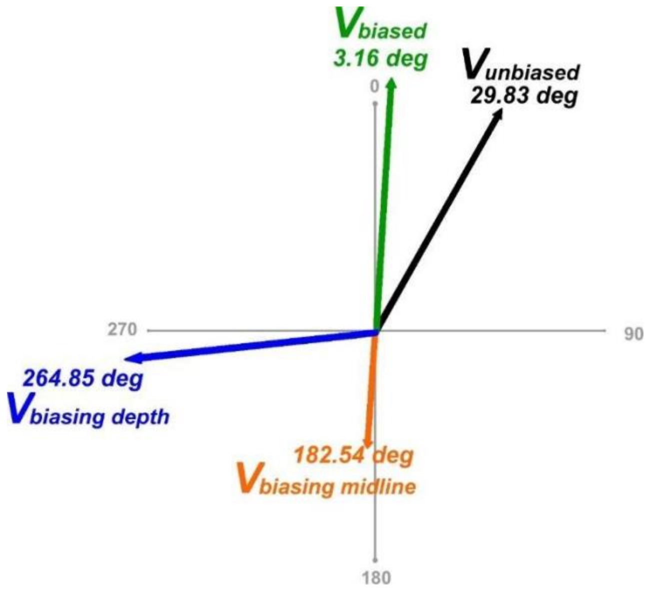

**Figure B.30** Example 2 of unbiased vector, biasing vectors and final vector.

2. Checks for land patch: Before the dolphin actually moves, it looks ahead; if there is a land patch <= one-half patch ahead, it will stop and set a new heading approximately parallel to shore, then continue to move until the dolphin has travelled the total distance it should travel for that hour (based on velocity selected in dolphin-select-velocity procedure).
3. Divide velocity into mini-steps: Because of the dolphins' proximity to land, they cannot use up the desired velocity (i.e., travel distance) in one move; otherwise, the dolphin could end up several patches inland. Instead, we divide the full hour's travel velocity into 5 mini-steps; e.g., if velocity is 4 patches/tick (4 patches/hour), the mini-step = 0.8 patch.
4. Move forward one mini-step
5. Calculate remaining velocity: Dolphin subtracts the mini-step velocity from the full velocity to calculate remaining patches/tick it can travel. In our example, the dolphin just moved 0.8 patch and therefore has 3.2 patches left to travel in this tick (hour).
6. Check for gillnets: If there is a gillnet in this patch, dolphin puts it in a running list that tracks gillnets encountered this tick; duplicates are not added to the list

7. Decide if migrating to a new area: if it wanders in a patch adjacent to a dolphin area different from its current one, the dolphin could switch areas. The frequency of switching is controlled via an input slider on the UI and reflects the fraction of the entire population that is expected to migrate in a year. The actual odds of migrating are calculated based on: the current population, the number of migrations that have already occurred this year, and the projected opportunities to migrate based on dolphin movement so far. See dolphin-calculate-migration-odds procedure section for detailed description of the migration odds calculation. If a random uniform number draw is < the probability, the dolphin migrates. If the dolphin does migrate to a new area, it will pick a new home patch, home range shoreline midpoint patch, and home range centered on that patch.
8. Check to see if wandered onto a land patch: Just in case the look-ahead fails, move to closest ocean patch and turn parallel to shoreline
9. Repeat steps 4-8: Dolphin continues to move forward in mini-steps until the remaining velocity = 0, which means it has travelled the correct distance.

*7.4.2.3 dolphin-detect-trawling-nearby-and-decide-to-travel*

Dolphins that are alerted by a trawler that a trawl has started/continuing from the previous tick via trawler-alert-dolphins-trawling-happening run this procedure to ultimately decide if they will move towards the trawler. In this procedure each dolphin group:

1. Determines the actual trawl noise that reaches them based on the current distance to trawler using the dolphin-calculate-trawl-noise-level procedure
2. Uses that noise and a probability curve to determine the probability an adult in the group detects the trawl via dolphin-calculate-probability-of-detecting-trawl procedure
3. Assigns each adult in the group a random-uniform number from 0-1
4. If any adult's random number is less than the detection probability, that individual adult dolphin detected the trawl, and we assume the adult informs the entire dolphin group that the trawl was detected. If no adults detect the trawl, the dolphin group will continue to wander.

5. If the trawl is detected, the dolphin calculates the probability of moving towards the trawler via dolphin-calculate-probability-of-traveling-to-trawler procedure
6. Finally, the dolphin group picks a random-uniform number from 0-1. If the random number is < travel probability, the dolphin has decided to travel to the trawler. If not, the dolphin group will continue to wander within its home range.

*7.4.2.4 dolphin-calculate-trawl-noise-level*

In this procedure, the dolphin calculates the actual trawl noise received based on trawler's trawl noise and distance to trawler. It calculates spreading transmission losses based on actual distance from trawler (R) via:

$$TL_{spreading} = TL\_equation\_coefficient * logR$$

Traditionally, 20 is used for the TL_equation_coefficient in deep water (spherical spreading) and 10 is used in shallow water (cylindrical spreading). In our case, trawling is carried out in intermediate depths so we used 15 as the TL_equation_coefficient, also known as "intermediate" spreading (Lynch and Newhall, 2017). The dolphin then calculates absorption transmission losses via:

$$TL_{absorption} = alpha\_broadband * log\ R$$

$$where\ alpha\_broadband\ is\ broadband\ absorption\ coefficient\ in\ seawater\ 1.273e - 05\ dB/m$$

Finally, the actual trawl noise received by the dolphin is calculated as:

$$RL\ (dB) = Trawl\ noise - TL_{spreading} - TL_{absorption}$$

*7.4.2.5 dolphin-calculate-probability-of-detecting-trawl*

This procedure determines the probability a dolphin detects a trawl based on the trawl noise that reaches the dolphin and an estimated probability curve as shown in Figure B.31. We used Figure 1 from Kastelein et al. (2017) to estimate 100 dB as the received level of noise that gives a dolphin a 50% chance of detecting a trawl. From there it is a straight-line regression to 200 dB. From 200 dB on, the probability of detection is 1.0.

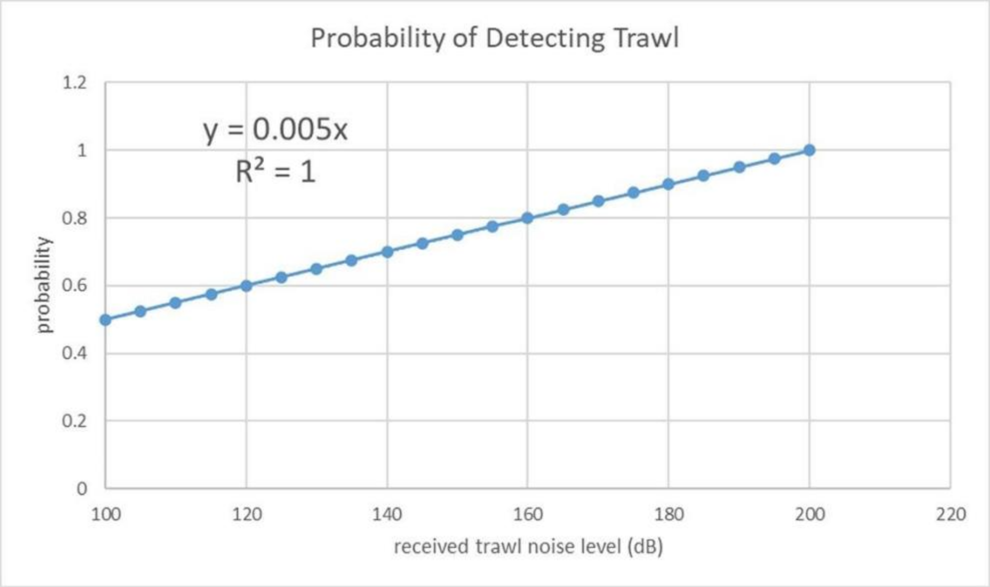

**Figure B.31** Dolphin probability of detecting trawl regression.

*7.4.2.6 dolphin-calculate-probability-of-traveling-to-trawler*

This procedure determines the probability that a dolphin group will travel to the trawler given the distance to the trawler per the following regression curve in Figure B.32:

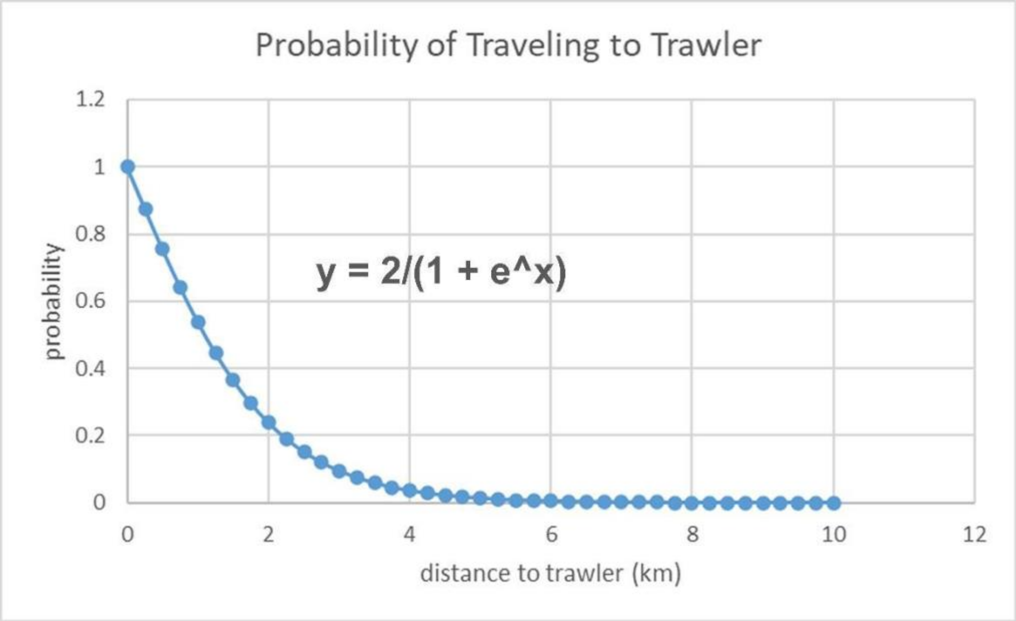

**Figure B.32** Probability dolphin group travels to trawler.

*7.2.4.7 dolphin-calculate-migration-odds*

This procedure runs when the dolphin group is deciding whether to switch areas (migrate). It calculates the specific probability the dolphin group will migrate based on field data (Fletcher et al., 2002). The calculation steps are:

1. Calculate expected migrations/year based on total dolphin count:

$$expected\ migrations\ per\ year = \ count\ dolphins * fraction\ of\ dolphins\ migrating/yr$$

2. Calculate expected number of total migrations at the end of current simulation year:

$$expected\ total\ migrations = expected\ migrations\ per\ year * year\ of\ sim$$

3. Estimate daily opportunities to migrate based on total opportunities so far (the model counts and retains each time a dolphin has the opportunity to migrate):

$$migration\ opportunities\ per\ day = \frac{total\ migrations\ opportunites}{day\ of\ sim}$$

4. Calculate remaining expected migrations to achieve expected total migrations at end of current year:

$$remaining\ expected\ migrations$$
$$= expected\ total\ migrations - count\ actual\ dolphin\ migrations$$

5. Estimate remaining number of opportunities to migrate this year:

$$remaining\ migration\ opportunities\ this\ year$$
$$= (365 - day\ of\ year) * migration\ opportunities\ per\ day$$

6. Calculate migration probability based on how many migrations are still needed to reach the expected total at end of year considering the remaining opportunities for the rest of the year:

$$migration\ probability = \frac{remaining\ expected\ migrations}{remaining\ migration\ opportunities\ this\ year}$$

For example, assume we are in day 500 of the sim (i.e., day 135 of year 2), there are 1000 dolphins, there have been 65 total migrations so far, there have been 100,000 total opportunities to migrate so far, and the expected migration rate is 5%/year. The migration probability calculation is as follows:

1. Expected migrations/year = 1000 * 0.05 = 50
2. Expected total migrations at end of year 2 = 50 * 2 = 100
3. Migration opportunities/day = 100,000/500 = 200
4. Remaining expected migrations for the rest of year 2 = 100 – 65 = 35
5. Remaining migration opportunities this year = (365 – 135) * 200 = 46,000
6. Migration probability = 35/46,000 = 7.61e-4

*7.2.4.8 dolphin-update-postpartum-timer*

The postpartum timer represents the two-to-four-year interval between births as observed in field data (Slooten 1991; Gormley 2009; Bennington 2025). For those mature females who have given birth and are still waiting for their postpartum timer to elapse so they can enter estrus and reproduce, this procedure reduces the timer by one day, but does not decrease it below 0.

*7.2.4.9 dolphin-update-calf-timer*

For those female dolphins that are pregnant with gestation time remaining, this procedure reduces the calf timer by one day. When the calf timer reaches 0, gestation is complete which kicks off the dolphin-give-birth procedure.

*7.2.4.10 dolphin-give-birth*

This procedure adds a new individual dolphin to the group with a random 50-50 choice on gender. It adjusts all the dolphin group's attributes (e.g., age list, group size, etc.) to account for the new calf. It also adds the calendar year to the mother's list of years she has given birth, calls dolphin-set-postpartum-timer procedure to pick this female's minimum interval between births, updates the total calves by subspecies metric, and increments the patch attribute that tracks number of dolphins born in that particular patch.

*7.2.4.11 dolphin-set-mature-male-status*

If there is a male > six years old in the group, it sets the group attribute to true; otherwise, it is false. This makes it easier for a female to check if a dolphin group has a mature male to mate with.

*7.2.4.12 dolphin-set-estrus-status*

This procedure checks each individual female dolphin to see if she is > six years old and the day of the year is within one of her seven-day estrus periods. If so, it sets her estrus status to true; otherwise, it is false.

*7.2.4.13 dolphin-set-female-ready-to-mate-status*

This procedure checks to see if there is a female dolphin in the group that is: in estrus, not currently pregnant, and has satisfied her postpartum interval. If all are true for at least one female, it sets the group attribute to true; otherwise, it is false. Groups with a female ready to mate can actually run the dolphin-mate procedure (although only the individual female(s) that satisfy all three criteria can actually become pregnant).

*7.2.4.14 dolphin-mate*

The dolphin-mate procedure flowchart is shown in Figure B.33. If there is at least one mature male of the same subspecies within the detection distance of a female dolphin, they will mate (note that the dolphins do not actually move to each other in the simulation). The baseline odds that mating will result in pregnancy are controlled via an input slider on the UI (Slooten 1991; Slooten and Lad 1991; Gormley 2009). That probability is adjusted based on the age of the female dolphin with dolphins younger than eight years old having less-than-baseline odds. The model does a random uniform draw and if that number is < pregnancy probability, the female becomes pregnant and a gestation time from 300-365 days is selected from a random normal distribution with $\mu = 330$ and $\sigma = 10$ (via dolphin-select-gestation-days procedure). This gestation time ensures calf delivery in November-February to match field data on calving (Slooten 1991; Gormley 2009). Lastly, pregnant female dolphins set their estrus status to false. Note that female dolphins do not experience reproductive senescence since there is no evidence from field data (Slooten and Lad 1991; Gormley 2009).

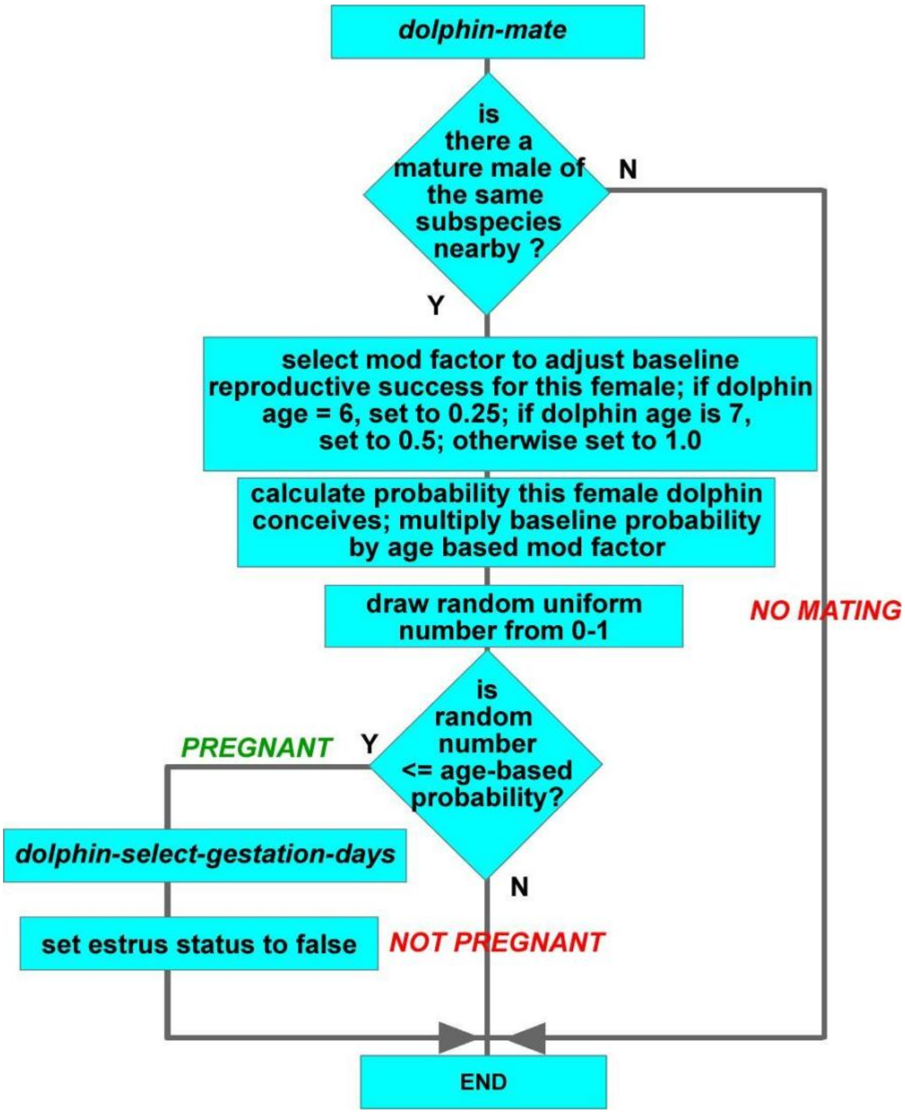

**Figure B.33** *Dolphin-mate* procedure flowchart.

*7.2.4.15 dolphin-age*

This procedure runs once per day and ages every individual dolphin by one day. It also does a random number draw for each individual and compares it to the age-based daily mortality risk. If the random number is < daily probability of dying or dolphin turns 31 years old, the individual dolphin dies. When that occurs, the individual is removed from the group and group attributes (e.g., age list, group size, etc.) are adjusted via the dolphin-die procedure.

*7.2.4.16 dolphin-set-postpartum-timer*

This procedure runs immediately after a dolphin gives birth and sets a timer that prevents the dolphin from mating until the timer counts down to 0. The timer expires on January 1 of the next eligible birth year so the female can come into estrus. To set the timer, the model:

1. Calculates days left in year; e.g., if dolphin gives birth on Nov 14, 2022 (day 318), there are 47 days left in the year
2. Picks the calf interval from 2-4 years based on a triangular distribution; e.g., if model picks a three-year interval, this dolphin cannot give birth again until 2025
3. Calculate number of days in postpartum interval as:

$$Postpartum\ Interval = ((yearly\ interval - 1) * 365) + days\ remaining\ in\ current\ year$$

e.g., ((3 – 1) * 365) + 47 = 777 days

*7.2.4.17 dolphin-group-split*

Dolphin groups have the opportunity to split into two smaller groups once per day at 23:00. The probability of splitting is based on the current group size. Maui dolphins tend to form smaller groups; therefore, the splitting and merging probabilities are different for the two sub-species. Dolphin group split probability is determined based on the following logistics growth equation with different parameters for each subspecies:

$$Pgroup\ split\ with\ n\ members\ =\ P\ +\ (L\ *\ P\ *\ (1\ -\ P/K)$$

$$where:$$

$$P\ =\ probability\ of\ group\ splitting\ with\ (n\ -\ 1)\ members$$

$$splitting\ probability\ K\ = 1$$

$$L\ =\ curve\ shape\ factor$$

Each curve starts with the probability of a group with two dolphins splitting, and then uses the equation for the remaining group sizes. For Hector’s dolphins, the split probability for a group of two = 0.2 and shape factor L = 0.1, which generates the probability curve shown in Figure B.34.

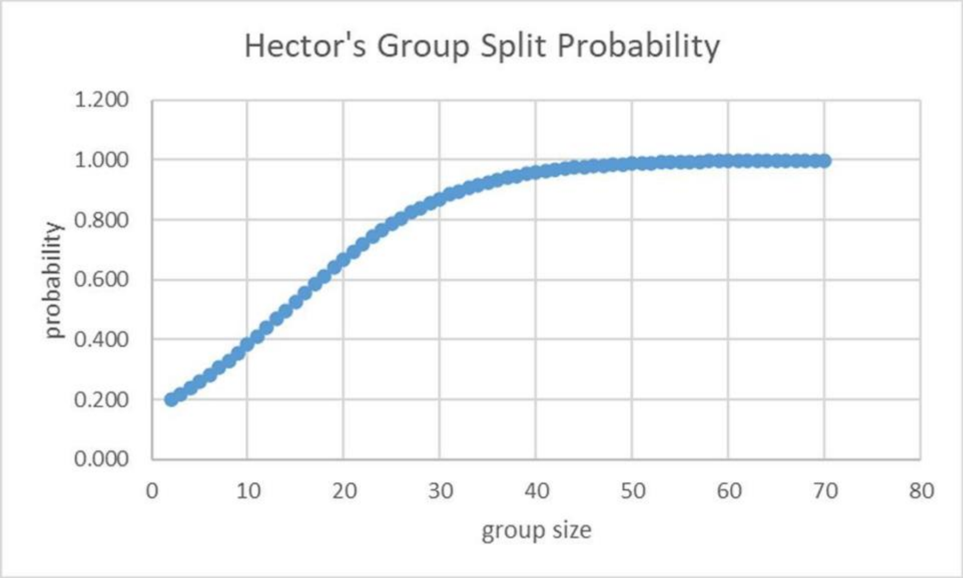

**Figure B.34** Splitting probabilities for Hector's dolphin groups by group size.

For Māui dolphins, the split probability for a group of two = 0.2 and L = 0.4, which generates the split probability curve shown in Figure B.35.

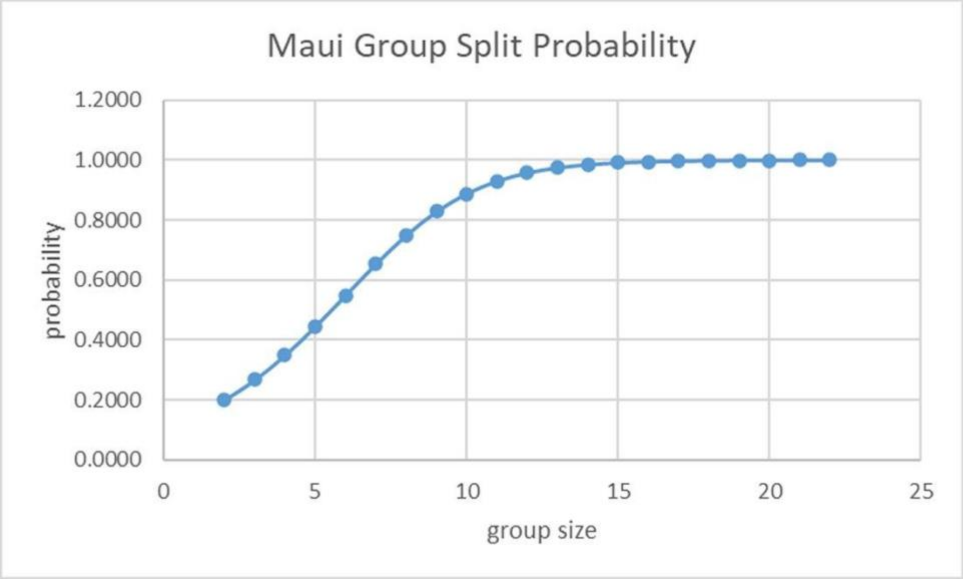

**Figure B.35** Splitting probabilities for Maui dolphin groups by group size.

The model does a random uniform draw between 0 and 1, and if the number is < split probability, the group splits into two dolphin groups, with the “breakaway” group taking a random number of individual dolphins away from the original group.

*7.2.4.18 dolphin-groups-merge*

Merges can only happen if the group initiating the merge has not split today and another dolphin group of the same subspecies is within the detection distance. The probability of merging is based on the combined size if the two groups actually merge and is different between Hector's and Māui dolphins because their observed group sizes in the wild are different (Dawson et al. 2004; Slooten et al. 2004, 2006a; Rayment et al. 2009; Turek et al. 2013; MacKenzie and Clement 2014, 2016, 2019; Constantine et al. 2021).

The probability of a dolphin group merging with another group of the same sub-species is determined via the following exponential decay equation with different parameters for each subspecies:

$$Pgroup\ merge\ with\ n\ total\ members\ =\ P - (L * P)$$

$$where:$$

$$P\ =\ probability\ of\ groups\ merging\ with\ (n\ -\ 1)\ total\ members$$

$$L\ =\ curve\ shape\ factor$$

Each curve starts with the probability of two single-member groups merging to form a single group of two. For Hector's dolphins, the merge probability to create a group of two = 0.35 and shape factor L = 0.083, which generates the merging curve shown in Figure B.36:

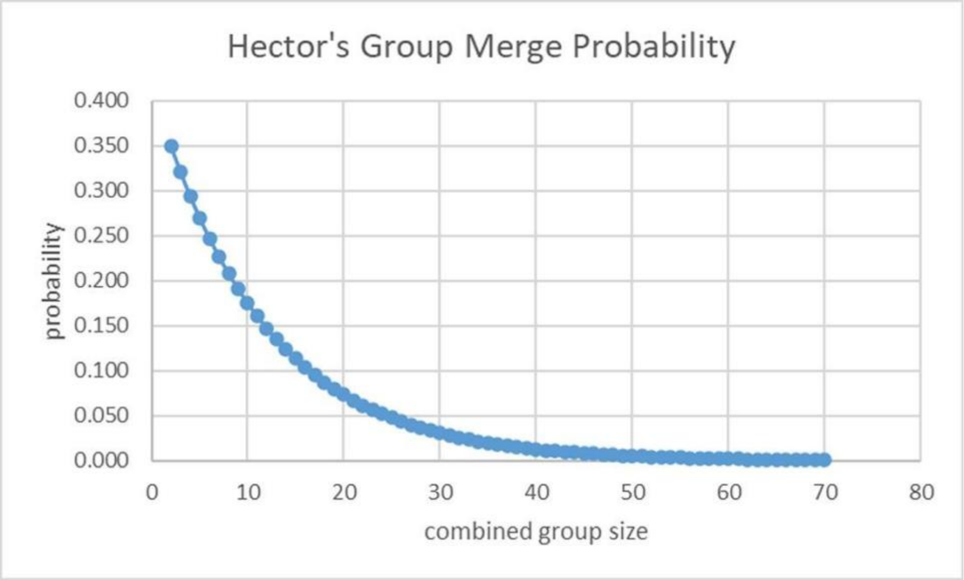

**Figure B.36** Merging probabilities for Hector's dolphins by group size.

For Māui dolphins, the merge probability to create a group of two = 0.75 and L = 0.297, which generates the merging curve in Figure B.37. A random uniform number drawn between 0 and 1 determines if the merge occurs.  If it does, the dolphins of the "merging" group are folded into the other group to build the larger group.

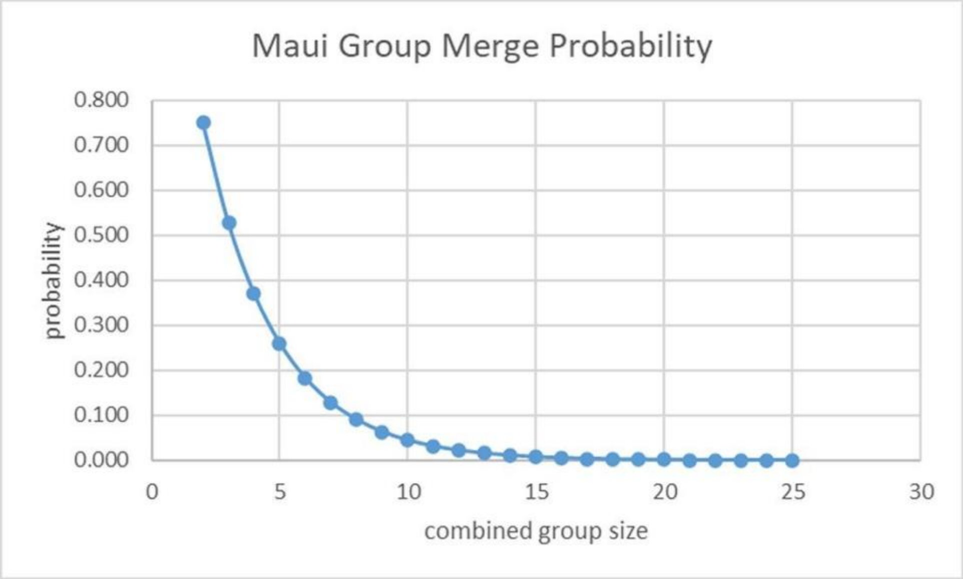

**Figure B.37**  Merging probabilities for Maui dolphins by group size.

*7.5 Gillnet Submodels*

*7.5.1 Overview of Gillnet Processes*

Figure B.38 is an overview of gillnet processes.  Every day at 23:00, the gillnets-relocate procedure runs.  That kicks off a series of procedures:

a) *increment-total-gillnet-hauls:* With the gillnets relocating, the model counts up the total number of deployed gillnets and adds that amount to the model parameter total_gillnet_hauls

b) *gillnet-select-next-patch:*  Each gillnet selects its next fishing location from ocean patches that allow gillnet fishing with patches that have greater historical fishing effort given a higher probability

c) each gillnet moves to its new patch

d) *gillnet-decide-if-deployed:* Depending on the goal gillnet fishing days for the year and how many gillnets exist, the gillnet may or may not be "deployed", which means the gillnet is eligible to kill dolphins. Gillnets that are not deployed are inactive as far as the model is concerned; they do not contribute to the total number of gillnet hauls or gillnet fishing days, and they cannot kill dolphins. Gillnets use this procedure to set their deployed? attribute to true or false.

As dolphins wander, they occasionally encounter a gillnet and risk being killed (we define an encounter as any time a dolphin group and deployed gillnet are in the same patch). Those encounters trigger the gillnet-kill-dolphins procedure which does the probability checks that an individual dolphin within the group is killed by the gillnet.

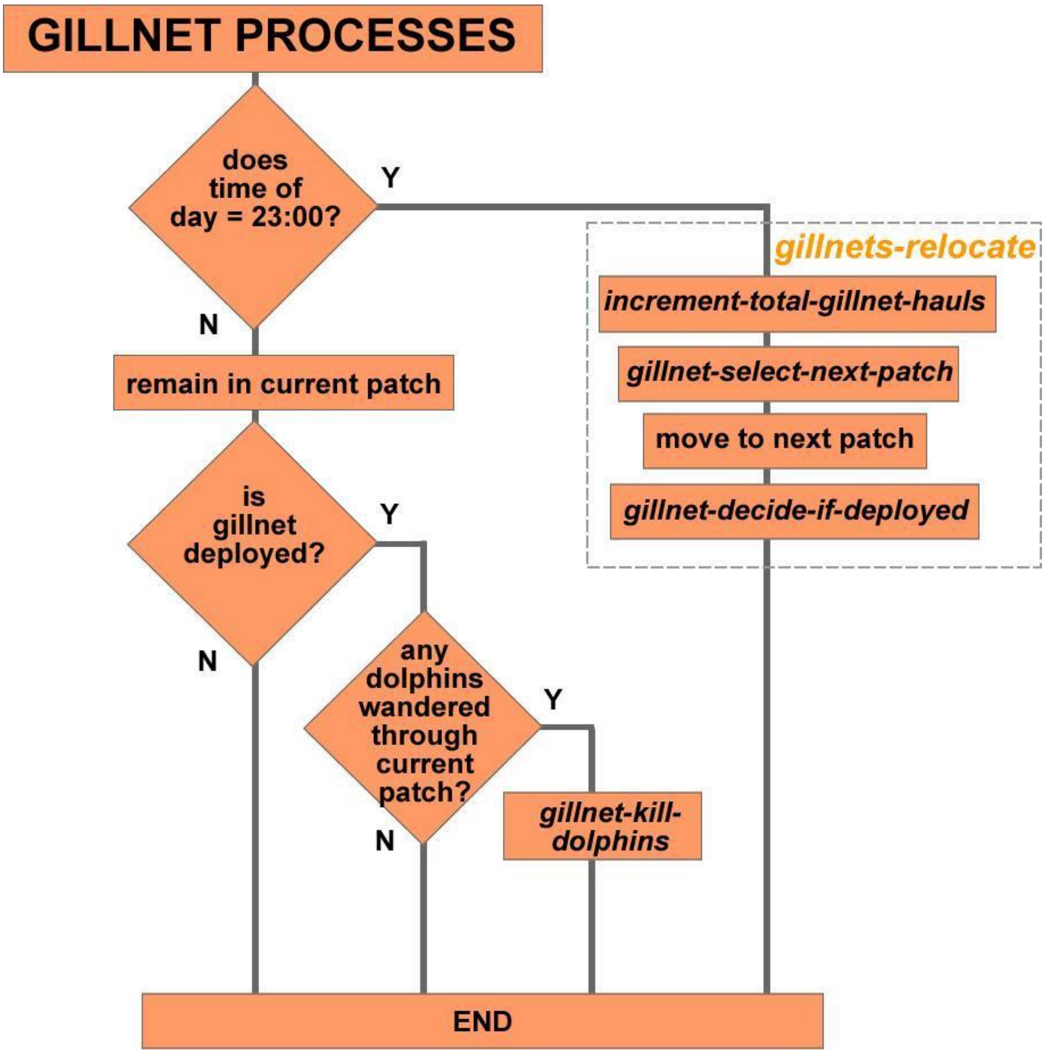

**Figure B.38** Gillnet Processes Flowchart.

*7.5.2 Detailed Gillnet Submodels*

*7.5.2.1 increment-total-gillnet-hauls*

The model counts up the total number of deployed gillnets and adds that amount to the model parameter total_gillnet_hauls.

*7.5.2.2 gillnet-select-next-patch*

Gillnets choose their next patch based on a random weighted draw from unprotected gillnet patches in their placement area based on historical fishing effort data provided by MPI (2023, 2024). These data are stored as patch attribute gillnet_gradient, with more popular areas having a higher probability of being selected.

*7.5.2.3 gillnet-decide-if-deployed*

This procedure reads the gillnet_deployment_probability_by_area_table to get the probability each gillnet will be deployed that day in their placement area. For example, assume 500 gillnet fishing days is the target for the area. The model has already calculated and created the minimum integer number of gillnets needed to get 500 deployed days in a year (500/365 = 1.37 which requires 2 gillnets minimum). Since there are more gillnets than needed, the model calculates the deployment probability (500/(2 gillnets * 365 days/yr = 0.68) and stores it in the gillnet_deployment_probability_by_area_table. Each gillnet in the area performs a random uniform draw and if the number is < the probability for its area, the gillnet is deployed.

*7.5.2.4 gillnet-kill-dolphins*

Each time a dolphin group and a deployed gillnet share a patch, there is a chance that individual dolphins will be killed. Gillnets count the number of dolphins in the group and determine the total dolphins killed by running a straight probability draw against the probability of a gillnet killing an individual dolphin. For example, if a group of 20 dolphins passes through a patch with a gillnet and the probability of a gillnet-caused dolphin death is 0.01, the gillnet does 20 random uniform draws from 0-1 and those that came back < 0.01 would count as a “death” and increment the total dolphins killed. The gillnet would then randomly remove individual dolphins from the group until it reached the calculated value for total dolphins killed.  Note that gillnet-dolphin encounters (and therefore bycatch deaths) occur only when dolphins are wandering in patches with deployed gillnets.

#### Supporting information C:

#### Parameter estimation and pattern-oriented modelling (POM)

This appendix describes evaluation and selection of model parameters and compares real world patterns with model-generated patterns (Wiegand et al. 2003; Grimm et al. 2005). We used pattern-oriented modelling (POM; Kramer-Schadt et al. 2007; Grimm et al. 2005) to parameterize the model and confirm model-produced patterns match field data. For most model inputs, data were available for direct parameterization (e.g. 40 years of field data on dolphin movements, home ranges, reproduction, and survival rates). Values for unknown or poorly known parameters were estimated by testing a wide range of values (as in Kramer-Schadt et al. 2007 and Lusseau et al. 2023) using POM to select final parameter values based on overall system behavior.

For example, during development of their model on road mortality of lynx, Kramer-Schadt et al. (2007) varied parameters such as daily movements and the probability of leaving dispersal habitat over a wide range to determine the best fit between observed and simulated patterns. Observed patterns were used as filters to reject parameterizations that did not reproduce observations and to considerably reduce parameter uncertainty. POM allows use of more different kinds of data than a conventional approach, since field data themselves could not be used for parameterization as they were obtained at a higher hierarchical level. This is especially important in situations of scarce data. Kramer-Schadt et al.'s (2007) results illustrate how, despite sparse data, ABMs can be successfully developed, parameterized, and applied to real ecological and conservation problems. They concluded, that "pattern-oriented modelling can be used for detecting the underlying processes that reproduce the observed patterns as well as plausible parameterizations, and inferences can be made from more aggregated data to processes working on a lower level or vice versa" (Kramer-Schadt et al. 2007 p.555). Data are thus not only included from previous studies to construct the model itself, as suggested by Peck (2004), but also contain 'hidden information’ (Wiegand et al. 2003), which can be revealed using POM. Thus, this approach is especially suitable for models in conservation of rare or elusive species, where data generally are scarce and variable (Rossmanith et al. 2007; Wiegand et al. 2003; Kramer-Schadt et al. 2007).

Nabe Nielsen et al. (2014), van Beest et al. (2017) and Lusseau et al. (2023) used POM in their agent-based models studying the responses of harbor porpoise to noise, including noise from pingers in gillnets. Lusseau et al. (2023) showed a nonlinear effect: While high pinger use reduced bycatch, low use increased porpoise movement, increasing bycatch risk from encounters with unprotected nets. This demonstrated the importance of consistent and widespread protective measures. New Zealand dolphins have not shown any response to pingers (Stone et al. 1997; Dawson and Lusseau 2006), therefore protected areas are the main management tool used in New Zealand (e.g. TMP 2019).

##### ***Fisheries data***

Model fishing parameters were based on two sources of New Zealand-specific data:

A. New Zealand Ministry for Primary Industries (MPI 2023, 2024, 2025a,b) - data on the number of trawlers associated with each port, total fishing effort including the total number of trawl and gillnet events and their spatial distribution
B. Global Fishing Watch (GFW 2025) - GFW aggregates data from Automatic Identification Systems (AIS) transceivers on fishing vessels around the world and therefore provides data on individual fishing vessel movements around New Zealand

The spatial distribution of fishing effort for inshore trawlers and gillnetting vessels is available from government data (MPI 2023, 2024, 2025a,b). Additional information from GFW (2025) is available for about half of the medium-sized trawlers and a few small trawlers and gillnetting vessels (e.g. travel time from port to fishing area, number of days in port between fishing trips). In addition, we have direct observations at sea from three trawling vessels and observations of the amount of time they spend at sea vs. in port. The MPI and GFW data on length of each fishing trip (in hours), length of each trawl (in km), number of days in port between fishing trips and total number of trawls were used as the starting point for calibrating trawler movements and distributions. The following parameters were iteratively adjusted until the spatial distribution of fishing effort approximated the MPI, GFW and field data: Maximum travel time between leaving port and the first trawl of the day, travel time between subsequent trawl events, minimum and maximum trawl start depth and the probability of each subsequent trawl being within the same fishing area or switching to a new fishing area. To determine the match between the fishing effort data and trawler behavior in the model, heatmaps of model trawl start coordinates were created in R (R Core Team), using the kde.points function in the GISTools package to make kernel density estimates for simulated and observed positions of trawl starts. Sums of squared differences (SSD) were used to determine the per 1.63 x 1.63km grid-cell fit between modelled and the raw fishing effort data provided by MPI. A low SSD indicates a good fit with government data.

#### ***Patterns:***

*Trawl and Gillnet Location and Effort*

Heat maps of trawl and gillnet fishing effort data, comparing government estimates of fishing effort to model-generated data are shown in Figure C.1. The best fitting model was found by exploring a range of input values for maximum travel time between leaving port and the first trawl of the day (2-5 hrs for small and 2-7 for medium trawlers), travel time between subsequent trawl events (1-4 hrs for small, 2-5 for medium trawlers), finding the combination of inputs that resulted in the smallest SSD between government estimates of trawl fishing effort within 100m water depth and model outputs (Figure C.2). The travel time ranges explored were bounded by data on the behavior of real trawlers (GFW 2025). For example, maximum travel time from port to the first fishing location of 10 minutes or 10 hours would have been unrealistic compared to the behavior of real fishing vessels (GFW 2025). The initial calibration was in steps of 1 hour. Once getting close to the minimum SSD steps of 30 minutes were used. Gillnets were placed in model space based directly on the fishing effort data provided by government.

*Individual Trawler Behavior*

Figure C.3 compares the trawl tracks of a small trawler from GFW (2025; white lines) and the model (yellow lines) from the same port on Banks Peninsula and shows similar trawl lengths and preferred locations.

*Dolphins*

Data on the offshore range of Hector's and Maui dolphins from population surveys show that most sightings are in waters less than 100m deep with a small proportion (<5%) of sightings in deeper water (e.g. Slooten et al. 2006b; MacKenzie and Clement 2014, 2016, 2019). In the shallower waters off the east coast of the South Island of New Zealand, 41% of the population is found within 4 nautical miles (nmi) offshore, 46% between 4 and 12nmi offshore and 11% between 12nmi and the 100m depth contour, with some 2% of the population in waters deeper than 100m. Off the west coast of the South Island, water depth increases more rapidly with distance from shore. Here 76% of the population is found within 4nmi offshore, 22% between 4 and 12nmi offshore and 2% between 12nmi and the 100m depth contour, and no dolphins have been sighted in waters deeper than 100m.

*Dolphin Habitat Utilization (Wandering)*

We calibrated the model to ensure that dolphin movements and spatial distribution matched field data. To ensure that the dolphins responded realistically to water depth, the model was calibrated by generating heatmaps for a range of values of depth bias (k 10, 15, 20, 25, 30) and turning angle (wiggle 10, 15, 20, 25, 30 degrees) exploring all the permutations and combinations of k and wiggle. Each time a dolphin selects a new wandering heading, it uses its current heading plus a random normal value with mean zero and a Standard Deviation known as “wiggle” within Netlogo.

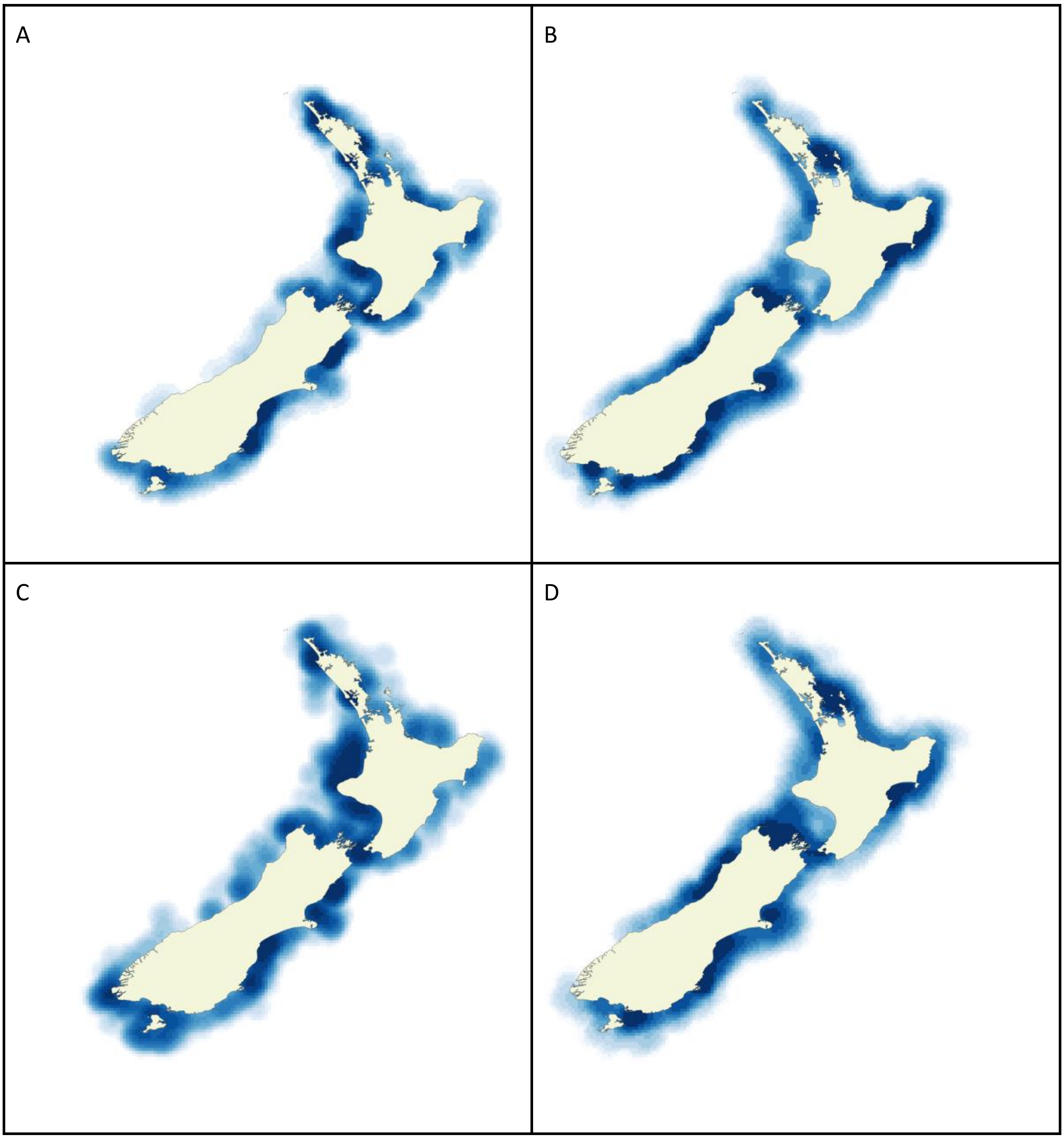

**Figure C.1** Heatmaps of inshore fishing effort showing government data on gillnets (A) and trawling (B), compared to fishing effort in the model for gillnets (C) and trawling (D).

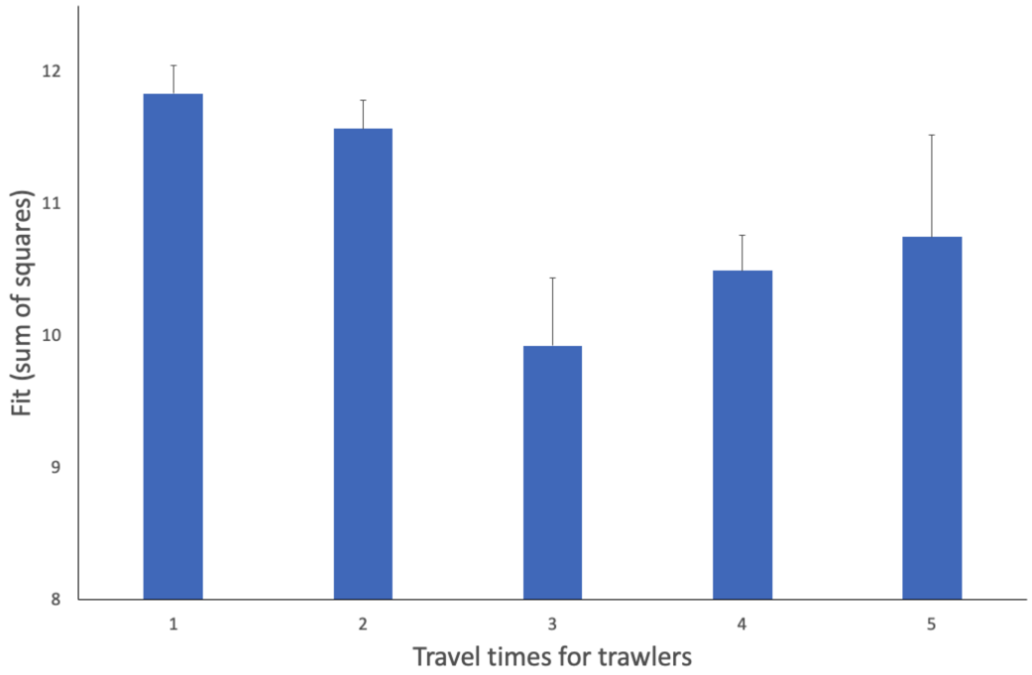

**Figure C.2** Match of spatial distribution of trawl fishing effort between our model and the government data (MPI 2023, 2024) depending on trawler travel times. After testing a wide range of travel times, the best fitting model (model 3, with lowest SSD) resulted from small trawlers initially travelling up to 3.5 hours between leaving port and their first trawl and then up to 2 hours between one trawl and the next, with medium trawlers travelling up to 6 hours initially and then up to 4.5 hours between trawls. In this same sequence, the other four models had the following travel times: Model 1 (3, 4, 2, 2); Model 2 (2, 1, 2, 3); Model 4 (4, 3, 5, 4); Model 5 (5, 3, 6, 5).

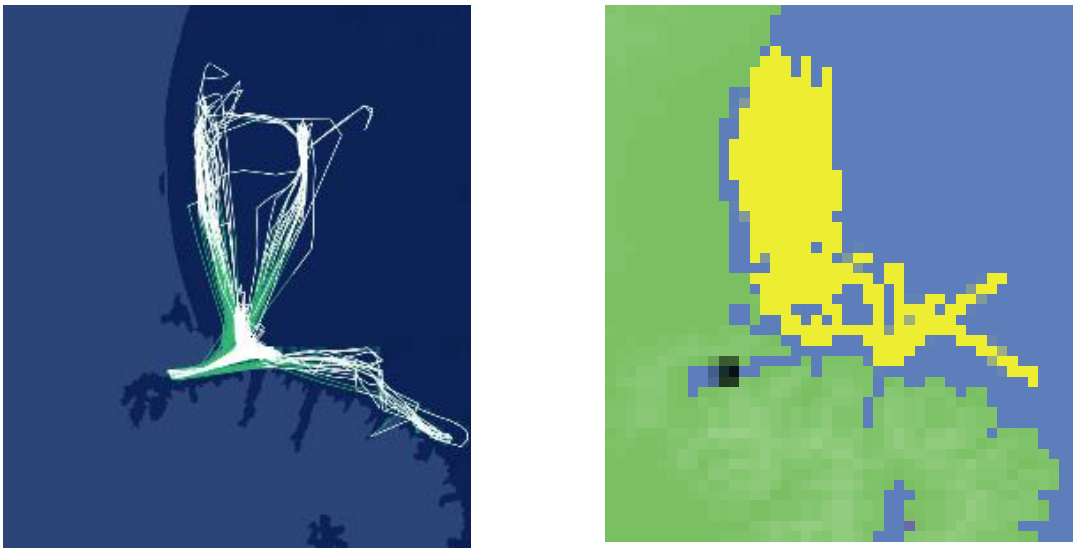

**Figure C.3** Individual Trawler Trawl Patterns: Data from Global Fishing Watch (2023) for a randomly selected trawler (left) compared to model data (right).

This initial, unbiased heading is then modified, based on the depth bias parameter (k). The strength of the dolphins' response to water depth is controlled by a logistic curve, with k determining the steepness of the curve. At high values of k, dolphins only respond when they are close to the maximum water depth of 100m and then respond strongly by changing their direction away from deep water. At low values of k, dolphins respond more gradually, turning away from deeper water when still in relatively shallow water and showing a relatively weaker response in terms of turning angle.

Results from the calibration runs were compared with dolphin density estimates from the three survey strata (0-4, 4-12 and 12-20nmi offshore) for the east coast of the South Island, the area with the most intensive population survey effort (MacKenzie and Clement 2014, 2016, 2019; Dawson et al. 2004; Rayment et al. 2009). The best model fit resulted from k = 15 and wiggle = 20 (Figure C.4). As expected, with a relatively gradual response to water depth dolphins spent too much time inshore (stippled bars in Figure C.4). With a very steep response to water depth, dolphin distribution was too 'flat', with too many dolphins offshore and too few inshore (striped bars in Figure C.4). Intermediate values for the depth response (k = 15) and turning angle (wiggle = 20) resulted in a very close fit between the survey data (black bars in Figure C.4) and dolphin distribution within the model (green bars in Figure C.4).

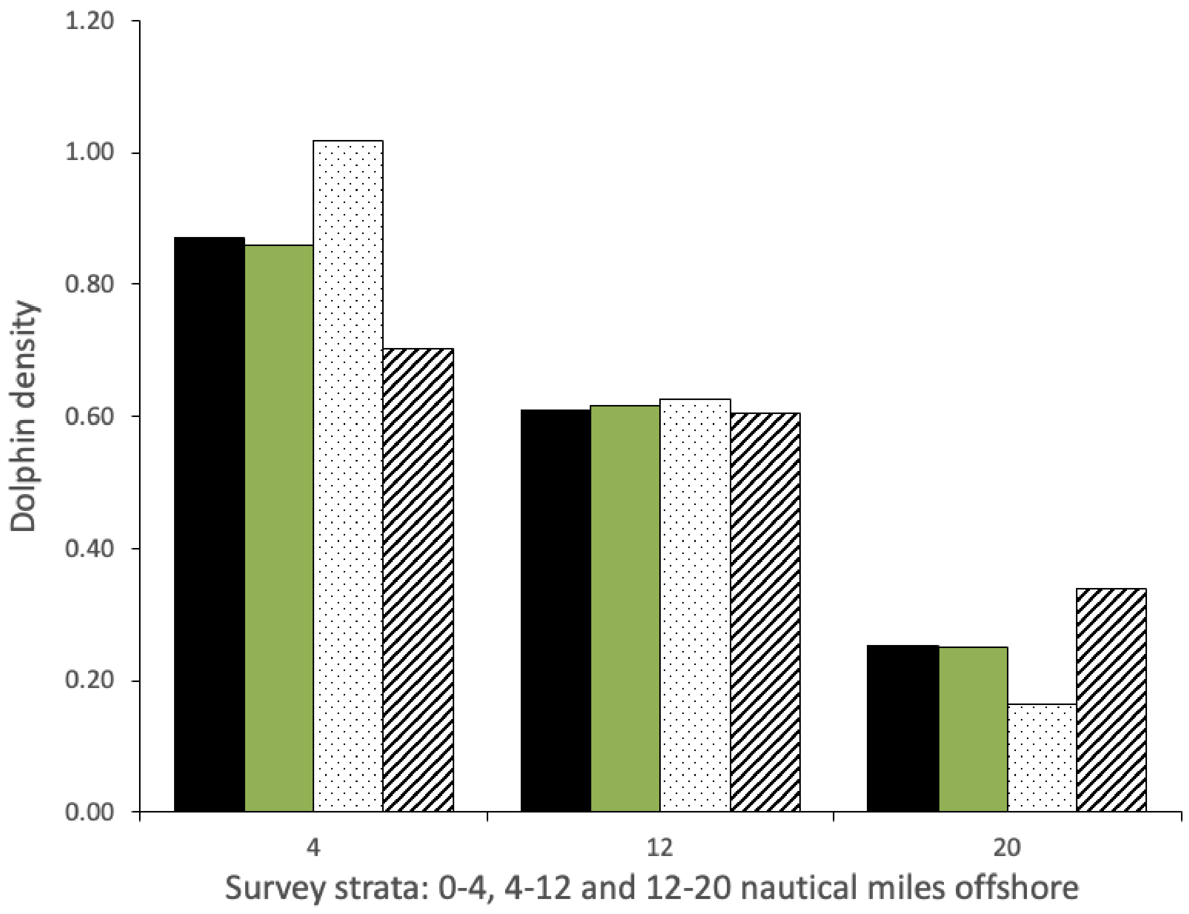

**Figure C.4** Dolphin densities (number of dolphins / km$^2$) in the three strata from the population surveys (black), for the best fitting model (green, k = 15, wiggle angle = 20) and for two other models for comparison (stippled: k = 10, wiggle = 10; striped:  k = 20, wiggle = 20).

*Dolphin Group Size*

Dolphins move in groups and the number of individual dolphins in each group fluctuates as dolphin groups split and merge. The probability of a group splitting or merging with another group depends on group size. The following parameters were iteratively modified to achieve the best fit between dolphin group size in the model and in field data: Minimum splitting probability (for a group of 2 dolphins) and shape factor (L) determining the shape of the curve relating minimum and maximum splitting probability (for groups of > 70 dolphins). The same process was followed for the probability of two groups merging, with very small groups having the highest probability of merging and the probability declining to zero for groups of > 50 individuals. Calibrations comparing Hector's dolphin group size in the model with field data consisted of systematically modifying the minimum merging probability (range 0.3-0.7), the shape factor for merging (L 0.1-0.5), the minimum splitting probability (range 0.1-0.5) and the shape factor for splitting (L 0.1-0.8). Figure C.5 shows the best fitting model for Hector's dolphin group size not following trawlers and examples of poor fitting models. For more information, please see ODD (Appendix B). Splitting and merging probabilities were calibrated separately for the Maui dolphin sub-species which has much lower overall group sizes in field data (Figure C.6).

Trawlers frequently attract more than one Hector's dolphin group, resulting in much larger numbers of dolphins following trawlers than the group sizes in Figure C.5. Calibrations comparing the number of Hector's dolphins following trawlers in the model with field data consisted of systematically modifying the minimum (range 1-4 hours) and maximum time (range 3-6 hours) dolphins spent following trawlers, and the minimum (2-8) and maximum time (6-12) not responding to trawlers after a period of following a trawler.

In Figure C.7 the best fitting model resulted from dolphins spending 3-5 hours (min-max) following a trawler (flocking) followed by 6-10 hours not following **trawlers** (wandering). The two poor-fitting models had the following inputs: 4-6, 4-8 (stippled bars) and 2-4, 4-8 (striped bars) and resulted in the number of dolphins following trawlers being too low or too high compared to the field data. The best fitting model was the best compromise, with the lowest discrepancy for both the smallest and largest group sizes. In the absence of data on the number of Maui dolphins seen following trawlers at any one time, we have assumed that they respond in a similar manner to trawlers.

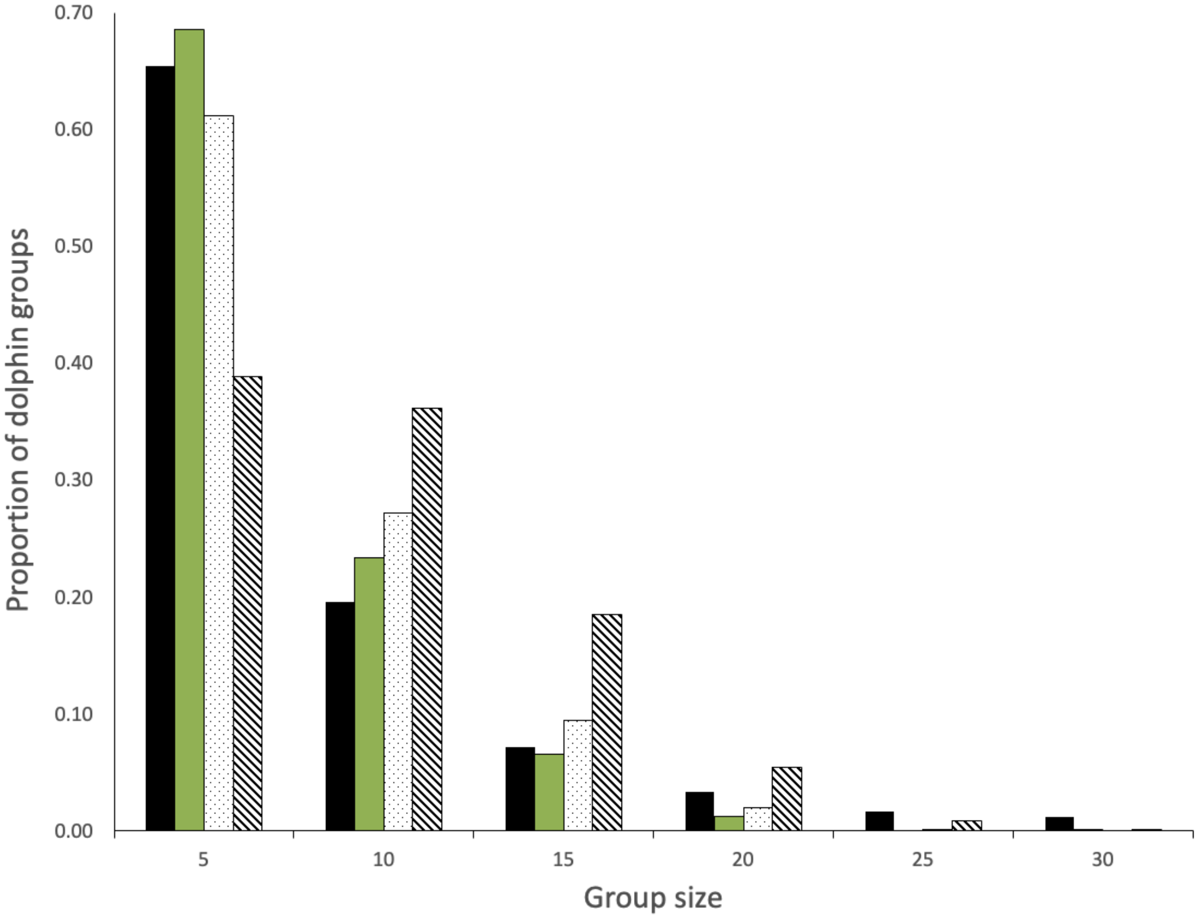

**Figure C.5** Observed group size of Hector's dolphins (field data in black), group size in the best fitting model (green) and in two poorly fitting models. Splitting probabilities for the smallest group (5 dolphins) were 0.2, 0.3 and 0.1 for the best fitting model and the two poor fitting models, respectively. The L parameter for splitting was 0.1, 0.7 and 0.7 in the same order (green, stippled, striped). Merging probabilities were 0.4, 0.4 and 0.4, with L of 0.1, 0.4 and 0.5 respectively.

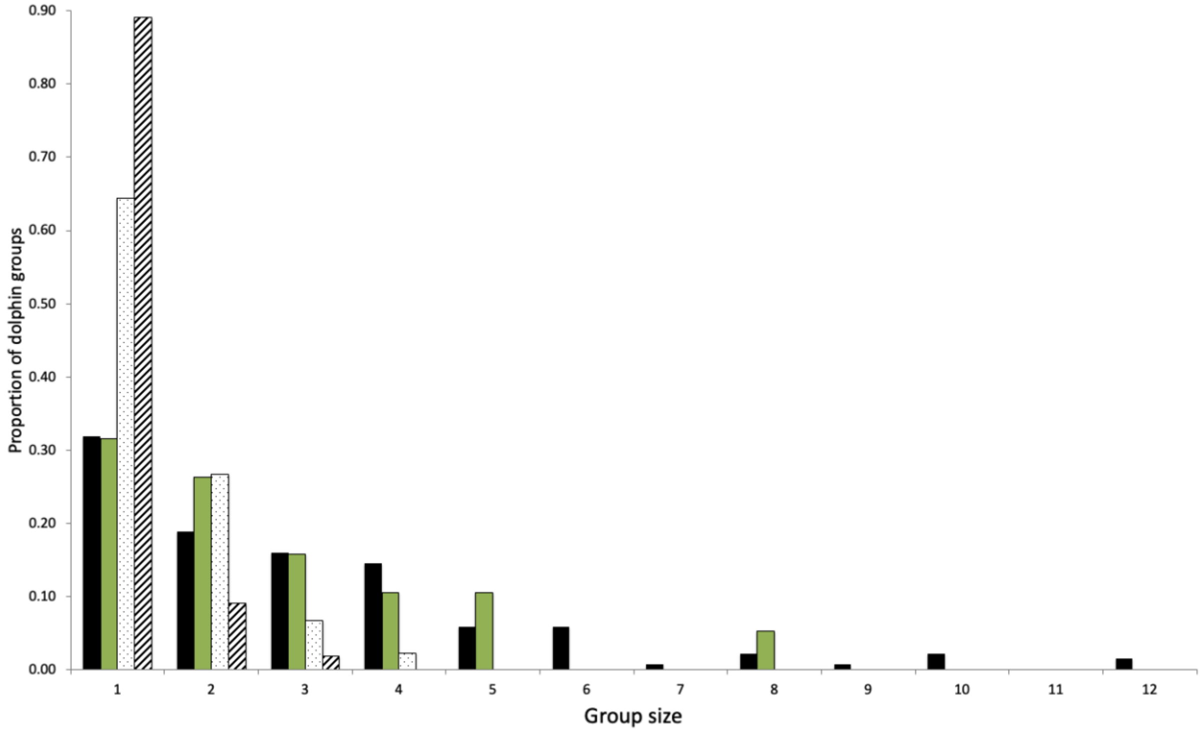

**Figure C.6** Observed group size of Maui dolphins in the field (black) compared to group size in the best fitting model (green) and two poor fitting models (stippled and striped). Splitting probabilities for the smallest group (2 dolphins) were 0.0001, 0.1 and 0.1 for the best fitting model and the two poor fitting models, respectively. The L parameter for splitting was 0.2, 0.5 and 0.5 in the same order (green, stippled, striped). Merging probabilities were 0.8, 0.4 and 0.6, with L of 0.1, 0.3 and 0.3 respectively.

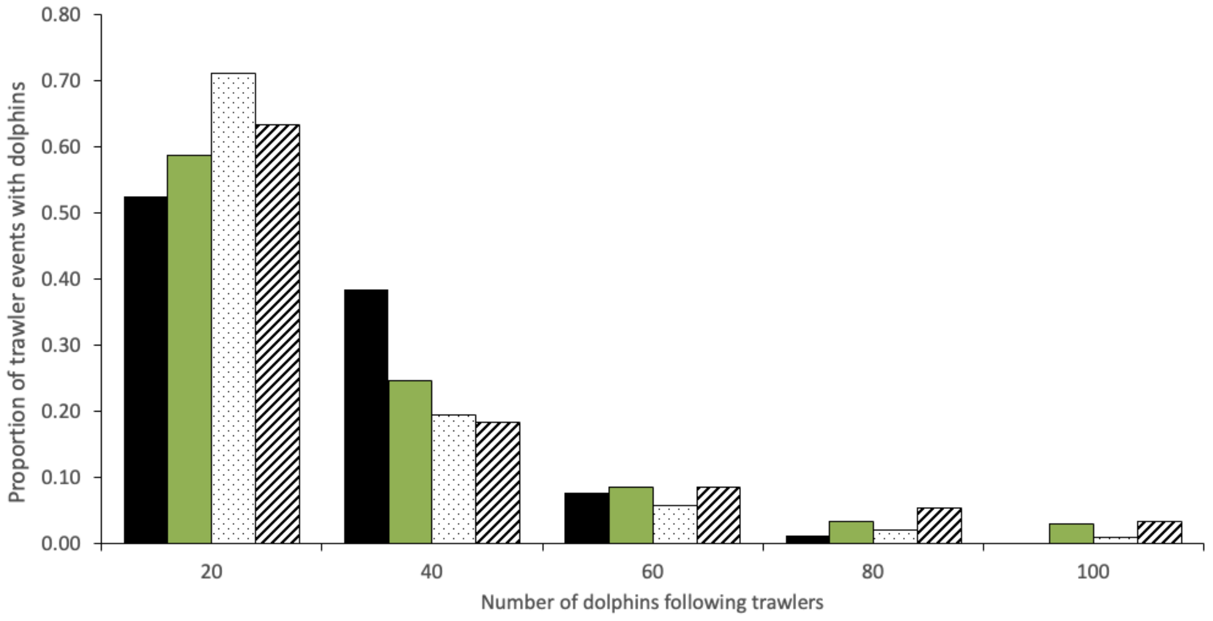

**Figure C.7** Number of Hector's dolphins following trawlers in field observations (black), best fitting model (green) and two poor fitting models (stippled and striped).

### References:

Bennington S. 2025. Investigating the recovery potential of Hector's dolphin *(Cephalorhynchus hectori).* PhD thesis, University of Otago, New Zealand.

Cervin L, Harkonen T, Harding KC. 2020. Multiple stressors and data deficient populations; a comparative life-history approach sheds new light on the extinction risk of the highly vulnerable Baltic harbour porpoises *(Phocoena phocoena).* Environment International 144: 106076.

Constantine R, Steel D, Carroll E, Hamner RM, Hansen C, Hickman G, Hillock K, Ogle M, Tukua P, Baker CS. 2021. Estimating the abundance and effective population size of Māui dolphins *(Cephalorhynchus hectori maui)* in 2020-2021 using microsatellite genotypes, with retrospective matching to 2001. Report to Department of Conservation, Auckland, New Zealand.

Cooke JG, Constantine R, Hamner RM, Steel D, Baker CS. 2019. Population dynamic modeling of the Māui dolphin based on genotype capture-recapture with projections involving bycatch and disease risk. Report to Ministry for Primary Industries, Wellington, New Zealand.

Daly E, White M. 2021. Bottom trawling noise: Are fishing vessels polluting to deeper acoustic habitats? Marine Pollution Bulletin 162: 111877.

Dawson SM, Lusseau D. 2006. Pseudoreplication problems in studies of dolphin and porpoise reactions to pingers. Marine Mammal Science 21: 175-176.

Dawson SM, Slooten E. 1993. Conservation of Hector's dolphins: The case and process which led to establishment of the Banks Peninsula Marine Mammal Sanctuary. Aquatic Conservation: Marine and Freshwater Ecosystems 3: 207-221.

Dawson S, Slooten E, Dufresne S, Wade P, Clement D. 2004. Small-boat surveys for coastal dolphins: Line-transect surveys for Hector's dolphins *(Cephalorhynchus hectori)*. Fishery Bulletin 102: 441-451.

Fletcher D, Dawson S, Slooten E. 2002. Designing a mark-recapture study to allow for local emigration. Journal of Agricultural, Biological and Environmental Statistics 7(4): 1-8.

Gende S, Hendrix AN, Schmidt J. 2018. Somewhere between acceptable and sustainable: When do impacts to resources become too large in protected areas? Biological Conservation 223: 138-146.

GFW. 2025. Global Fishing Watch. Online database of fishing activity using AIS, satellite imagery and AI: https://globalfishingwatch.org/

Gormley AM. 2009. Population modelling of Hector's dolphins. PhD thesis, University of Otago, Dunedin, New Zealand.

Gormley AM, Slooten E, Dawson SM, Barker RJ, Rayment W, du Fresne S, Bräger S. 2012. First evidence that marine protected areas can work for marine mammals. Journal of Applied Ecology 49:474-480.

Grimm V, Berger U, Bastiansen F, Eliassen S, Ginot V, Giske J, Goss-Custard J, Grand T, Heinz S, Huse G, Huth A, Jepsen JU, Jørgensen C, Mooij WM, Müller B, Pe'er G,

Piou C, Railsback SF, Robbins AM, Robbins MM, Rossmanith E, Rüger N, Strand E, Souissi S, Stillman RA, Vabø R, Visser U, de Angelis DL. 2006. A standard protocol for describing individual-based and agent-based models. Ecological Modelling 198: 115–126.

Grimm V, Berger U, de Angelis DL, Polhill JG, Giske J, Railsback SF. 2010. The ODD protocol: a review and first update. Ecological modelling, 221: 2760-2768.

Grimm V, Railsback, SF 2012. Pattern-oriented modelling: a ‘multi-scope’ for predictive systems ecology. Philosophical Transactions of the Royal Society B: Biological Sciences 367(1586): 298-310.

Grimm V, Railsback SF, Vincenot CE, Berger U, Gallagher C, de Angelis DL, Ayllón, D et al. 2020. The ODD protocol for describing agent-based and other simulation models: A second update to improve clarity, replication, and structural realism. Journal of Artificial Societies and Social Simulation 23: 7.

Grimm V, Revilla E, Berger U, Jeltsch F, Mooij WM, Railsback SF, Thulke H-H, Weiner J, Wiegand T, DeAngelis DL. 2005. Pattern-oriented modeling of agent-based complex systems: lessons from ecology. Science 310: 987–991.

Harvey M. 2021. Ecology and conservation of Hector’s dolphins in Porpoise Bay, Southland. MSc thesis, University of Otago, New Zealand.

IWC 2023. International Whaling Commission, Report of the 2023 Scientific Committee Meeting. 2023 Scientific Committee Report

Kastelein RA, Helder-Hoek L, van de Voorde S. 2017. Hearing thresholds of a male and a female harbor porpoise *(Phocoena phocoena).* Journal of the Acoustical Society of America 142: 1006-1010.

Kramer-Schadt S, Revilla E, Wiegand T, Grimm V. 2007. Patterns for parameters in simulation models. Ecological Modelling 204: 553-556.

Lusseau D, Kindt-Larsen L, van Beest FM. 2023. Emergent interactions in the management of multiple threats to the conservation of harbour porpoises. Science of the Total Environment 855: 158936.

Lynch JF, Newhall AE. 2017. Shallow-water acoustics. Chapter 7 In *Applied Underwater Acoustics*; Neighbors, T.H., Bradley, D., Eds.; Elsevier: Amsterdam, The Netherlands, pp. 403–467.

MacKenzie DL, Clement DM. 2014. Abundance and distribution of South Island east coast Hector’s dolphin. New Zealand Aquatic Environment and Biodiversity Report 123, Report to Ministry for Primary Industries, Wellington, New Zealand.

MacKenzie DL, Clement DM. 2016. Abundance and distribution of South Island west coast Hector’s dolphin. New Zealand Aquatic Environment and Biodiversity Report 168, Report to Ministry for Primary Industries, Wellington, New Zealand.

MacKenzie DL, Clement DM. 2019. Abundance and distribution of Hector’s dolphin on South Coast South Island. New Zealand Aquatic Environment and Biodiversity Report 236, Report to Ministry for Primary Industries, Wellington, New Zealand.

MPI 2023 and 2024. Ministry for Primary Industries provided data on fishing effort and protected area boundaries to Elisabeth Slooten. Government fisheries data are also available publicly, in a summarised form, at: Dragonfly.co.nz

MPI 2025a. Ministry for Primary Industries: Online data on bycatch of seabirds and marine mammals. seabirds-and-protected-marine-species-caught-by-commercial-fishers

MPI 2025b. Ministry for Primary Industries: South Island Hector’s dolphin bycatch reduction plan. Annual Report 2023/24.

Nabe-Nielsen J, Sibly RM, Tougaard J, Teilmann J, Sveegaard S. 2014. Effects of noise and by-catch on a Danish harbour porpoise population. Ecological Modelling 272: 242-251.

NIWA 2016. New Zealand Regional Bathymetry. National Institute of Water and Atmospheric Research. https://niwa.co.nz/our-science/oceans/bathymetry/

Peck SL. 2004. Simulation as experiment: a philosophical reassessment for biological modeling. Trends in Ecology and Evolution 19: 530–534.

Railsback SF, Grimm V. 2019. Agent-based and individual-based modelling. Second Edition, Princeton University Press.

Rayment W, Dawson SM, Slooten E, Bräger S, DuFresne S, Webster T. 2009. Kernel density estimates of alongshore home range of Hector's dolphins *(Cephalorhynchus hectori)* at Banks Peninsula. Marine Mammal Science 25: 537-556.

Richardson WJ, Greene CR, Malme CI, Thomson DH. 2013. Marine Mammals and Noise. Academic Press.

Roberts JO, Webber DN, Roe WD, Edwards CTT, Doonan IJ. 2019. Spatial risk assessment of threats to Hector's and Māui dolphins. New Zealand Aquatic Environment and Biodiversity Report 214, Ministry for Primary Industries, Wellington, New Zealand. https://www.fisheries.govt.nz/dmsdocument/35007

Rossmanith E, Blaum N, Grimm V, Jeltsch F. 2007. Pattern-oriented modelling for estimating unknown pre-breeding survival rates: The case of the Lesser Spotted Woodpecker *(Picoides minor).* Biological Conservation 135: 571–580.

Slooten E. 1991. Age, growth and reproduction in Hector's dolphins. Canadian Journal of Zoology 69: 1689-1700.

Slooten E. 1994. Behavior of Hector's dolphin: Classifying behavior by sequence analysis. Journal of Mammalogy 75: 956-964.

Slooten E. 2013. Effectiveness of area-based management in reducing bycatch of the New Zealand dolphin. Endangered Species Research 20: 121-130.

Slooten E, Dawson SM, Rayment WJ, Childerhouse SJ. 2006a. A new abundance estimate for Maui's dolphin: What does it mean for managing this critically endangered species? Biological Conservation 128: 576-581.

Slooten E, Dawson SM, Whitehead H. 1993. Associations among photographically identified Hector's dolphins. Canadian Journal of Zoology 71: 2311-2318.

Slooten E, Fletcher D, Taylor BL. 2000. Accounting for uncertainty in risk assessment: Case study of Hector's dolphin mortality due to gillnet entanglement. Conservation Biology 14: 1264-1270.

Slooten E, Lad F. 1991. Population biology and conservation of Hector's dolphin. Canadian Journal of Zoology 69: 1701-1707.

Slooten E, Rayment WJ, Dawson SM. 2006b. Offshore distribution of Hector’s dolphins at Banks Peninsula: Is the Banks Peninsula Marine Mammal Sanctuary large enough? New Zealand Journal of Marine and Freshwater Research 40: 333-343.

Stone G, Kraus S, Hutt A, Martin S, Yoshinaga A. 1997. Reducing by-catch: Can acoustic pingers keep Hector’s dolphins out of fishing nets? MTS Journal 31: 3-7.

TMP 2019. Threat Management Plan for Hector’s and Māui dolphins. 2019. Fisheries New Zealand and Department of Conservation. June 2019.

Turek J, Slooten E, Dawson S, Rayment W, Turek D. 2013. Distribution and abundance of Hector’s dolphins off Otago, New Zealand. New Zealand Journal of Marine and Freshwater Research 47: 181-191.

van Beest FM, Kindt-Larsen L, Bastardie F, Bartolino V, Nabe-Nielsen J. 2017. Predicting the population-level impact of mitigating harbor porpoise bycatch with pingers and time-area fishing closures. Ecosphere 8: e01785

Wickman L. 2024. Evaluating the long-term impacts of area-based protection on Hector’s dolphins at Banks Peninsula, New Zealand. PhD thesis, University of Otago, New Zealand.

Wiegand T, Jeltsch F, Hanski I, Grimm V. 2003. Using pattern-oriented modelling for revealing hidden information: a key for reconciling ecological theory and application. Oikos 100: 209–222.

Williams, H. 2022. Abundance and distribution of Hector's dolphins off the coast of Dunedin, New Zealand, and overlap with commercial fishing. MSc thesis, University of Otago, New Zealand. https://ourarchive.otago.ac.nz/handle/10523/13624
